# Compromised retinoic acid receptor beta (RARβ) accelerates the onset of motor, cellular and molecular abnormalities in mouse model of Huntington’s disease

**DOI:** 10.1101/2024.03.23.586058

**Authors:** Nicolas Zinter, Tao Ye, Valerie Fraulob, Damien Plassard, Wojciech Krezel

## Abstract

The mechanisms underlying detrimental effects of mutant huntingtin on striatal dysfunction in Huntington’s disease (HD) are not well understood. Although retinoic acid receptor beta (RARβ) emerged recently as one of the top regulators of transcriptionally downregulated genes in the striatum of HD patients and mouse models of HD its involvement in disease progression remains elusive. We report that genetically compromised RARβ signaling accelerates onset of motor abnormalities in R6/1 mouse model of HD. Transcriptional profiling revealed that downregulation of RARβ expression in *Rarβ*^*+/-*^; R6/1 mice also accelerates transcriptional signature of disease progression by emergence of upregulated cluster of genes related to cell-cycle, stem cell maintenance and telencephalon development with concomitant downregulation of striatal cell-identity genes. The reactivation of proliferative activity demonstrated in the neurogenic niche and development-related transcriptional programs in the striatum prompt an attempt of lineage infidelity in HD striatum which may lead in consequence to disease-driving energy crisis as suggested by concomitant downregulation of transcripts essential for oxidative phosphorylation, a well-accepted correlate of HD physiopathology, and a metabolic change required for maintenance of proliferative activity and differentiation but not compatible with high energetic demand of differentiated and active neurons.

**Highlights:**

- Compromising RARβ expression in R6/1 mouse model of HD accelerates onset of HD-like motor abnormalities
- Compromised RARβ signaling contributes to the progression of disease-related transcriptional changes in R6/1 mice
- RARβ supports cell-identity maintenance in HD mouse model

## INTRODUCTION

Huntington’s disease (HD) is an autosomal dominant neurodegenerative disease, characterized by progressive motor and cognitive symptoms. The age of onset of motor symptoms is highly variable, but starts, on average, in early 40s and progresses until the death of patients, which proceeds about 10-20 years after (Ross et al., 2014). Impairment of the striatum, and in particular the selective degeneration of medium spiny neurons (MSNs) is the hallmark of this disease (Vonsattel and DiFiglia, 1998). Although these changes are key components of the HD physiopathology, a number of additional correlates were also associated with disease progression, including increased proliferation and neurogenesis in the neurogenic niche of the ventricular and subventricular zone (SVZ), but meaning of this correlate remains not well understood (Curtis et al., 2005, 2003; Gil-Mohapel et al., 2011).

HD is caused by abnormal CAG repeat expansion in the first exon of the Huntingtin (HTT) gene. The length of the resulting polyglutamine repeats correlates negatively with the age of onset. This knowledge allowed generation of multiple genetic models including R6/1 transgenic mice carrying the first exon of human mutant Huntingtin (*mHTT*) (Mangiarini et al., 1996), but also an allelic series of mouse models carrying knock-in (KI) of human *mHTT* exon 1 with different lengths of CAG (Q) repeats ranging from Q20 corresponding to a non-toxic form of CAG stretch up to Q175 corresponding to the toxic form (Pouladi et al., 2013). Like in human, these mouse models develop progressive motor and cognitive dysfunction with different age of onset, which in mice depends on the model and the length of CAG repeats (Langfelder et al., 2016; Menalled et al., 2012; Nithianantharajah et al., 2008; Smith et al., 2014). Transcriptional changes appeared to be much more sensitive markers of disease progression then behavior as they were observed at presymptomaptic age in Q140 and Q175 *mHTT* KI mice but were also present in Q80, Q92, Q111 models which do not display any major phenotype (Langfelder et al., 2016). Such molecular signatures of disease progression may also provide insight into its physiopathology which is complex. Specifically, the latter study allowed to define 37 modules (M1-37) of striatal genes which were co-expressed at 2, 6, and 10 months of age in allelic series of Q20-Q175 KI models. Additional modules were identified in non-striatal compartments and in some cases were indicative of the whole-brain/body HD-specific changes. Although some of these modules co-varied in age- and CAG repeat-dependent manner the determination of the most relevant one(s) for understanding mechanisms of the disease onset remains a challenge.

Retinoic acid receptor beta (RARβ), a ligand-dependent transcription factor mediating retinoic acid (the major active form of vitamin A) signaling emerged recently as potential atop regulator of gene networks affected in HD (Ciancia et al., 2022; Lee et al., 2020; Niewiadomska-Cimicka et al., 2017). Thus, reduced RARβ expression observed consistently in the striatum of HD patients and animal model (Hodges et al., 2006; Luthi-Carter et al., 2000; Niewiadomska-Cimicka et al., 2017) could contribute to the physiopathology of HD. In line with this possibility, RARβ loss of function in *Rarβ*^*-/-*^ mice led to HD-like motor deficits most probably through mitochondrial dependent manner (Ciancia et al., 2022). RARβ deficiency may directly affect striatonigral and striatopallidal projection MSNs (called hereafter snMSNs and spMSNs respectively), the main cell types expressing RARβ in mouse and human striatum (Ciancia et al., 2022; Lee et al., 2020). The spMSNs function and survival appear particularly dependent on RARβ activity. Accordingly, loss of dopamine D2 receptor (*Drd2*) expressing spMSN was observed in *Rarβ*^*-/-*^ striatum whereas somatic inactivation of *Rarβ* in postmitotic *Drd2*+ neurons or in the whole striatum recapitulated both behavioral and spMSN deficits displayed by *Rarβ*^*-/-*^ mice (Ciancia et al., 2022).

Despite these correlative observations the role of compromised RARβ signaling in HD is not clear. By investigating animal behavior, we report here that genetically compromised RARβ signaling accelerated onset of HD-like motor abnormalities in R6/1 transgenic mice. Transcriptional profiling revealed that compromising RARβ signaling in R6/1 mice accelerated transcriptional signature of disease progression by emergence of upregulated cluster of genes related to cell-cycle, stem cell maintenance and telencephalon development with concomitant downregulation of striatal cell-identity genes. The reactivation of cell division and developmental program further documented by increased proliferative activity in the neurogenic niche in the striatum could prompt an attempt of lineage infidelity which may lead to disease-driving energy crisis in striatal neurons as suggested by transcriptional changes coherent with reduction of oxidative phosphorylation, required for maintenance of proliferative and developmental activities but unsustainable for high energy demand of differentiated and active neurons.

## MATERIAL AND METHODS

### Animals

Hemizygous R6/1 mice (Mangiarini et al., 1996) were maintained on CBA genetic background for more than 10 generations whereas mice carrying a null mutation for *Rarβ* (*Rarβ*^*-/-*^) were generated from heterozygous crosses as previously described (Ghyselinck et al., 1997) and were maintained on mixed genetic background of about 50% C57BL6J and 50% 129svEmx for more than 10 generations. Both lines were crossed to generate the first generation of R6/1^Tg/0^; *Rarβ*^*+/-*^ mice and corresponding littermate controls which were used for behavioral tests and subsequent molecular analyses. For all experiments, mice were housed (3–4/cage) with water and food available *ad libitum* and 12 h/12 h light/dark cycle with the beginning of the light phase at 7 am. Mice were sacrificed 2 days after the last behavioral experiment for further cellular and molecular analyses. All experiments were carried out in accordance with the Directive of the European Parliament: 2010/63/EU and in compliance with the guidelines of CNRS and the French Agricultural and Forestry Ministry (decree 87848).

### Behavioral phenotyping

All behavioral analyses were carried out using 8-week-old males. Behavioral test battery consisted of following tasks which were performed in the order of appearance with one test on each consecutive day.

#### Open field

Mice were tested in automated open field arenas (Bioseb, France). Briefly, mice were individually placed in the periphery of the open field and allowed to explore the apparatus freely for 30 min. The distance travelled, the number of rears and time spent in the central and peripheral regions were recorded over the test session. The test was performed in a room homogeneously illuminated at 150 Lux.

#### Rotarod

Accelerated (4–40 rpm in 5 min) rotarod (Bioseb, France) was used for all experiments. Each test consisted of three trials separated by 15 min recovery intervals. The latency time to fall from the rotarod was recorded. Unless mice fell from the rotating rod, to avoid rotarod passive turning habituation, the cut off time was settled after 3rd passive rotation. Before the first trial mice were allowed to stay on the rotarod for about 30 s of habituation period before starting the acceleration phase.

#### Spontaneous locomotion in actimetric cages

Spontaneous locomotor activity was measured in actimetric cages (Immetronic, Pessac, France) during 32h (11 am until 7 pm the next day). Briefly, each mouse was placed in an individual cage equipped with infra-red photo beam cells that measure horizontal movements. During the experiment mice had free access to food and water.

### In situ hybridization and cell counts

14μm cryosections were thawed and dried for 30 min at RT, post-fixed with 4% paraformaldehyde (PFA) dissolved in phosphate-buffered saline (PBS), acetylated with 0.1 M triethanolamine (TEA), rinsed with standard saline citrate (SSC), dehydrated by an increasing ethanol gradient, air-dried and hybridized at 65°C overnight with digoxigenin-labelled Drd2 RNA probe (1681 bp fragment of 5’ region of Drd2 cDNA). After hybridization the sections were rinsed, blocked and incubated for 1 h at RT with an anti-digoxigenin, alkaline phosphatase-conjugated antibody (Roche). The sections were subsequently rinsed and incubated in the dark with a NBT/BCIP solution (Nitroblue tetrazolium chloride/ 5-bromo-4chloro-3-indyl-phosphate; Roche) until the staining appeared (overnight or up to 48 h). If the staining reaction was longer than overnight, the NBT/BCIP solution was changed after 24 h. Imaging was performed using Nanozoomer digital scanner (Hamamatsu Photonics, France) and cells were counted manually by two independent experimenters using ImageJ software.

### Western blotting

Striatum samples from newly born mice (P0) and adults (20 weeks-old) were homogenized manually in RIPA lysis buffer (Tris HCl 50 mM, NaCl 30 mM, EDTA 2 mM, Triton 0.01X) supplemented with a protease inhibitor cocktail (#11873580001, Roche Diagnostic). Samples were then sonicated (2 ×3 s) and left 30 min on ice. Following a 30 minutes centrifugation, protein concentration was quantified using Bradford assay (#5000006, Biorad) and 10 μg of protein extracts were separated by SDS–PAGE, and electro-transferred (400 mA during 1 h) onto 0.2μm nitrocellulose membrane (#A29612452, Amersham). Immunodetection was carried out using rabbit anti-RARβ, 1:1000 (Rochette-Egly et al., 1992); mouse anti-GAPDH, 1:5000 (CAB932Hu22, USCN) and horseradish peroxidase (HRP)-conjugated secondary antibodies (Goat anti-mouse, 1:5000, G-21040, Invitrogen; Goat anti-rabbit, 1:5000, #111 035 44, Jackson Immunoresearch). Enzymatic detection was carried out using Super Signal West Femto Maximum Sensitivity Substrate (#34095, Thermoscientific). GAPDH was used as a loading control when quantifying RARβ expression.

### Immunohistochemistry

Immunohistochemistry were performed on 14μm brain cryosections that were thawed and dried for 10 min at room temperature. Sections were then fixed with paraformaldehyde 4% for 15 min and washed 3 times in PBS. Antigen retrieval was then carried out for 20 minutes at 90°C in 0.01M citrate buffer pH6. Slides were cooled down for 1h at room temperature under agitation and washed 3 times in PBS during 10 min and permeabilized and blocked in a solution of PBS + Triton 0.1% + FCS 10% for 1h at room temperature. Incubation with primary antibodies (see below) was done overnight at +4°C, slices were washed 3 times 10 minutes in PBS and incubated with secondary antibodies (see below) for 1h at room temperature.

| Primary antibodies |  |  |  |
| --- | --- | --- | --- |
| Name | Species | Reference | Dilution |
| Ki67 | Rabbit | Synaptic System, #HS-398103 | 1:500 |
| PH3 | Mouse IgG1 | Proteintech, #66863 | 1:500 |
| GFAP | Mouse IgG2a | Synaptic System, #173011 | 1:500 |
| NESTIN | Chicken | Avès-labs, #NSE-0020 | 1:5000 |
| SOX2 | Rabbit | Proteintech, #11064-1-AP | 1:500 |

| Secondary antibodies |  |  |
| --- | --- | --- |
| Name | Reference | Dilution |
| Donkey@Rabbit | Invitrogen, #31572 | 1:500 |
| Donkey@Mouse<sub>IgG1</sub> | Invitrogen, #21121 | 1:500 |
| Goat@Mouse<sub>IgG2a</sub> | Invitrogen, #21131 | 1:500 |
| Goat@Chicken | Invitrogen, #21449 | 1:500 |

### Transcriptomic analysis

Nac samples were punched on thick cryosections (300μm) and RNA was extracted using RNeasy MicroPlus kit from Qiagen following the manufacturer protocol. RNA-Seq libraries were generated from 200 ng of total RNA using TruSeq Stranded mRNA Library Prep Kit and TruSeq RNA Single Indexes kits A and B (Illumina, San Diego, CA), according to the manufacturer’s instructions. Briefly, following purification with poly-T oligo attached magnetic beads, the mRNA was fragmented using divalent cations at 94oC for 2 minutes. The cleaved RNA fragments were copied into first strand cDNA using reverse transcriptase and random primers. Strand specificity was achieved by replacing dTTP with dUTP during second strand cDNA synthesis using DNA Polymerase I and RNase H. Following addition of a single ‘A’ base and subsequent ligation of the adapter on double stranded cDNA fragments, the products were purified and enriched with PCR (30 sec at 98oC; [10 sec at 98oC, 30 sec at 60oC, 30 sec at 72oC] x 12 cycles; 5 min at 72oC) to create the cDNA library. Surplus PCR primers were further removed by purification using AMPure XP beads (Beckman-Coulter, Villepinte, France) and the final cDNA libraries were checked for quality and quantified using capillary electrophoresis. Libraries were then single-read sequenced with a length of 50 bp on an Illumina HiSeq 4000 sequencer. Image analysis and base calling were carried out using RTA v.2.7.7 and bcl2fastq v.2.17.1.14. Reads were mapped onto the mm10 assembly of Mus musculus genome using TopHat v.2.0.14 (Kim et al., 2013) and Bowtie 2 v.2.1.0 (Langmead and Salzberg, 2012). Gene expression was quantified from uniquely aligned reads using HTSeq-count v.0.6.1 (Anders et al., 2015) with annotations from Ensembl release 90 and intersection-nonempty mode. Only non-ambiguously assigned reads have been retained for further analyses. Comparisons of interest have been performed using R 3.5.1 with DESeq2 version 1.22.1 (Love et al., 2014). More precisely, read counts were normalized from the estimated size factors using the median-of-ratios method and, for the comparisons between each group, a Wald test was used to estimate the P-values.

To check the existence of an interaction between the mHTT and the *Rarβ*^*+/-*^ as an aggravating factor, a Likelihood-Ratio Test was used with DESeq2. Unwanted variation was identified with sva v.3.30.1 (Leek, 2014) and taken into account in the statistical model. P-values were then adjusted for multiple testing with the IHW method (Ignatiadis et al., 2016).

The enrichment analyses of deregulated genes from present study within all the gene modules identified by Langfelder et al. (Langfelder et al., 2016) was performed using the ORA methods of EGSEA R package (1.26.0; (Alhamdoosh et al., 2017)). For this purpose, the differentially expressed genes (p<0.05) were converted to NCBI Entrez gene id with Ensembl biomart release 90. A custom gene set collection was created from the modules downloaded from (Langfelder et al., 2016) using buildCustomIdx of EGSEA. Over-representation Analysis of each deregulated gene list was performed using egsea.ora function.

The enrichment analyses were additionally conducted using GSEA method implemented in GSEA software version 4.3.2 (Mootha et al., 2003; Subramanian et al., 2005). The identical custom gene set collection and the ontology gene sets of MSigDB(citing, maybe other gene sets as well?) were employed. Genes were preranked using the formula sign(logFC) * -log10(pval) following the DESeq2 analysis.

## RESULTS

### Compromised RARβ exacerbates motor deficits in mouse model of HD

To gain insight into the role of RARβ in HD, we investigated whether compromising *Rarβ* expression in HD mouse model accelerates the onset of HD-like symptoms. To this end, we generated R6/1 mice carrying only one copy of functional *Rarβ* and monitored their motor behaviors at pre-symptomatic age as readout of disease symptoms. We found that 8-week-old R6/1^Tg/0^; *Rarβ*^*+/-*^ mice displayed significant ∼50% deficit in motor coordination in the rotarod which was not observed in mice carrying only a transgene of human mHTT exon 1 (R6/1^Tg/0^; *Rarβ*^*+/+*^) or were heterozygous for *Rarβ* null mutation (R6/1^0/0^; *Rarβ*^*+/-*^) (Fig. 1A). Accordingly, two-way ANOVA analyses revealed a significant interaction between mHTT transgene and *Rarβ* heterozygosity (F(1,50)=4.2, p<0.05) for this performance whereas post-hoc analyses supported significant reduction of latency to fall from accelerated rotarod in R6/1^Tg/0^; *Rarβ*^*+/-*^ mice as compared to wild-type (R6/1^0/0^; *Rarβ*^*+/+*^) and R6/1^0/0^; *Rarβ*^*+/-*^ or R6/1^Tg/0^; *Rarβ*^*+/+*^ littermates. Such locomotor coordination deficit did not affect overall novelty-induced locomotion and rearing behaviors in the open field which were not altered in any group of mice and attained on average respectively ∼100 meters of distance and ∼170 rears during 30 minutes of the test (Fig. S1A, S1B). These measures were neither biased by abnormal anxiety states as the percentage of time spent in the center of the open arena during the first 5 minutes of the test was on average ∼7% and not different between any of the tested groups (Fig. S1C). Importantly, R6/1^Tg/0^; *Rarβ*^*+/+*^ mice carrying mHTT transgene alone displayed spontaneous hyperlocomotion in actimetric cages during an active, dark phase of the light/dark cycle and the *Rarβ* heterozygosity exacerbated such hyperactivity in R6/1^Tg/0^; *Rarβ*^*+/-*^ mice remaining without any effect on locomotion in transgene negative R6/1^0/0^; *Rarβ*^*+/-*^ mice. This is supported by significant *R6/1 x Rarβ* interaction (F(1,52)=14.2, p<0.001, two-way ANOVA) and respective post-hoc analyses (Fig. 1B, C).

**Fig. 1.**
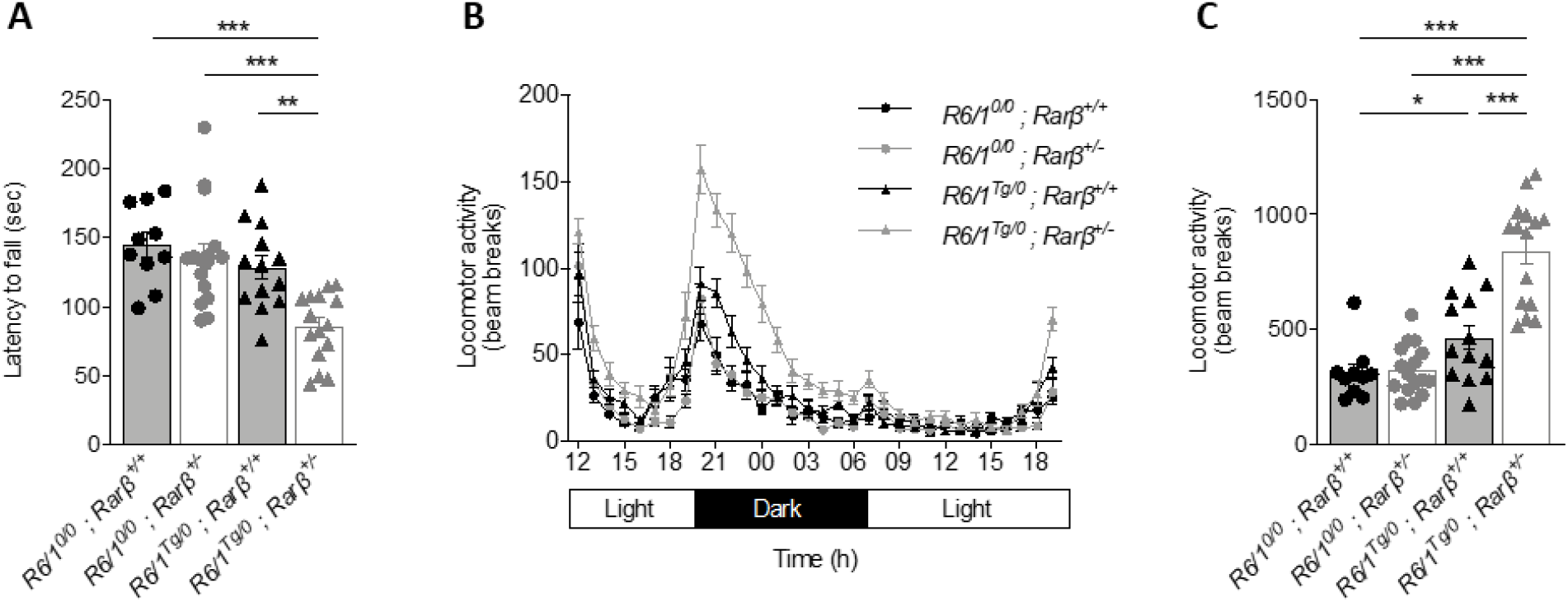
Compromised RARβ signaling in HD mouse model exacerbates motor deficits linked to HD. (A) Motor coordination evaluated in accelerated rotarod as mean latency to fall from the rotating cylinder. (B) Spontaneous locomotor activity measured in actimetric cages over 32 h. (C) Spontaneous locomotor activity scores for the dark phase of the light/dark cycle in actimetric cages measured between 7pm and 7am. statistical differences were calculated using Bonferroni multiple comparisons as post-hoc follow-up of ANOVA analyses : *, p < 0.05; * *, p < 0.01; * **, p< 0.001. Error bars represent SEM.

### Compromised RARβ signaling in R6/1 mice does not induce loss of spMSNs

Since studies of *Rarβ* knockout mice revealed role of RARβ in post-natal protection of about 20% of spMSNs from cell death (Ciancia et al., 2022), we hypothesized that compromised RARβ expression in HD striatum may contribute and possibly explain a particularly high sensitivity of *Drd2*+ spMSNs to die in HD striatum. Thus, exacerbating RARβ deficiency in HD model by inactivation of *Rarβ* allele in R6/1^Tg/0^; *Rarβ*^*+/-*^ mice, we could expect to observe *Drd2*+ spMSN cell death. To address this possibility, we first investigated whether RARβ heterozygosity indeed reduces RARβ expression. Our western blot analyses confirmed that in the caudate putamen (CPu) of R6/1^0/0^; *Rarβ*^*+/-*^ mice RARβ level was reduced to 30 ± 11% of that observed in wild-type R6/1^0/0^; *Rarβ*^*+/+*^ controls. Surprisingly RARβ protein was similarly reduced in transgenic, R6/1^Tg/0^; *Rarβ*^*+/+*^ mice, and did not progress any further in compound R6/1^Tg/0^; *Rarβ*^*+/-*^ mice remaining at 38 ± 14% and 38 ± 10% of wild-type expression level respectively (Fig. 2A). This is supported by the main effect of *Rarβ* and R6/1 genotypes (F (1, 8) = 21,7, p<0.01 and F(1, 8) = 13,3, p<0.01 respectively) but also significant interaction between the two genotypes (F(1, 8) = 21,76, p<0.01). Such effect was cumulative in the nucleus accumbens shell (NAc) as supported by main effect of each genotype (F(1, 8) = 22,8 for R6/1, p<0.01 and F (1, 8) = 57,7, p<0.001 for RARβ) and significant reduction of RARβ in R6/1^Tg/0^; *Rarβ*^*+/-*^ as compared to R6/1^0/0^; *Rarβ*^*+/-*^ or R6/1^Tg/0^; *Rarβ*^*+/+*^ striatum (Fig. 2B).

**Fig. 2.**
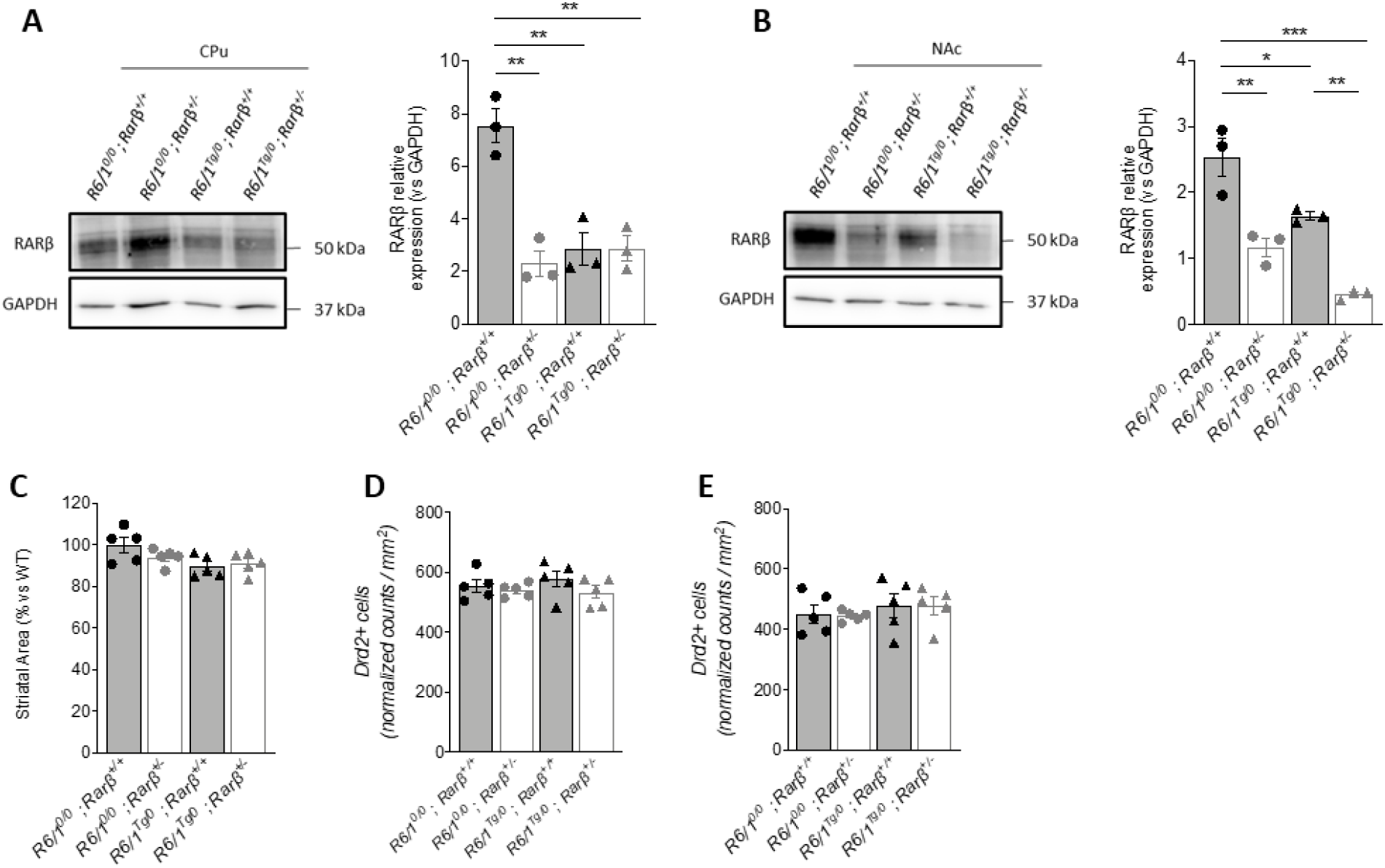
Compromised RARβ signaling in R6/1 mice does not induce loss of spMSNs. (A) Western blot detection (left panel) and quantification (right panel) of RARβ in CPu and (B) in NAc. (C) Measure of striatal area for each genotype was reported as percent of mean striatal area in the wild type (R6/1^0/0^; *Rarβ*^*+/+*^) group. (D) Numbers of *Drd2*+ MSNs cell were normalised with respect to striatal area changes for each genotype and reported in CPu and (E) in NAc. Statistical differences were calculated using Bonferroni multiple comparisons as post-hoc follow-up of ANOVA analyses : *, p < 0.05; * *, p < 0.01; * **, p < 0.001. Error bars represent SEM.

We have then quantified the number of *Drd2*+ spMSNs using *in situ* hybridization to detect corresponding cells in the striatum. In contrast to expected reduced density of *Drd2*+ cells per square millimeter, we have observed its increase both in CPu and NAc, which was associated exclusively with the presence of mHTT transgene as supported by main effect of R6/1 genotype as the only significant difference in two-way ANOVA analyses (F (1, 16) = 10,62, p<0.01 for CPu and F (1, 16) = 7,7, p=0.01 for NAc) even though follow-up post-hoc analyses did not support specific difference between any individual group (Fig. S2A, S2B, S2C). Such result could reflect a decrease of striatal volume due to mHTT-induced atrophy reported previously in several HD mouse models (Gil-Mohapel et al., 2011). To consider this possibility, we have calculated changes in striatal surface at each bregma level used for analyses (Fig. 2C). Reduction of striatal size was evident from the main effect of mHTT transgene (F (1,16)= 5,71, p<0.05) supporting an atrophy induced by the presence of mHTT transgene, but it was not aggravated by monoallelic inactivation of *Rarβ*. Normalization of cell counts with respect to the surface change clearly supported the absence of significant change in the number of *Drd2*+ cells in both CPu and NAc in any of the genotypes as compared with wild-type group (Fig. 2D, 2E, S2C).

### Transcriptomic analyses reveal synergistic dysregulation of cell cycle and neurodevelopmental gene set associated with HD progression

In addition to hypothesis driven analyses of *Drd2*+ cell number, we also used transcriptomic profiling as an unbiased approach to determine the contribution of reduced RARβ expression in HD-related changes. We focused such analyses on NAc region in which RARβ downregulation was exacerbated in *R6/1*^*Tg/0*^; *Rarβ*^*+/-*^ mice. We found that, at 8 weeks of age, *R6/1*^*Tg/0*^; *Rarβ*^*+/+*^ mice carrying only mHTT transgene were in fact already severely affected with 358 down and 93 upregulated genes in comparison to wild-type littermates (at cutoff of p<0.05; Tab. S1). This and additional pool of genes was affected by further inactivation of one copy of *Rarβ* in *R6/1*^*Tg/0*^; *Rarβ*^*+/-*^ mice which displayed 402 down and 140 upregulated genes when compared to wild-type condition and 247 down- and 248 upregulated genes in comparison to *R6/1*^*Tg/0*^; *Rarβ*^*+/+*^ mice (Tab. S1). These RARβ-dependent changes in R6/1 background were not merely an additive effect of *Rarβ* heterozygosity as there were only 14 down and 13 upregulated genes in *R6/1*^*0/0*^; *Rarβ*^*+/-*^ as compared to wild-type *R6/1*^*0/0*^; *Rarβ*^*+/+*^ striatum (Tab. S1). Functional annotations of differentially expressed genes (DEGs) were made using gene set enrichment analyses (GSEA). As expected, mHTT alone in *R6/1*^*Tg/0*^; *Rarβ*^*+/+*^ mice affected some of the key biological functions associated with HD including synaptic transmission, monoamine and dopaminergic signaling, mitochondrial components and activity (Tab. 1 and Fig. S3). Although, many similar functions were also affected in compound *R6/1*^*Tg/0*^; *Rarβ*^*+/-*^ mice, the direction of changes was not always congruent. For example, gene sets associated with control of animal behavior, monoamine and catecholamine transport and signaling were similarly downregulated, however, mitochondrial complex and ribosome functions were upregulated in presence of mHTT alone in *R6/1*^*Tg/0*^; *Rarβ*^*+/+*^ but downregulated in presence of concomitant *Rarβ* deletion in *R6/1*^*Tg/0*^; *Rarβ*^*+/-*^ mice. Finally, reduced synaptic plasticity observed in *R6/1*^*Tg/0*^; *Rarβ*^*+/+*^ was not among the top affected functions in *R6/1*^*Tg/0*^; *Rarβ*^*+/-*^ and inversely, an upregulation of genes related to telencephalon development, cell-cycle transition or stem-cell maintenance, increased cilium, axoneme assembly, in latter condition were not observed in *R6/1*^*Tg/0*^; *Rarβ*^*+/+*^ mice (Tab. 1 and Fig. S3).

**Table 1.**
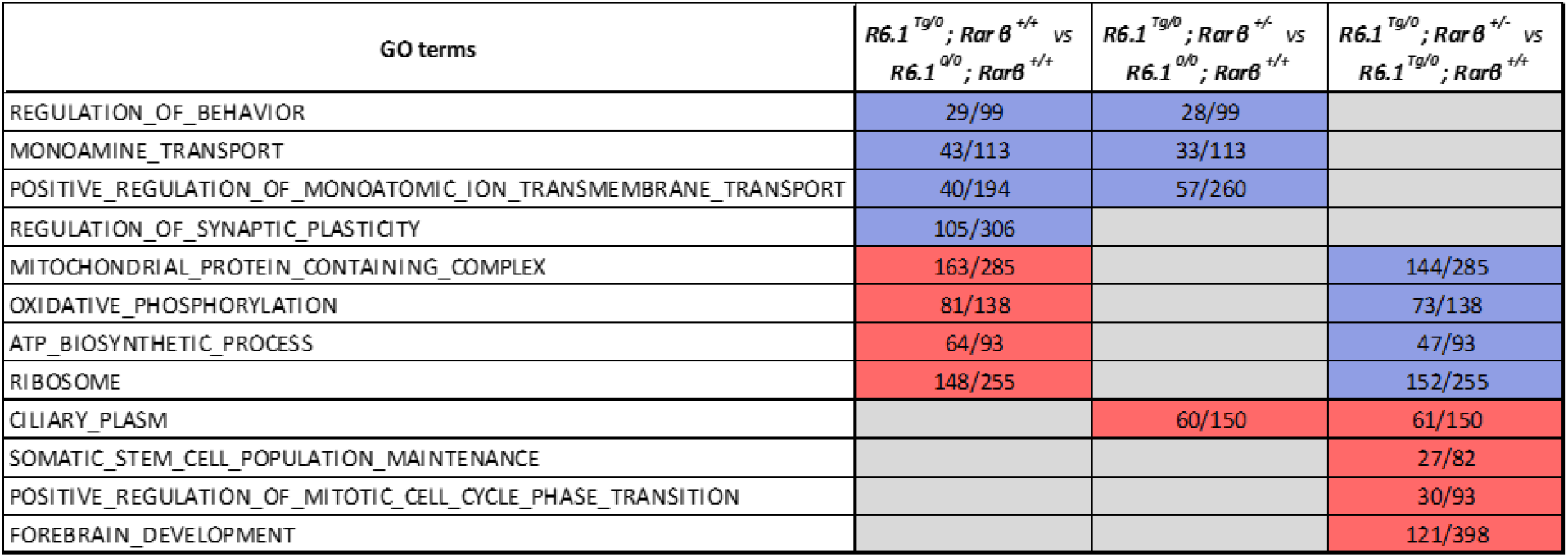
Functional annotations of DEGs in distinct comparisons. Examples of Gene Ontology Biological Processes or Cellular Components affected by mHTT and Rarb expression were drawn from the top-20 list of significant changes with FDR < 0.0001. Upregulation of gene expression was represented in red and downregulation in blue. Core esnrichment / enrichment score values were shown in corresponding cells.

To identify transcriptional changes associated with HD onset and/or progression, we have compared our data with Huntingtin CAG length- and age-dependent gene co-expression network identified by Langfelder et al. (Langfelder et al., 2016). Using pre-ranked enriched GSEA (EGSEA), as expected, we found that DEGs in *R6/1*^*Tg/0*^; *Rarβ*^*+/+*^ carrying mHTT alone were the strongest enriched in M2, the module (module’s designation according to Langfelder et al.) harboring downregulated striatal cell identity genes and identified as having the strongest association with CAG length and best preserved in human HD datasets (Fig. 3A). Significant enrichments were also observed for M11, the module with high numbers of glial genes, M43 and M9, associated with mitochondrial functions, M7 with cell death genes. Comparison of *R6/1*^*Tg/0*^; *Rarβ*^*+/-*^ and *R6/1*^*0/0*^; *Rarβ*^*+/+*^ wild-type mice revealed similar but enhanced profile of enrichment with emergence of genes belonging to M39 related to stress response and DNA damage repair (Fig. 3B). The genetic interaction between compromised *Rarβ* expression and mHTT transgene was, however, the best visible through comparison of DEGs in *R6/1*^*Tg/0*^; *Rarβ*^*+/-*^ and *R6/1*^*Tg/0*^; *Rarβ*^*+/+*^ mice (Fig. 3C-G). The major transcriptional changes associated with inactivation of one *Rarβ* allele in R6/1 mice were enriched in modules ordered according to its significance from 22 through 16, 5, 2, 30, 11, 24, 25, 9, 42 to 4 (Fig. 3C) out of which modules M2, 11, 25 and 9 were reported as highly correlated with disease progression by Langfelder et al. Importantly, DEGs from *R6/1*^*Tg/0*^; *Rarβ*^*+/-*^ vs *R6/1*^*Tg/0*^; *Rarβ*^*+/+*^ comparison displayed congruent changes with those observed in striatum of early symptomatic, 6-month-old Q175 mice, as illustrated by marked downregulation of mitochondrial genes from module M22, and upregulation of cell cycle and development-related genes in M16 and M20, as illustrated by EGSEA results (Fig. 3D, E). This suggests that *R6/1*^*Tg/0*^; *Rarβ*^*+/-*^ specific changes which were sometimes opposite to those observed in *R6/1*^*Tg/0*^; *Rarβ*^*+/+*^ littermates (mitochondrial functions and cell-cycle, developmental genes) are perhaps more relevant for disease progression. In particular, M16 and M20 modules emerged as synergistically dysregulated by mHTT transgene and *Rarβ* heterozygosity in Likelihood-ratio analyses (Fig. 3E, F).

**Fig. 3.**
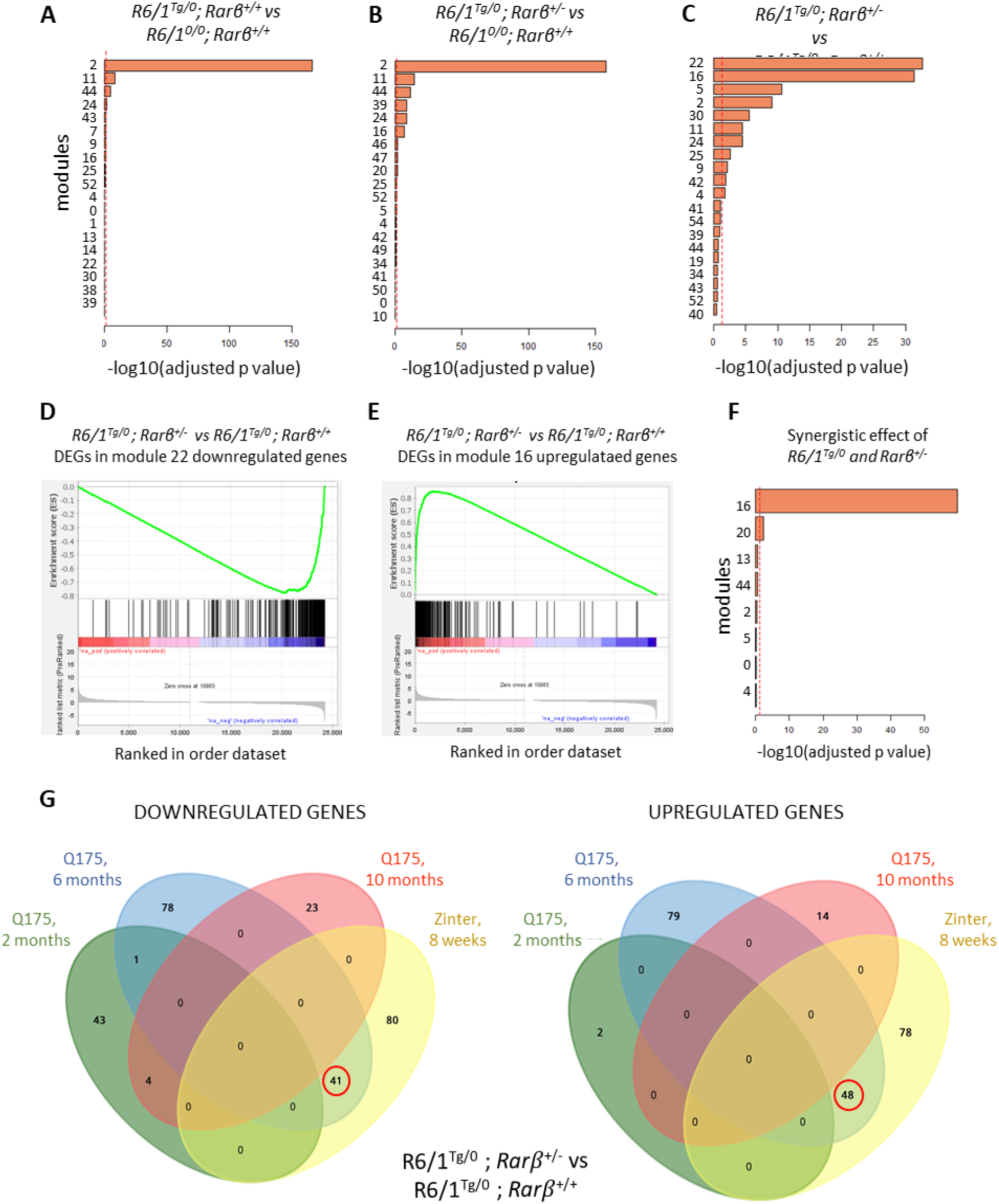
Comparison of DEGs associated with compromised RARβ signaling in R6/1 NAc. (A) DEGs from *R6/1*^*Tg/0*^; *Rarβ*^*+/-*^*vs R6/1*^*Tg/0*^; *Rarβ*^*+/+*^comparison and (B) *R6/1*^*0/0*^; *Rarβ*^*+/-*^ *vs R6/1*^*0/0*^; *Rarβ*^*+/+*^comparison, and (C) *R6/1*^*Tg/0*^; *Rarβ*^*+/-*^*vs R6/1*^*Tg/0*^; *Rarβ*^*+/+*^comparison were represented as enrichment in gene modules associated with HD progression determined by Langfelder et al. (Langfelder *et al.*, 2016). (D) GSEA for *R6/1*^*Tg/0*^; *Rarβ*^*+/-*^*vs R6/1*^*Tg/0*^; *Rarβ*^*+/+*^deregulated gene set as compared with downregulated genes in Q,.175 HD mouse model at 6 months of age and belonging to module 22. (E) GSEA for *R6/1*^*Tg/0*^ ; *Rarβ*^*+/-*^*vs R6/1*^*Tg/0*^; *Rarβ*^*+/+*^gene changes compared with upregulated genes in Q.175 HD mouse model at 6 months of age and belonging to module 16. (F) Gene expression changes synergistically affected by R6/1 and *Rarβ*^*+/-*^in *R6/1*^*Tg/0*^ ; *Rarβ*^*+/-*^ striatum were determined by Likelihood-Ratio test and analysed fee their enrichment in gene modules associated with HD progression. (G) Venny plot representing *R6/1*^*Tg/0*^ ; *Rarβ*^*+/-*^specific DEGs related to module M16 in comparison to M16 downregulated (left panel) and upregulated (right panel) genes In Q175 mice across different ages.

We next analyzed the age dependence of M16-related changes which we have observed in *R6/1*^*Tg/0*^; *Rarβ*^*+/-*^ striatum as compared to *R6/1*^*Tg/0*^; *Rarβ*^*+/+*^ littermates at 8 weeks by comparing them to age-dependent transcriptional deregulation in Q175 KI mouse model reported by Langfelder et al.. Specifically, we found that more than 50% of such changes were found in symptomatic, 6-month-old Q175 mice but were absent from pre-symptomatic, 2-month-old, or advanced stage 10-month-old Q175 mice (Fig. 3G).

### Compromised RARβ signaling drives reactivation of cell-cycle programs and lineage infidelity in HD mouse

As supported by comparisons of individual compound mutant mice but also Likelihood-Ratio Test of synergy between mHTT transgene and *Rarβ*^*+/-*^, an increase of genes associated with cell-cycle regulation was the most consistent change induced by *Rarβ* heterozygosity in R6/1 mice. Furthermore, the direction of changes of this cluster of genes correlated with those observed in DEGs of M16 and M20 modules by Langfelder and colleagues. We hypothesized that such changes in the NAc may reflect an overall reactivation of cell-cycle/division transcriptional programs, which should be best visible in the ventricular and subventricular zone (VZ and SVZ respectively, both referred as SVZ hereafter) (Fig. 4A), an adult neurogenic niche harboring actively dividing cell. Analyses of Ki67+ proliferating cells in the SVZ region revealed a significant effect of *Rarβ* heterozygosity (F(1,8)=74,4, p<0.001) which affected the number of proliferating cells in opposite direction depending on the presence of *mHTT* as supported by significant *R6/1 x Rarβ* interaction (F(1, 8) = 214, p<0.001) (Fig. 4B, 4C). Accordingly, whereas loss of one copy of *Rarβ* alone led to significant reduction of Ki67+ cells, their number was significantly increased in presence of *mHTT* transgene in *R6/1*^*Tg/0*^; *Rarβ*^*+/-*^ SVZ. Unexpectedly, *mHTT* transgene, alone, strongly reduced the number of Ki67+ cells in SVZ of *R6/1*^*Tg/0*^; *Rarβ* ^*+/+*^ mice. As revealed by analyses of phospho-histone 3 co-immunostaining (PH3+), the percentage of cells in S-phase was similar among all genotypes and ranged about 5.5% of Ki67+ cells with exception of *R6/1*^*Tg/0*^; *Rarβ*^*+/+*^ mice in which the few Ki67+ cells were also PH3-positive (Fig. 4D). We therefore asked whether changes in the number of proliferating cells may result from covariation of the reservoir of neuroblasts / neural stem cells (called hereafter collectively NSCs). Despite the almost complete loss of Ki67+ cells, the number of GFAP+; Sox2+; Nestin+ triple positive NSCs was not affected in *R6/1*^*Tg/0*^; *Rarβ*^*+/+*^ mice (Fig. 4E, 4F). However, a marked increase of NSCs correlated with higher cell proliferation index in *R6/1*^*Tg/0*^; *Rarβ*^*+/-*^ SVZ. This is supported by significant group effects for both *R6/1* (F (1,8) = 5.77, p<0.05) and *Rarβ* (F (1,8) = 8.80, p<0.05.; Fig. 4F) and suggests that increased proliferation rate observed in these mice results from normal activation of an increased pool of ependymal NSCs.

**Fig. 4.**
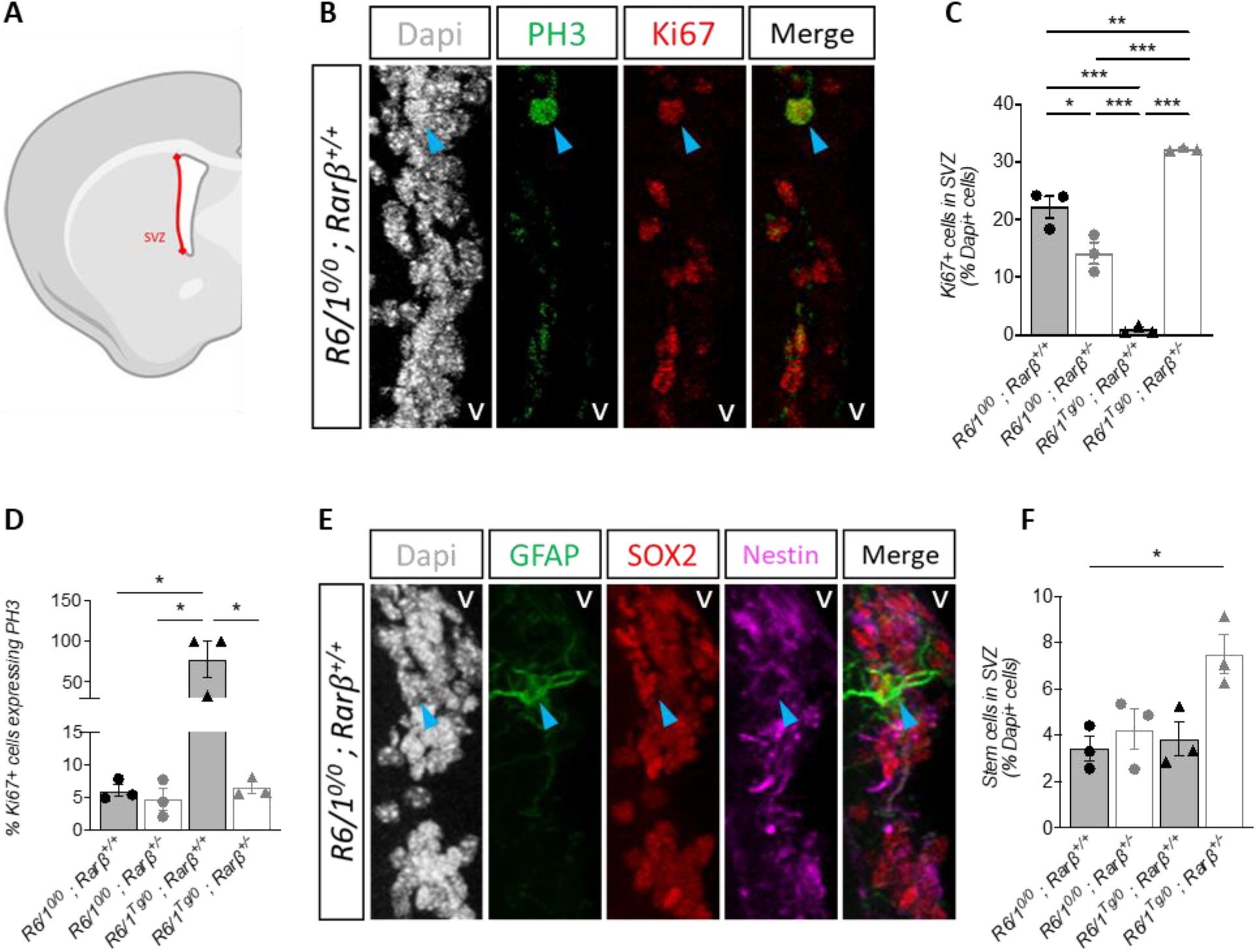
Compromised RARβ signaling drives reactivation of cell-cycle programs and lineage infidelity in HD mouse. (A) Cartoon indicating ventricular /subventricular zone (called throughout SVZ, red line) in a mouse coronal brain section. (B) Examples of immunofluorescence identifying proliferating Ki67+ and PH3+ cells in *R6/1*^*0/0*^; *Rarβ*^*+/+*^ SVZ. Blue arrows represent double positive cels. The letter V indicates position of the ventricle. (C) Quantification of Ki67+ cells (% of cells). (D) Quantification of PH3+ cells among Ki67+ cells in SVZ (% of cells). (E) Examples of immunofluorescence identifying GFAP+ ; S0X2+. Nestin+ neural stem cells (NSC) in *R6/1*^*0/0*^; *Rarβ*^*+/+*^SVZ. Blue arrows represent triple positive cells. The lener V indicates position of the ventricle. (F| NSC quantification (% of cells). Statistical differences were calculated using Bonferroni multiple comparisons as post-hoc fdow-up of ANOVA analyses : *, p < 0.05; * *, p< 1***;, p < 0.001. Error bars represent SEM.

## DISCUSSION

Reduced expression of *Rarβ* and dysregulation of a large number of its transcriptional target genes observed in the striatum of HD patients and animal models, suggested that *Rarβ* may play a critical role in progression of HD related transcriptional changes driving thereby disease progression (Lee et al., 2020; Luthi-Carter et al., 2000; Niewiadomska-Cimicka et al., 2011). Although this possibility is further supported by neuroprotective activity of RARβ in *Drd2* neuron functions and survival (Ciancia et al., 2022), the actual contribution of compromised RARβ signaling to HD physiopathology have been mostly correlative. By investigating animal behavior, we revealed that compromised RARβ signaling is sufficient to precipitate HD-like motor abnormalities in R6/1 mouse model of HD at 8 weeks of age, which corresponds to motor pre-symptomatic period in R6/1 model (Naver et al., 2003; Nithianantharajah et al., 2008). Accordingly, R6/1 mice carrying deletion of one copy of *Rarβ* displayed significant coordination deficits and spontaneous hyperlocomotion, whereas R6/1 littermates displayed only weak spontaneous hyperlocomotion at this age whereas *Rarβ*^*+/-*^ mice appeared haplosufficient in every aspect of analyzed behavior. Such precocious coordination deficits and exacerbation of hyperlocomotion in *R6/1*^*Tg/0*^; *Rarβ*^*+/-*^ indicate that RARβ acts as disease modifier preventing disease progression. The critical time window of RARβ protective activity is not clear, but may take place in early postnatal life or during adolescence, as at 8 weeks, mHTT transgene alone was sufficient to reduce *Rarβ* expression levels to those observed in *R6/1*^*Tg/0*^; *Rarβ*^*+/-*^ CPu, whereas in NAc, the R6/1 and *Rarβ* heterozygosity displayed only some additive effect in reducing *Rarβ* expression. Likewise, significant HD-like transcriptional changes were present already at 8 weeks of age in the striatum of *R6/1*^*Tg/0*^; *Rarβ*^*+/+*^ mice carrying R6/1 transgene alone.

A large array of transcriptional changes have been revealed in HD patients and animal models with the most dramatic ones observed in the striatum, the primary site of degeneration (Benn et al., 2005; Hodges et al., 2006, 2006; Lee et al., 2022). Although determination of meaningful changes still remains a challenge, it was much advanced by a large study of Huntingtin CAG length- and age-dependent gene co-expression networks (Langfelder et al., 2016). Among 37 expression modules (M1-37) of striatal genes defined by these authors, many co-varied in age- and CAG repeat-dependent manner. Through transcriptomic profiling of striatal tissue, we found that R6/1 transgenic mice display a significant HD-related profile of gene expression changes including downregulation of genes related to cell-identity, monoaminergic and synaptic signaling already at 8 weeks of age. These genes belonged to M2 module identified by Langfelder et al. as the strongest associated with CAG length and best preserved in human HD datasets. Compromising RARβ signaling in R6/1 mice advanced this profile. Of particular interest was emergence of upregulated cluster of genes related to cell-cycle, stem cell maintenance and telencephalon development. This change was functionally relevant as the number of proliferating cells in *R6/1*^*Tg/0*^; *Rarβ*^*+/-*^ SVZ was significantly increased reflecting an increased number of GFAP+; Sox2+; Nestin+ triple positive NSCs and neuroblasts in the ependymal and SVZ layer of these mice. Whereas such an increase was evident in the *R6/1*^*Tg/0*^; *Rarβ*^*+/-*^ adult neurogenic niche, where proliferation is naturally occurring, no evident proliferating cells were observed in other parts of the striatum including CPu but also NAc, the region used for transcriptional profiling. Several lines of evidence suggest however, that increased expression of those genes may be of direct relevance for the progression of HD. Accordingly, increased mitotic activity was reproducibly observed in HD post-mortem studies (Curtis et al., 2005, 2003; Pelegrí et al., 2008), although it was rarely studied and found only weakly increased or unchanged in mouse models of HD (Gil et al., 2005; Gil-Mohapel et al., 2011; Jin et al., 2005; Lazic et al., 2006, 2004). A change found more consistently between HD patients and mouse models, was upregulation of striatal genes related to cell division and brain development. Using Likelihood-ratio analyses, we found that *R6/1*^*Tg/0*^ synergized with *Rarβ*^*+/-*^ in upregulation of those genes as compared to mice carrying only mHTT transgene (Fig. 3F). Thus, these genes were synergistically enriched in module M16 and M20, the latter positively correlated with CAG length in Langfelder’s study (Langfelder et al., 2016). Interestingly, more than 50% of *R6/1*^*Tg/0*^; *Rarβ*^*+/-*^ DEGs enriched in module 16 were almost exclusively associated with the onset of symptoms in Q175 KI mice at 6 months of age as compared to pre-symptomatic 2-month old or advanced stage 10-month-old mice (Fig. 3G and (Langfelder et al., 2016)), suggesting potential importance of corresponding deregulations in the control of the onset of the disease. The reactivation of cell-cycle and developmental genes could possibly be considered as lineage infidelity and part of the putative regenerative process as in case of wound repair (Ge et al., 2017). However, even if such process is ineffective in HD, it does not come at no cost. Whereas neuronal functions critically depend on energy produced during oxidative phosphorylation (OXPHOS), this metabolic mode is not compatible with cell-cycle and developmental process which are supported by glycolytic cycle, much less efficient in producing energy (Gascón et al., 2016; Tsogtbaatar et al., 2020). Accordingly, concomitant to upregulation of genes related to cell cycle, stem cell maintenance and development *R6/1*^*Tg/0*^; *Rarβ*^*+/-*^ mice also displayed also strong downregulation of oxidative phosphorylation, ATP biosynthesis, but also ATP-dependent processes like ribosomal functions. We therefore propose that, the role of RARβ is to delay HD progression by preventing lineage infidelity attempt, associated upregulation of neuronal redifferentiation program and resulting downregulation of OXPHOS and ATP production leading in consequence to energy crisis unsustainable for neuronal functions and driving thereby onset of behavioral deficits.

*Drd2*+ spMSNs are the most prone to neurodegeneration and RARβ activity was shown previously to play a role in their protection against cell death otherwise observed in *Rarβ*^*-/-*^ mice but also in wild-type mice treated with 3NP, a mitochondrial toxin (Ciancia et al., 2022). Despite evident behavioral deficits in *R6/1*^*Tg/0*^; *Rarβ*^*+/-*^ mice and downregulation of a large set of mitochondrial genes, we did not observe decrease in the number of *Drd2*+ cells in the NAc or CPu of *R6/1*^*Tg/0*^; *Rarβ*^*+/-*^ animals. This data further supports that RARβ neuroprotective role goes beyond prevention of cell death, but is more essential for spMSN functions.

Whether neuroprotective activity of RARβ in R6/1 mice is entirely cell-autonomous, remains unclear. In fact, whereas an increase of the number of neuroblasts and/or neural stem cells was observed in ependymal and adjacent cell layers where RARβ was not shown to be expressed, the activation of the cell cycle, including increased maintenance of NSCs, may also involve extrinsic signals such as secreted factors.

Finally, this study shed light on previously not reported shut-down of proliferative activity in R6/1 mice at 8 weeks of age. The origin and meaning of such a block is not clear but may correspond to a transient posing at G0 or cell cycle block at G2 to M transition. Such reduced proliferation might be highly age-specific as previous studies of older R6/1 mice did not report such changes in SVZ region (Lazic et al., 2006, 2004) which also agrees with our unpublished data suggesting even slight increase of proliferation in 14-week-old R6/1 mice. Thus, to better understand the contribution of RARβ deficiency to HD progression there is a need for systematic analysis of RARβ functions during early postnatal life in physiological conditions and HD-mouse models.

## ACKNOWLEDGMENTS

We thank Mr. Valentin Fraxial for assistance in behavioral analyses, Dr. Gill Bates for sharing R6/1 mouse model of HD and Mouse Clinical Institute animal facility for excellent animal care. The project was supported by ERA-Net RAinRARE and institutional grant ANR-10-LABX-0030-INRT, a French State fund managed by the Agence Nationale de la Recherche under the frame program Investissements d’Avenir ANR-10-IDEX-0002-02. Sequencing was performed by the GenomEast platform, a member of the ‘France Génomique’ consortium (ANR-10-INBS-0009).

