## Supplemental data for "Compromised retinoic acid receptor beta (RARβ) accelerates the onset of motor, cellular and molecular abnormalities in mouse model of Huntington’s disease"

### *Zinter et al., 2024*

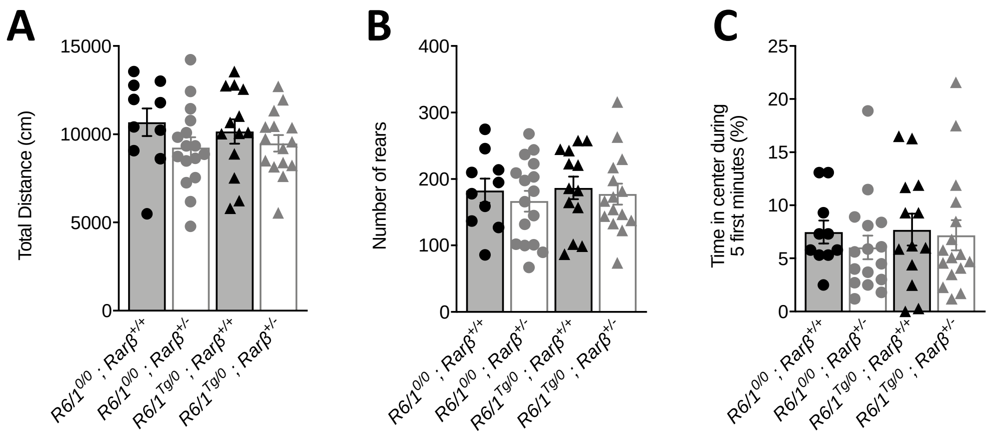


**Fig. S1. Compromised RARβ signaling in R6/1$^{Tg/0}$; *Rarβ*$^{+/-}$ R6/1 mice does affect novelty induced locomotion.** (A) Novelty induced locomotor activity in the open field evaluated by total distance covered during 30min of the test. (B) Quantification of the number of rears in open field during 30-minute test. (C) Evaluation of the anxiety induced by novelty in open field during the five first minutes. Error bars represent SEM.

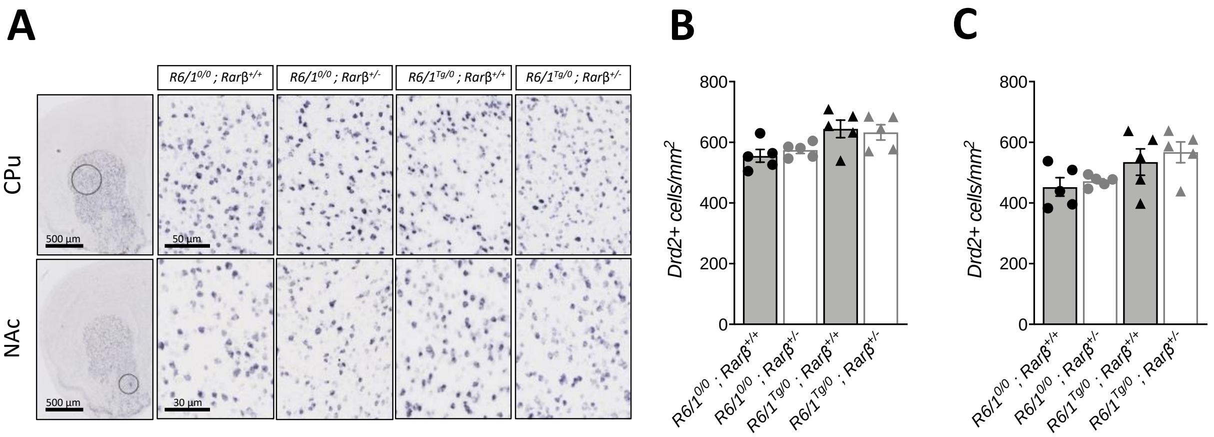


**Fig. S2. Analyses of Drd2+ neurons in the striatum.** (A) Examples of in situ hybridization identification of *Drd2*+ spMSNs in CPu and NAc. (B) *In situ* hybridization quantification of *Drd2*+ MSNs per square millimiter in CPu and (C) in NAc. Error bars represent SEM.

# A

Enrichment of downregulated genes in *R6/1^{Tg/0}; Rarβ^{+/+} vs R6/1^{0/0}; Rarβ^{+/+}* comparison

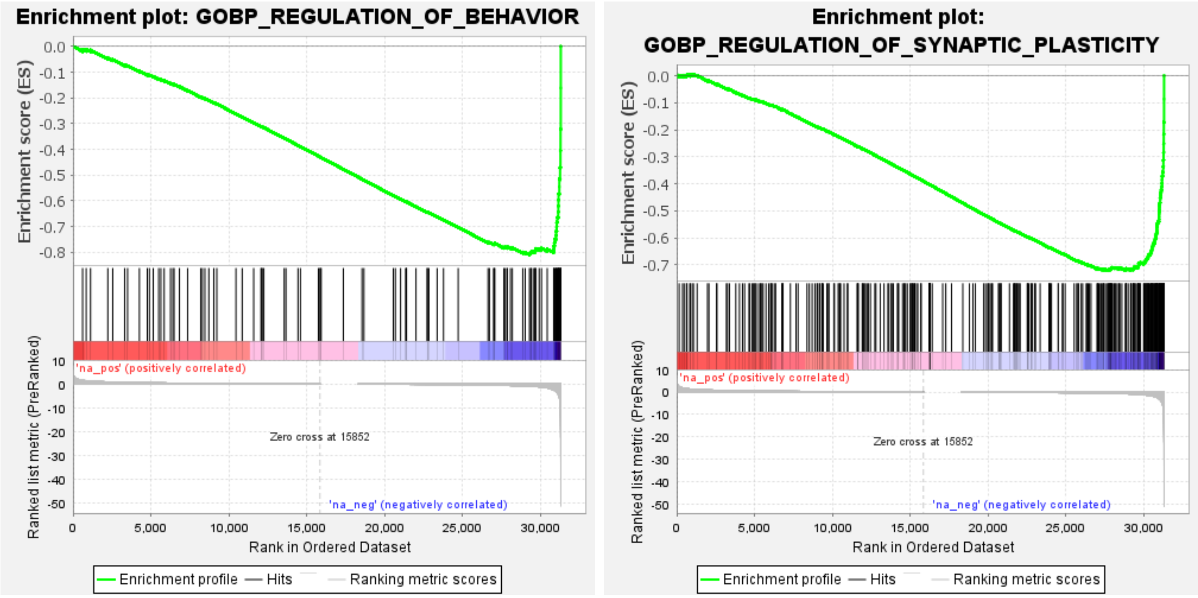


Enrichment of upregulated genes in *R6/1^{Tg/0}; Rarβ^{+/+} vs R6/1^{0/0}; Rarβ^{+/+}* comparison

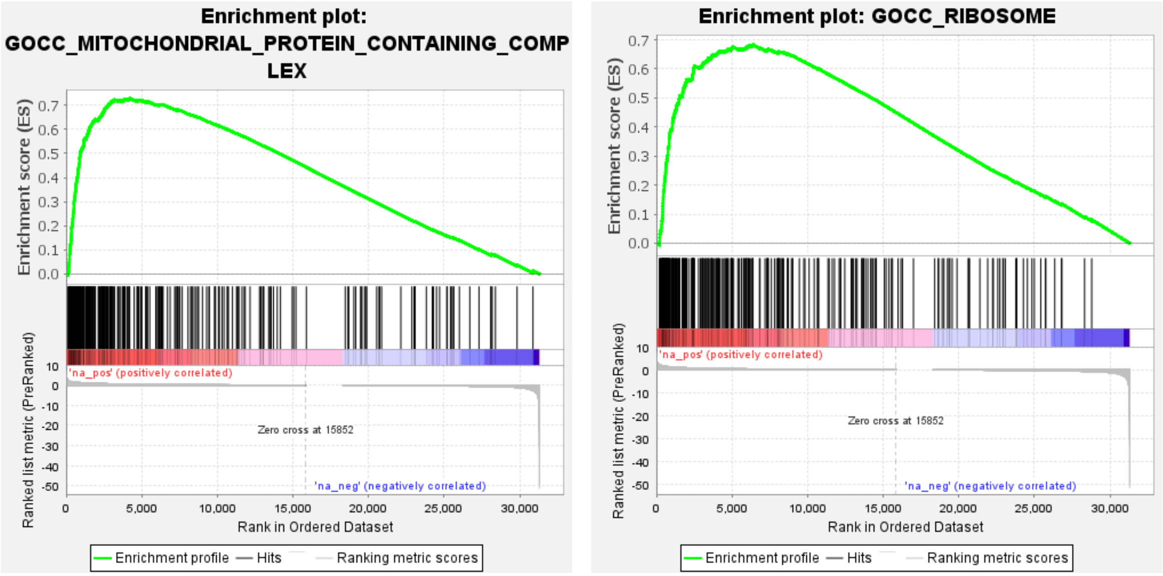


**Fig. S3 A. Ontology gene set enrichment analyses.** (A) Top biological processes associated with significant enrichment of downregulated (upper panel) and upregulated (bottom panel) genes in *R6/1^{Tg/0}; Rarβ^{+/+} vs R6/1^{0/0}; Rarβ^{+/+}* comparison.

**B**

Enrichment of downregulated genes in *R6/1*$^{Tg/0}$*; Rarβ*$^{+/-}$ *vs R6/1*$^{0/0}$*; Rarβ*$^{+/+}$ comparison

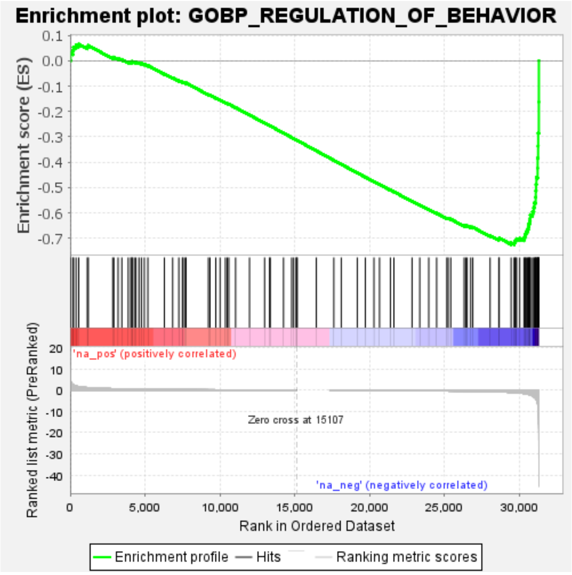


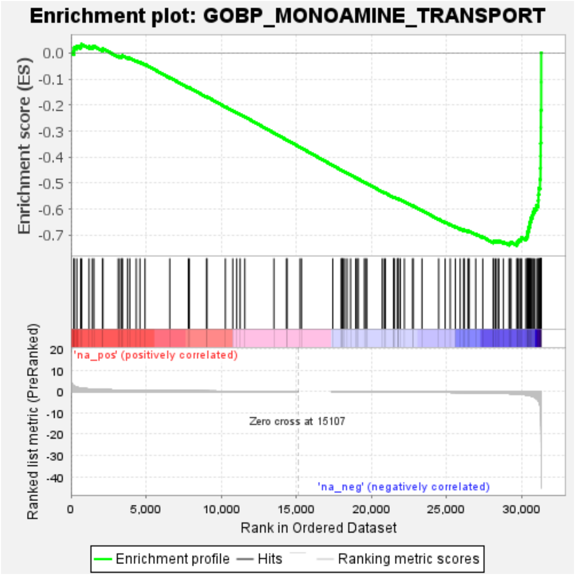


Enrichment of upregulated genes in *R6/1*$^{Tg/0}$*; Rarβ*$^{+/-}$ *vs R6/1*$^{0/0}$*; Rarβ*$^{+/+}$ comparison

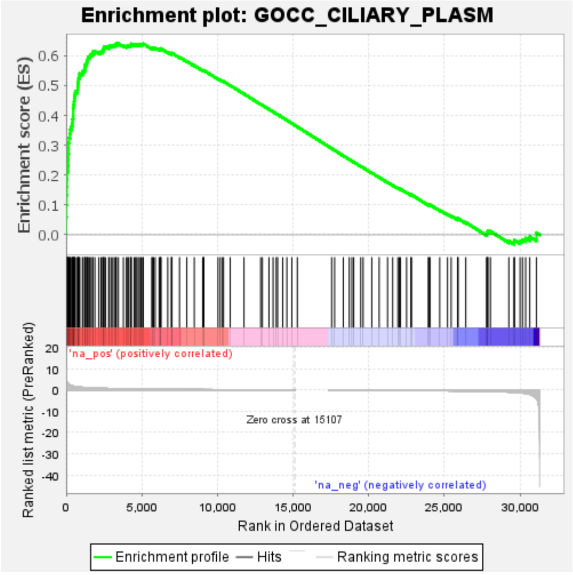


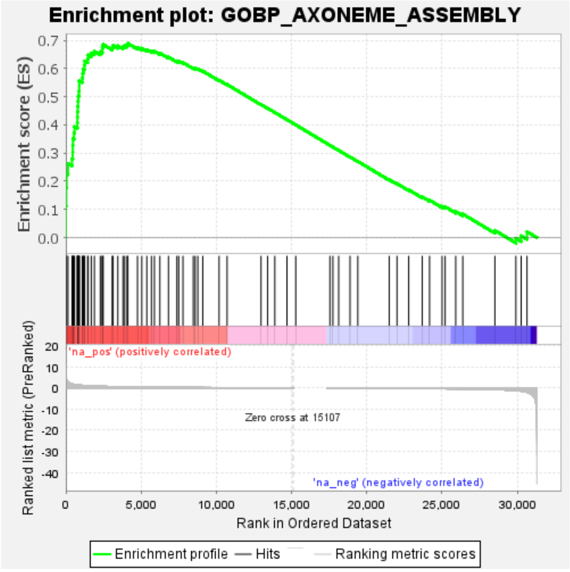


**Fig. S3 B. Ontology gene set enrichment analyses.** (B) Top biological processes associated with significant enrichment of downregulated (upper panel) and upregulated (bottom panel) genes in *R6/1*$^{Tg/0}$*; Rarβ*$^{+/-}$ *vs R6/1*$^{0/0}$*; Rarβ*$^{+/+}$ comparison.

**C**

Enrichment of downregulated genes in *R6/1*$^{Tg/0}$*;Rarβ*$^{+/-}$ *vs R6/1*$^{Tg/0}$*;Rarβ*$^{+/+}$ comparison

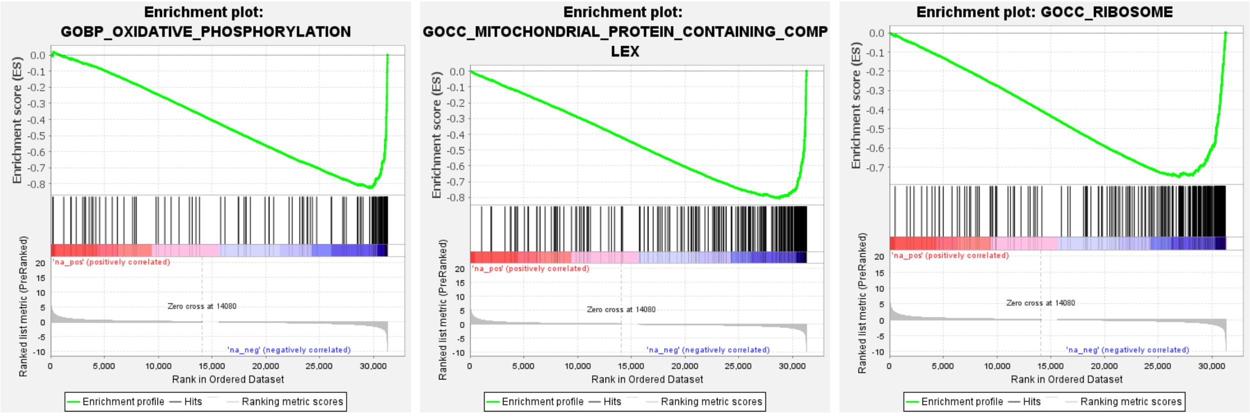


Enrichment of upregulated genes in *R6/1*$^{Tg/0}$*;Rarβ*$^{+/-}$ *vs R6/1*$^{Tg/0}$*;Rarβ*$^{+/+}$ comparison

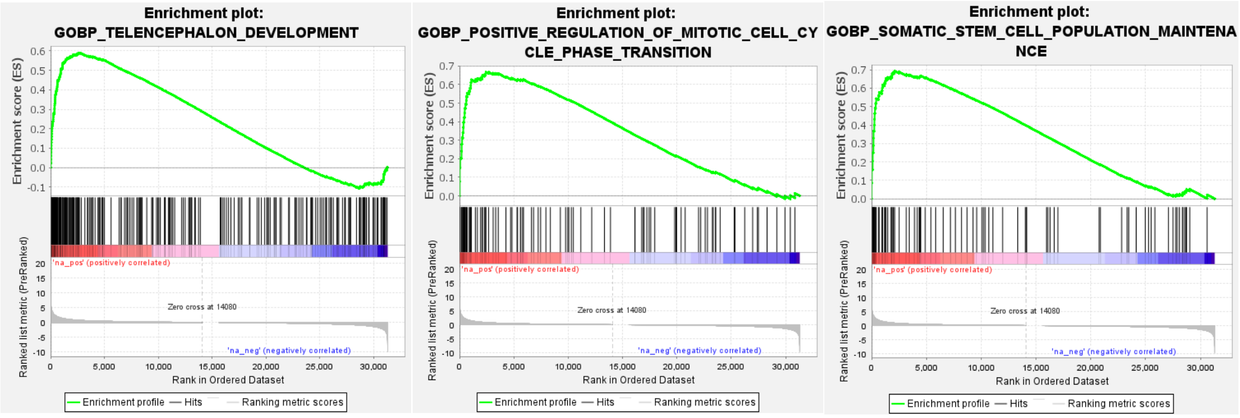


**Fig. S3 C. Ontology gene set enrichment analyses.** (C) Top biological processes associated with significant enrichment of downregulated (upper panel) and upregulated (bottom panel) genes in *R6/1*$^{Tg/0}$*;Rarβ*$^{+/-}$ *vs R6/1*$^{Tg/0}$*;Rarβ*$^{+/+}$ comparison.

**Table S1. DEGs in the Nac of 8 weeks-old males**

**Comparison (p<0.05 cut-off) : *R6/1*** $^{0/0}$ ***; Rarβ*** $^{+/-}$ ***vs. R6/1*** $^{0/0}$ ***; Rarβ*** $^{+/+}$

**Down regulated**

| Gene Name | Log2 Fold Change | p-value | adjusted p-value |
|---|---|---|---|
| Fezf2 | -0,942081639 | 5,78E-07 | 0,002587569 |
| Dlx1 | -0,915813904 | 4,95E-08 | 0,000189542 |
| Fos | -0,893730849 | 3,05E-06 | 0,004507674 |
| Nr4a1 | -0,878597454 | 2,20E-08 | 0,000189542 |
| Neat1 | -0,804103498 | 1,02E-06 | 0,003926086 |
| Stbd1 | -0,804015585 | 2,11E-05 | 0,024756224 |
| Rspo1 | -0,794376251 | 3,40E-05 | 0,027198986 |
| AC159264.1 | -0,781732596 | 8,70E-06 | 0,010405676 |
| Gm43398 | -0,775349657 | 4,21E-05 | 0,027198986 |
| Btg2 | -0,753839108 | 2,46E-05 | 0,027198986 |
| Arc | -0,729230316 | 3,89E-05 | 0,023660602 |
| Foxm1 | -0,728228774 | 0,000128875 | 0,027198986 |
| Lamb3 | -0,727233303 | 5,95E-05 | 0,026193942 |
| Dlk2 | -0,537428775 | 6,65E-05 | 0,027198986 |

**Table S1. DEGs in the Nac of 8 weeks-old males**
**Comparison (p<0.05 cut-off) : *R6/1 $^{0/0}$ ; Rarβ $^{+/-}$* *vs. R6/1 $^{0/0}$ ; Rarβ $^{+/+}$***

**Up regulated**

| Gene Name | Log2 Fold Change | p-value | adjusted p-value |
|---|---|---|---|
| Dclk3 | 0,526090396 | 7,87E-05 | 0,034372086 |
| Gfra1 | 0,621909379 | 3,48E-06 | 0,004507674 |
| C1ql3 | 0,68706693 | 7,28E-05 | 0,037056143 |
| Tenm3 | 0,69781713 | 3,97E-05 | 0,025684981 |
| Hbb-bs | 0,714185374 | 3,59E-06 | 0,004936196 |
| Ano2 | 0,72995225 | 1,30E-05 | 0,013185973 |
| Gm43305 | 0,789633016 | 3,76E-05 | 0,026193942 |
| Hs6st3 | 0,802839367 | 1,55E-05 | 0,023735881 |
| Slc9a7 | 0,822457651 | 1,78E-05 | 0,007861155 |
| Prkg1 | 0,828949157 | 1,69E-05 | 0,023660602 |
| Gm8420 | 0,864965998 | 2,72E-07 | 0,001334492 |
| Gm14094 | 0,872044899 | 1,12E-07 | 0,000327137 |
| Gm10704 | 1,062161149 | 2,28E-08 | 0,000189542 |

**Table S1. DEGs in the Nac of 8 weeks-old males**

**Comparison (p<0.05 cut-off) : *R6/1*$^{Tg/0}$ *; Rarβ*$^{+/+}$ *vs. R6/1*$^{0/0}$ *; Rarβ*$^{+/+}$**

**Down regulated**

| Gene Name | Log2 Fold Change | p-value | adjusted p-value |
|---|---|---|---|
| Cd4 | -2,517099408 | 3,58E-50 | 2,54E-46 |
| Scn4b | -2,096487656 | 6,87E-52 | 8,20E-48 |
| Ptprv | -1,807642166 | 3,44E-21 | 5,10E-18 |
| Lrrc10b | -1,513477606 | 3,21E-23 | 7,24E-20 |
| Gpr6 | -1,37245646 | 9,84E-24 | 2,92E-20 |
| Adora2a | -1,331452988 | 6,64E-31 | 2,79E-27 |
| Syndig1l | -1,311195613 | 2,47E-33 | 2,83E-29 |
| Cartpt | -1,309016589 | 9,11E-18 | 9,95E-15 |
| Penk | -1,302341565 | 1,17E-30 | 2,79E-27 |
| Kcnh4 | -1,288108904 | 2,24E-17 | 1,30E-14 |
| Ier5l | -1,282311101 | 8,13E-20 | 9,79E-17 |
| Ptpn7 | -1,276476409 | 7,88E-12 | 4,13E-09 |
| Car12 | -1,226223099 | 1,88E-16 | 1,86E-13 |
| Rasgrp2 | -1,215918461 | 8,15E-20 | 1,06E-16 |
| Pde10a | -1,203934683 | 7,95E-23 | 8,30E-20 |
| Slc4a11 | -1,178165827 | 1,03E-09 | 4,55E-07 |
| Scube3 | -1,132305867 | 1,38E-11 | 3,91E-09 |
| Ptch2 | -1,097315614 | 2,07E-09 | 6,42E-07 |
| Dmkn | -1,074356178 | 2,12E-08 | 5,97E-06 |
| Rgs9 | -1,073679848 | 2,16E-18 | 1,35E-15 |
| Arc | -1,044738175 | 2,17E-09 | 7,26E-07 |
| Fndc9 | -1,036050191 | 1,64E-10 | 6,16E-08 |
| Pde1b | -1,035153297 | 9,33E-23 | 1,17E-19 |
| Drd2 | -1,019069693 | 2,50E-16 | 2,24E-13 |
| Gpr83 | -1,018660704 | 9,07E-14 | 5,46E-11 |
| Gm36737 | -1,013090026 | 1,75E-07 | 8,47E-05 |
| Drd1 | -1,003963224 | 3,87E-13 | 2,59E-10 |
| Plk5 | -0,993573679 | 5,36E-10 | 1,43E-07 |
| Serpina9 | -0,990805259 | 1,12E-14 | 1,37E-11 |
| Arpp19 | -0,980936984 | 6,67E-13 | 3,09E-10 |
| Rasd2 | -0,965862016 | 4,41E-14 | 1,88E-11 |
| Ppp1r1b | -0,964501017 | 4,43E-23 | 5,94E-20 |
| Trib3 | -0,959214952 | 8,40E-07 | 0,000167159 |
| Adcy5 | -0,958045715 | 2,30E-19 | 1,96E-16 |
| Ido1 | -0,95647167 | 2,84E-07 | 6,12E-05 |
| Gabrd | -0,95566548 | 1,98E-14 | 1,70E-11 |
| 1810020O05Rik | -0,935087731 | 1,92E-06 | 0,00029057 |
| Rasd1 | -0,930590749 | 4,36E-07 | 9,75E-05 |
| Arpp21 | -0,929399125 | 3,85E-14 | 2,00E-11 |
| Tnfrsf25 | -0,92791332 | 3,88E-07 | 8,78E-05 |
| Upb1 | -0,919098833 | 2,28E-06 | 0,00039972 |
| Dusp18 | -0,916530881 | 6,10E-10 | 1,47E-07 |
| St8sia2 | -0,908402521 | 1,05E-06 | 0,000177938 |
| Plxnd1 | -0,895379849 | 2,44E-09 | 8,52E-07 |
| Itga9 | -0,893243639 | 1,56E-07 | 3,36E-05 |
| Ryr1 | -0,887554202 | 3,01E-07 | 4,65E-05 |
| Dnah1 | -0,886943611 | 3,87E-07 | 7,86E-05 |
| Abhd11os | -0,879982748 | 7,21E-06 | 0,000864296 |
| Lingo3 | -0,876670939 | 3,94E-13 | 2,59E-10 |
| Acvr1c | -0,861266572 | 1,68E-06 | 0,000249424 |
| Gsg1l | -0,860382339 | 2,40E-09 | 8,53E-07 |
| Dlx1as | -0,859925846 | 1,05E-05 | 0,001505225 |
| Oprk1 | -0,856317285 | 2,44E-06 | 0,000258897 |
| Rrm2 | -0,856021899 | 1,29E-05 | 0,001597441 |

| | | | |
|---|---|---|---|
| Egr3 | -0,842080049 | 3,40E-10 | 1,43E-07 |
| Wnk4 | -0,841432279 | 1,79E-05 | 0,002336898 |
| Dhrs3 | -0,839086667 | 2,32E-08 | 4,27E-06 |
| Traip | -0,833582259 | 5,33E-08 | 1,43E-05 |
| Dok3 | -0,831322438 | 1,64E-06 | 0,000258897 |
| Inf2 | -0,828637334 | 1,46E-12 | 6,23E-10 |
| Shisa2 | -0,828497174 | 1,44E-05 | 0,001645335 |
| Ddn | -0,82611399 | 5,71E-09 | 1,33E-06 |
| Egr2 | -0,819757632 | 1,44E-06 | 0,001216476 |
| Hrh3 | -0,812684159 | 3,03E-11 | 9,14E-09 |
| Ecel1 | -0,812480506 | 2,34E-08 | 9,51E-06 |
| Itpka | -0,805479804 | 1,16E-10 | 6,00E-08 |
| Pde7b | -0,804273168 | 3,34E-09 | 2,05E-06 |
| Slc5a7 | -0,802165039 | 6,65E-06 | 0,000534458 |
| Itga5 | -0,800679287 | 3,08E-05 | 0,002857084 |
| Spink8 | -0,793080284 | 3,54E-05 | 0,008724568 |
| Gm12992 | -0,789448492 | 1,06E-05 | 0,001174347 |
| Rspo1 | -0,787492394 | 5,38E-05 | 0,005594934 |
| Fam212b | -0,781864753 | 2,89E-08 | 9,04E-06 |
| Gm45645 | -0,778724011 | 6,91E-05 | 0,005376785 |
| Pdyn | -0,777053733 | 3,52E-10 | 1,19E-07 |
| Btg2 | -0,776767459 | 1,06E-05 | 0,00078218 |
| Igfbp5 | -0,775694471 | 3,18E-07 | 8,00E-05 |
| Rin1 | -0,774279316 | 5,67E-12 | 3,53E-09 |
| Mxd3 | -0,772751567 | 8,00E-05 | 0,006798856 |
| AC132384.16 | -0,765650861 | 9,66E-05 | 0,006795273 |
| Kcnip2 | -0,761335535 | 1,33E-11 | 4,13E-09 |
| Npas2 | -0,760900662 | 2,31E-09 | 7,58E-07 |
| Slc18a3 | -0,760704917 | 3,33E-05 | 0,00333492 |
| Tmem200b | -0,760222014 | 6,59E-05 | 0,005637876 |
| AI506816 | -0,759565851 | 7,30E-05 | 0,006273829 |
| Hrk | -0,756425632 | 1,03E-07 | 2,67E-05 |
| Impg1 | -0,749539434 | 3,19E-05 | 0,008389172 |
| Gm15721 | -0,748552552 | 0,000137514 | 0,008943265 |
| Dpy19l3 | -0,747505697 | 3,70E-10 | 1,49E-07 |
| Chrm4 | -0,746832344 | 4,03E-06 | 0,000613102 |
| Pde1c | -0,738599142 | 6,98E-06 | 0,000847897 |
| Lamb3 | -0,73746034 | 3,89E-05 | 0,003960649 |
| Gprin3 | -0,736244892 | 5,69E-06 | 0,000550533 |
| Nnat | -0,734766359 | 8,41E-10 | 1,84E-07 |
| Akap5 | -0,73452808 | 4,93E-08 | 1,43E-05 |
| Ctgf | -0,732897412 | 4,49E-06 | 0,000601156 |
| D7Ertd443e | -0,727623627 | 6,84E-05 | 0,005697848 |
| Foxm1 | -0,722968622 | 0,000165269 | 0,011406773 |
| Tmem40 | -0,722648622 | 0,000217853 | 0,014790945 |
| 1700001O22Rik | -0,718490897 | 0,000225393 | 0,016706471 |
| Fam78a | -0,717831895 | 0,000217028 | 0,014573025 |
| Olfr1393 | -0,712066823 | 0,000228649 | 0,014573025 |
| Pgam2 | -0,706243942 | 6,60E-07 | 9,00E-05 |
| Gm44210 | -0,704644165 | 0,000167669 | 0,011475995 |
| Htr1b | -0,699667903 | 3,42E-05 | 0,002336898 |
| Cyr61 | -0,698880371 | 0,000352814 | 0,018372594 |
| Ugt8a | -0,692021542 | 5,41E-07 | 8,00E-05 |
| Hic1 | -0,691337794 | 0,000426326 | 0,026300592 |
| Nlrp10 | -0,689260526 | 0,000391941 | 0,019746559 |
| Npy2r | -0,689053988 | 0,000444081 | 0,021588709 |
| Gm39043 | -0,688712313 | 0,00012869 | 0,02379052 |
| Cacng4 | -0,688552198 | 1,82E-09 | 6,42E-07 |
| Tmod1 | -0,688396981 | 1,70E-07 | 8,01E-05 |

| | | | |
|---|---|---|---|
| Trh | -0,686770801 | 0,000421208 | 0,023420604 |
| Stra6 | -0,684269508 | 3,15E-06 | 0,000319481 |
| Ankrd63 | -0,680654579 | 1,06E-09 | 4,47E-07 |
| Nuf2 | -0,676497982 | 0,000536796 | 0,024805201 |
| Vip | -0,676275697 | 3,60E-05 | 0,002505046 |
| Camk1g | -0,670340257 | 2,75E-05 | 0,004129057 |
| BC049352 | -0,670139566 | 0,000175301 | 0,029455418 |
| Nr4a1 | -0,670038341 | 8,87E-06 | 0,001174347 |
| Rgs20 | -0,669566814 | 2,85E-07 | 7,37E-05 |
| Cdk1 | -0,667716461 | 0,000336041 | 0,022030741 |
| Cpne9 | -0,667122446 | 0,000328029 | 0,017712233 |
| Fam184b | -0,666459587 | 1,71E-06 | 0,000207156 |
| Sh3rf2 | -0,664038484 | 4,79E-06 | 0,000485998 |
| Kif23 | -0,661935475 | 0,000742976 | 0,038136767 |
| Oprd1 | -0,659610643 | 0,000587979 | 0,029455418 |
| Malat1 | -0,658496116 | 4,93E-06 | 0,000471183 |
| Cplx3 | -0,655718256 | 3,70E-05 | 0,002519354 |
| Peg10 | -0,655542079 | 0,000579176 | 0,026212919 |
| Igfbp4 | -0,654422289 | 5,72E-07 | 0,000120727 |
| Dlx1 | -0,654051866 | 6,13E-05 | 0,00548192 |
| Agbl2 | -0,653014313 | 0,000868373 | 0,037147043 |
| Pmepa1 | -0,651886017 | 8,30E-07 | 0,000170596 |
| Gm37711 | -0,651800511 | 0,000216264 | 0,013076725 |
| Ssc5d | -0,651749622 | 0,000645098 | 0,028193634 |
| Carhsp1 | -0,651131439 | 3,54E-09 | 1,14E-06 |
| Sik1 | -0,650462297 | 1,04E-05 | 0,000878049 |
| Lzts3 | -0,6489793 | 8,18E-11 | 3,01E-08 |
| Epha8 | -0,648408668 | 5,16E-06 | 0,000793367 |
| Cldn11 | -0,646700469 | 9,94E-07 | 0,000197456 |
| Thrsp | -0,645597616 | 0,00039576 | 0,020038494 |
| Jcad | -0,643366976 | 6,50E-07 | 0,000144749 |
| Spata2l | -0,637201437 | 4,75E-06 | 0,001216675 |
| Arhgap33 | -0,636488731 | 4,81E-10 | 1,26E-07 |
| Klf10 | -0,636392673 | 1,42E-05 | 0,001142569 |
| Tpbgl | -0,636343875 | 4,50E-05 | 0,003019489 |
| Uhrf1 | -0,635650475 | 0,001184607 | 0,041923143 |
| Fancd2 | -0,635113421 | 0,000131022 | 0,005697848 |
| Cntnap3 | -0,633203878 | 0,000567075 | 0,020247812 |
| Ccdc187 | -0,627949552 | 4,15E-05 | 0,003960649 |
| Pipox | -0,627042241 | 0,001302272 | 0,049846965 |
| 6430584L05Rik | -0,626734951 | 0,000926169 | 0,035924417 |
| Camk2n1 | -0,624234656 | 8,27E-06 | 0,000734177 |
| Sec14l1 | -0,623891362 | 8,21E-08 | 1,56E-05 |
| Plp1 | -0,62277443 | 3,38E-06 | 0,00039972 |
| Slc24a4 | -0,622740398 | 5,46E-05 | 0,004482239 |
| Prc1 | -0,620142679 | 0,001260229 | 0,049846965 |
| Zmynd10 | -0,619947617 | 0,000558403 | 0,02786248 |
| Prag1 | -0,619753564 | 0,000726079 | 0,030269677 |
| Rasl10a | -0,619441574 | 0,000446625 | 0,021512858 |
| Phactr1 | -0,61840056 | 8,50E-09 | 1,94E-06 |
| Fzd5 | -0,617082926 | 0,000735249 | 0,033393506 |
| Thsd4 | -0,615904271 | 0,001430636 | 0,047705589 |
| Cecr6 | -0,615698768 | 1,79E-05 | 0,002024556 |
| Elfn1 | -0,614631806 | 4,08E-05 | 0,005581578 |
| Slc35d3 | -0,614067195 | 7,40E-05 | 0,006139777 |
| Pdzd2 | -0,612147979 | 3,73E-07 | 8,47E-05 |
| Fbxo32 | -0,611231244 | 7,47E-06 | 0,000878049 |
| Nxph3 | -0,610797236 | 0,000326068 | 0,0194302 |
| Id4 | -0,603691717 | 8,13E-06 | 0,000836458 |

| | | | |
|---|---|---|---|
| Pkp2 | -0,598535758 | 0,000432198 | 0,021294007 |
| Col6a1 | -0,595032078 | 0,000219912 | 0,009136361 |
| Fam117a | -0,594971015 | 0,00023554 | 0,008943265 |
| Mcam | -0,593486088 | 0,000524254 | 0,01953813 |
| Xkr8 | -0,591919456 | 0,000495136 | 0,022865275 |
| Syt2 | -0,590573548 | 0,000238754 | 0,015059372 |
| Rxrg | -0,588272352 | 5,46E-07 | 0,000117753 |
| Gng7 | -0,588132859 | 2,11E-09 | 4,55E-07 |
| Stk32a | -0,587820911 | 0,000490291 | 0,021884073 |
| Tesc | -0,586288062 | 1,62E-06 | 0,00029057 |
| Strn | -0,586220155 | 6,47E-05 | 0,006031312 |
| Nos1 | -0,585043191 | 0,000104451 | 0,008529333 |
| Mal | -0,583237836 | 2,27E-05 | 0,002671157 |
| Spint1 | -0,582864931 | 0,001284892 | 0,049846965 |
| Fut11 | -0,582268967 | 9,50E-05 | 0,005321521 |
| Lsp1 | -0,579348761 | 0,000708681 | 0,02359533 |
| Grk5 | -0,578830616 | 0,000120134 | 0,006188423 |
| Lpar1 | -0,577874287 | 9,61E-05 | 0,005376785 |
| Ldlr | -0,576750521 | 0,000138636 | 0,014239621 |
| Emp2 | -0,576127401 | 0,000146665 | 0,007163537 |
| Msmo1 | -0,572495524 | 6,83E-06 | 0,000836458 |
| Camk4 | -0,571934753 | 1,56E-05 | 0,001373045 |
| Tle2 | -0,571737185 | 0,000246773 | 0,01039042 |
| Bmper | -0,571618537 | 0,000791475 | 0,034714745 |
| A230056P14Rik | -0,570352687 | 9,63E-05 | 0,030656036 |
| Homer1 | -0,568904351 | 4,38E-05 | 0,005334132 |
| Epor | -0,568398441 | 0,001173833 | 0,029427227 |
| Foxo1 | -0,568283795 | 7,99E-06 | 0,001853824 |
| Klf16 | -0,565791062 | 4,35E-06 | 0,000673686 |
| Lypd6 | -0,560613076 | 0,000278953 | 0,015263253 |
| Gpd1 | -0,560170149 | 9,94E-05 | 0,013822198 |
| Grm3 | -0,559405622 | 0,000247706 | 0,014216156 |
| Erf | -0,555398687 | 4,99E-05 | 0,004593717 |
| Gli3 | -0,554960812 | 0,00135307 | 0,046286337 |
| 1110032F04Rik | -0,552545576 | 0,000582527 | 0,021052945 |
| Alk | -0,552042652 | 0,001190924 | 0,042195398 |
| Samd1 | -0,547845262 | 1,16E-05 | 0,001503305 |
| Kcnh3 | -0,546626323 | 5,70E-05 | 0,006296165 |
| Prr18 | -0,543165352 | 2,15E-05 | 0,002336898 |
| Crocc | -0,539335235 | 1,52E-05 | 0,001759134 |
| Ctxn1 | -0,538719123 | 1,59E-06 | 0,00022073 |
| Ntng2 | -0,53710286 | 0,000380444 | 0,029206515 |
| Gm13889 | -0,534320865 | 0,000125608 | 0,00832598 |
| Necab2 | -0,534043553 | 1,11E-06 | 0,000266579 |
| Pitpnm2 | -0,533157538 | 1,50E-06 | 0,000207156 |
| Shkbp1 | -0,532703146 | 0,000589963 | 0,020849348 |
| Cacna2d3 | -0,531262794 | 2,38E-05 | 0,002619526 |
| Aspa | -0,530909816 | 0,001131135 | 0,028780052 |
| Cdc25b | -0,527336109 | 0,000706116 | 0,023585771 |
| Ppp1r9b | -0,522034476 | 3,31E-05 | 0,002660124 |
| Kalrn | -0,519344368 | 9,33E-05 | 0,005697848 |
| Rgs14 | -0,519194428 | 0,000249681 | 0,019700396 |
| Junb | -0,518988065 | 0,000160982 | 0,011267877 |
| Sh2d5 | -0,513106349 | 4,28E-06 | 0,00039972 |
| Krt77 | -0,510797177 | 0,001448001 | 0,033538458 |
| Npas4 | -0,509359266 | 0,001037811 | 0,029638122 |
| Rem2 | -0,508602666 | 5,16E-05 | 0,005085523 |
| Sema4a | -0,507920184 | 0,000231548 | 0,015263253 |
| Adra2c | -0,505763526 | 0,00079816 | 0,036594635 |

| | | | |
|---|---|---|---|
| Gucy1a3 | -0,505667622 | 0,000624355 | 0,021052945 |
| Tbc1d8 | -0,50538841 | 4,42E-05 | 0,004482239 |
| Gpr88 | -0,505115991 | 0,001395811 | 0,040837779 |
| Shank3 | -0,503442012 | 4,82E-05 | 0,003549534 |
| Yjefn3 | -0,502752796 | 0,001223751 | 0,042195398 |
| Ptpn5 | -0,502478352 | 1,99E-06 | 0,00022073 |
| Gpr153 | -0,502128507 | 0,000471689 | 0,033495405 |
| Mobp | -0,501820866 | 0,000156107 | 0,008770828 |
| Il33 | -0,501655711 | 0,001072595 | 0,03019389 |
| Gjc3 | -0,499950873 | 0,000226205 | 0,0194302 |
| Sfrp1 | -0,499444818 | 0,002104167 | 0,047549428 |
| Tead1 | -0,499299499 | 0,001400112 | 0,046833354 |
| Pcp4l1 | -0,498818136 | 8,19E-06 | 0,000734177 |
| Tdrp | -0,497073551 | 0,000576758 | 0,038331979 |
| Prdm16 | -0,495885298 | 0,001677858 | 0,043093757 |
| Tmem44 | -0,49465164 | 8,02E-06 | 0,001049649 |
| Wfs1 | -0,493701239 | 0,000247601 | 0,010012769 |
| Dhcr7 | -0,493265946 | 8,41E-05 | 0,006663604 |
| Them6 | -0,491843462 | 7,73E-05 | 0,006550702 |
| Ccdc88c | -0,490210163 | 0,000296064 | 0,01786665 |
| Spata13 | -0,489253021 | 0,000342741 | 0,017905416 |
| Gfra2 | -0,487579456 | 0,00084411 | 0,047446299 |
| 5330417C22Rik | -0,484614455 | 2,59E-05 | 0,004944014 |
| Lmo2 | -0,484408131 | 0,000605843 | 0,039181225 |
| Etv5 | -0,483877221 | 0,000133934 | 0,009886111 |
| 1700037H04Rik | -0,482054968 | 4,83E-05 | 0,005665641 |
| Scube1 | -0,481137266 | 0,000365574 | 0,020768486 |
| Crym | -0,481085424 | 0,001263597 | 0,038693677 |
| Ppp4r4 | -0,480503437 | 0,000343632 | 0,017905416 |
| Opalin | -0,480455201 | 0,001408844 | 0,047016599 |
| Ppp1r1a | -0,478876941 | 0,000116833 | 0,009247493 |
| Egr1 | -0,478544792 | 0,000909941 | 0,040446609 |
| Arhgap10 | -0,477391843 | 0,000432098 | 0,024613372 |
| Myo5b | -0,477016827 | 0,000199908 | 0,02244007 |
| Sox8 | -0,475819685 | 0,000114932 | 0,008529333 |
| Slc8a2 | -0,474353487 | 0,000412217 | 0,017905416 |
| Tle1 | -0,472202872 | 0,000233741 | 0,015667026 |
| Peli2 | -0,47185774 | 0,000247634 | 0,014216156 |
| Ndnf | -0,470576133 | 0,000188046 | 0,013628947 |
| Tspan2 | -0,468410005 | 3,87E-05 | 0,003892075 |
| Fbxl16 | -0,468206287 | 1,08E-05 | 0,001049649 |
| Baiap2 | -0,466890499 | 1,99E-05 | 0,001373045 |
| Plk2 | -0,466596241 | 2,13E-05 | 0,001759134 |
| Dclk1 | -0,46519051 | 4,14E-05 | 0,003091259 |
| Tlnrd1 | -0,464915052 | 0,00016566 | 0,011475995 |
| Pwwp2b | -0,463094618 | 0,001450161 | 0,038876924 |
| Mchr1 | -0,462539439 | 0,000226313 | 0,015059372 |
| Gad2 | -0,456781124 | 0,000101229 | 0,006139777 |
| Crip2 | -0,456685612 | 0,000199504 | 0,011962195 |
| Nptx2 | -0,45648177 | 0,001682677 | 0,042658244 |
| Bcr | -0,455915129 | 5,17E-05 | 0,003740333 |
| Camkk2 | -0,454793757 | 9,78E-05 | 0,005321521 |
| Dio2 | -0,454378146 | 0,000627583 | 0,027818222 |
| Clic4 | -0,453796795 | 0,000474664 | 0,02244007 |
| Ngef | -0,452445186 | 5,31E-05 | 0,003866203 |
| Car11 | -0,451052412 | 8,83E-06 | 0,000770531 |
| Sez6 | -0,450085921 | 0,000140942 | 0,007915981 |
| Actn2 | -0,449083567 | 0,000434267 | 0,02120026 |
| Gm5607 | -0,448397979 | 0,001329079 | 0,045296116 |

| | | | |
|---|---|---|---|
| Gja1 | -0,448376851 | 4,11E-05 | 0,003181154 |
| Nxph1 | -0,448031042 | 0,000882663 | 0,034379311 |
| Neto1 | -0,446253233 | 0,000571731 | 0,026105885 |
| Insig1 | -0,446250863 | 0,00056912 | 0,029684024 |
| Efnb3 | -0,44617949 | 1,99E-05 | 0,00172217 |
| Caln1 | -0,446072239 | 0,00042833 | 0,018260624 |
| Dock10 | -0,444785357 | 0,001018833 | 0,038136767 |
| Fdft1 | -0,444284546 | 0,000523655 | 0,028378903 |
| Rragd | -0,443020386 | 0,000268442 | 0,014919491 |
| Itpr1 | -0,442999848 | 0,000269581 | 0,013242778 |
| Psd | -0,440226832 | 7,86E-05 | 0,005321521 |
| Camkv | -0,439678539 | 0,00018601 | 0,009653947 |
| Ppp1r16b | -0,43907629 | 0,000308631 | 0,016703363 |
| Nrsn1 | -0,437948626 | 0,001671308 | 0,047705589 |
| Rap1gap | -0,437725966 | 1,82E-05 | 0,001607577 |
| Prkcb | -0,437429317 | 0,000258056 | 0,013250979 |
| Ankrd33b | -0,437250567 | 0,000622838 | 0,030147059 |
| Igsf11 | -0,436681483 | 0,000707909 | 0,034204602 |
| Tyro3 | -0,433556133 | 0,000296047 | 0,014573025 |
| Rab40b | -0,433295968 | 0,001115019 | 0,045052359 |
| Rgs4 | -0,433136514 | 0,001417993 | 0,036859558 |
| Mbp | -0,43305884 | 0,001675671 | 0,046517454 |
| Lhx6 | -0,432869422 | 0,000862327 | 0,037155523 |
| Dpf1 | -0,432828196 | 0,000186155 | 0,021457219 |
| Akt2 | -0,432268183 | 0,000679257 | 0,029125982 |
| Reln | -0,429845082 | 0,000669228 | 0,031272167 |
| Nr1d1 | -0,429806822 | 0,000840619 | 0,038876924 |
| Sept5 | -0,427723131 | 0,000360223 | 0,016715928 |
| Foxp1 | -0,427463797 | 0,000787799 | 0,035805884 |
| Dpy19l1 | -0,426139405 | 0,000127439 | 0,009488864 |
| Josd2 | -0,425300339 | 0,001433101 | 0,047549428 |
| Tomm70a | -0,423655663 | 0,001117551 | 0,030147059 |
| Dbp | -0,423212684 | 0,000102224 | 0,006278328 |
| Ttyh1 | -0,423154813 | 1,11E-05 | 0,000864296 |
| Agfg2 | -0,423120555 | 0,000857242 | 0,038693677 |
| Syt12 | -0,421522361 | 0,000963353 | 0,039556355 |
| Arel1 | -0,421141452 | 0,000121394 | 0,006867686 |
| Ppm1f | -0,419853196 | 0,000486662 | 0,031131166 |
| Rasl10b | -0,414560163 | 4,24E-05 | 0,003145249 |
| Cplx2 | -0,41443491 | 0,000403792 | 0,018133297 |
| Kcnj10 | -0,413975323 | 0,00199156 | 0,046286337 |
| Kazn | -0,408915191 | 0,000181928 | 0,00832598 |
| Grasp | -0,406797672 | 0,000555323 | 0,029206515 |
| Dclk3 | -0,404703046 | 0,001309554 | 0,044948714 |
| Jph4 | -0,404484333 | 0,000207148 | 0,010522243 |
| Stard10 | -0,401666806 | 0,000584319 | 0,026300592 |
| Gnb5 | -0,401090327 | 9,98E-05 | 0,006188423 |
| Slc1a2 | -0,398309339 | 5,20E-05 | 0,003892075 |
| Rgs8 | -0,397898349 | 0,000499822 | 0,02814364 |
| Ptma | -0,397744878 | 0,000973711 | 0,032467027 |
| Car2 | -0,395870332 | 0,001029015 | 0,044348703 |
| Ap1s1 | -0,395054753 | 0,000169958 | 0,007491932 |
| Kctd1 | -0,38489669 | 0,000678645 | 0,033495405 |
| Chst15 | -0,384506074 | 0,00060556 | 0,029756058 |
| Hpca | -0,383970164 | 0,000724519 | 0,027002771 |
| Rab15 | -0,381820321 | 0,001663679 | 0,046611793 |
| Tenm4 | -0,380124026 | 0,001169083 | 0,036859558 |
| Ankrd13b | -0,376358974 | 0,000288156 | 0,01786665 |
| Stac2 | -0,372539666 | 0,001328587 | 0,049846965 |

| | | | |
|---|---|---|---|
| Rcan2 | -0,370448369 | 0,000593895 | 0,01953813 |
| Cpne5 | -0,369869051 | 0,001415864 | 0,041802444 |
| Mast3 | -0,368598999 | 0,001650851 | 0,038876924 |
| Celf5 | -0,362821393 | 0,001949915 | 0,045638335 |
| Rasgrp1 | -0,360141559 | 0,002105381 | 0,047705589 |
| Smarcd1 | -0,355715925 | 0,001143758 | 0,036023887 |
| Gnal | -0,342760671 | 0,000843262 | 0,026212919 |
| Ntrk2 | -0,340183297 | 0,000988199 | 0,032858354 |
| Glul | -0,33304357 | 0,001822763 | 0,043480778 |

**Table S1. DEGs in the Nac of 8 weeks-old males**
**Comparison (p<0.05 cut-off) : *R6/1 $^{Tg/0}$ ; Rarβ $^{+/+}$ vs. R6/1 $^{0/0}$ ; Rarβ $^{+/+}$***

**Up regulated**

| Gene Name | Log2 Fold Change | p-value | adjusted p-value |
|---|---|---|---|
| Snrpn | 0,326948192 | 0,000948526 | 0,032173836 |
| Luc7l | 0,328274684 | 0,001446268 | 0,035924417 |
| Mast1 | 0,391701838 | 0,000825225 | 0,038185879 |
| Gng3 | 0,392287644 | 0,001521483 | 0,038571527 |
| Coa3 | 0,396732947 | 0,001285935 | 0,044348703 |
| Anxa5 | 0,398923181 | 0,000374433 | 0,022227739 |
| Bex2 | 0,400681893 | 6,25E-05 | 0,00433737 |
| Srp9 | 0,408748009 | 0,001037785 | 0,044591608 |
| Yipf2 | 0,416324947 | 0,002279108 | 0,049846965 |
| Zcchc8 | 0,416760095 | 0,00033704 | 0,020650104 |
| Gsto1 | 0,418292647 | 0,000966605 | 0,039331941 |
| Copb1 | 0,419052797 | 0,000728809 | 0,033538458 |
| Higd2a | 0,430156495 | 0,000533806 | 0,028780052 |
| Trir | 0,436960915 | 0,00044131 | 0,038571527 |
| Cd99l2 | 0,440040959 | 0,000413274 | 0,028193634 |
| Gm23935 | 0,443443273 | 0,000574063 | 0,029125982 |
| Stoml1 | 0,444982379 | 0,000378107 | 0,021512858 |
| Capn10 | 0,447015507 | 0,000580057 | 0,029455418 |
| Asl | 0,44857787 | 0,000841479 | 0,049139151 |
| Scg5 | 0,452002251 | 1,05E-05 | 0,000987986 |
| Prkra | 0,452731737 | 0,001931281 | 0,046517454 |
| Pcdh19 | 0,456675269 | 0,000546598 | 0,028193634 |
| Zbtb24 | 0,457816096 | 0,001671305 | 0,042106542 |
| Ahsa2 | 0,463209185 | 0,000524078 | 0,028193634 |
| Acot9 | 0,464909351 | 0,001665443 | 0,040446609 |
| Dis3 | 0,469183098 | 0,001090057 | 0,032467027 |
| Prpsap1 | 0,471819123 | 5,44E-05 | 0,005321521 |
| Skap2 | 0,475904287 | 0,001042716 | 0,030656036 |
| Tmub1 | 0,475963987 | 0,000231861 | 0,014908007 |
| Fam173a | 0,479216138 | 0,000100472 | 0,007698999 |
| Actr5 | 0,480542828 | 0,001076399 | 0,039181225 |
| Nt5c | 0,480772447 | 0,000228167 | 0,01345747 |
| Particl | 0,486363721 | 0,000455624 | 0,017854408 |
| Psme1 | 0,488708611 | 0,000114978 | 0,009140767 |
| Spint2 | 0,48922045 | 0,000551886 | 0,035805884 |
| Cyp4x1 | 0,490672996 | 0,001479987 | 0,048167145 |
| Mlip | 0,491216613 | 0,000799212 | 0,026105885 |
| Sumf2 | 0,492983082 | 0,00069543 | 0,029455418 |
| Dhrs7 | 0,493622982 | 0,000108748 | 0,00832598 |
| Nenf | 0,498647627 | 0,000872205 | 0,038876924 |
| 1700123O20Rik | 0,498744084 | 0,000471628 | 0,01786665 |
| Fars2 | 0,500536935 | 0,000963229 | 0,036594635 |
| Gprasp2 | 0,500839946 | 1,18E-05 | 0,000941638 |
| Psmb10 | 0,503264166 | 0,000477765 | 0,02516032 |
| Cidea | 0,504960015 | 0,001533585 | 0,049846965 |
| Galns | 0,505033671 | 0,00225218 | 0,045296116 |
| Phyh | 0,505900727 | 2,16E-06 | 0,00039972 |
| Zfp367 | 0,512041058 | 0,002005224 | 0,048167145 |
| Srprb | 0,51454064 | 0,000130851 | 0,010012769 |
| Camk2n2 | 0,515511025 | 1,92E-06 | 0,000425605 |
| H2-D1 | 0,515969295 | 0,000836555 | 0,033393506 |
| Nrip2 | 0,518187805 | 0,000456973 | 0,016715928 |
| Gm1976 | 0,52716924 | 0,000340874 | 0,014175226 |
| Nfxl1 | 0,530624358 | 0,001573784 | 0,038876924 |

| | | | |
|---|---|---|---|
| Prr13 | 0,538351799 | 1,24E-05 | 0,001541239 |
| 2210016F16Rik | 0,544987379 | 0,000657503 | 0,022827943 |
| Optn | 0,555937488 | 0,000105757 | 0,005722007 |
| Fkbp14 | 0,563366476 | 0,000255913 | 0,01390984 |
| Ece2 | 0,568452426 | 1,19E-05 | 0,001503305 |
| Atad2b | 0,570579598 | 0,001467646 | 0,047981647 |
| Gm37928 | 0,572148317 | 0,001743401 | 0,038185879 |
| Hbb-bs | 0,574054353 | 0,000102336 | 0,01057814 |
| Abca7 | 0,598675149 | 4,82E-05 | 0,004515021 |
| Adamts17 | 0,600411964 | 0,001293626 | 0,047861867 |
| Fbn1 | 0,601552085 | 0,000610675 | 0,027495268 |
| Hyi | 0,60258242 | 0,000914665 | 0,035786666 |
| Rdh12 | 0,603775944 | 0,00122644 | 0,047861867 |
| Stat1 | 0,612004598 | 0,000345977 | 0,014216156 |
| P4ha3 | 0,613158103 | 0,00071352 | 0,033495405 |
| Gm26917 | 0,618764935 | 0,000321777 | 0,026212919 |
| Slc1a6 | 0,619392497 | 9,99E-05 | 0,006797585 |
| Cdhr1 | 0,630703717 | 8,20E-05 | 0,00474127 |
| 9430021M05Ri | 0,632615103 | 0,001229504 | 0,043248163 |
| Ifit3 | 0,633867992 | 0,000962792 | 0,040644132 |
| Rtp4 | 0,639479472 | 0,001093395 | 0,045786045 |
| Rnf135 | 0,639886109 | 0,000738557 | 0,034295893 |
| 0610009L18Rik | 0,650673332 | 0,000751377 | 0,035149075 |
| Myl6b | 0,65418132 | 4,24E-06 | 0,000438703 |
| Adcyap1 | 0,654760853 | 4,74E-05 | 0,003960649 |
| Kazald1 | 0,658417452 | 0,000745562 | 0,038170883 |
| 2310030G06Rik | 0,658963929 | 0,000684973 | 0,033495405 |
| Eif2ak2 | 0,659390676 | 0,000224743 | 0,015024851 |
| Oasl2 | 0,660845335 | 0,000695109 | 0,033174972 |
| Fahd2a | 0,666036406 | 7,47E-07 | 0,000207156 |
| Impg2 | 0,669163181 | 0,000478618 | 0,026433367 |
| Klhl14 | 0,680638137 | 0,000365937 | 0,018456929 |
| Ggact | 0,701819896 | 8,44E-06 | 0,000917625 |
| Irgm1 | 0,704157493 | 5,05E-05 | 0,003960649 |
| 4930426D05Rik | 0,707463078 | 0,000237599 | 0,014919491 |
| Dusp23 | 0,812057796 | 1,42E-05 | 0,001632956 |
| Sfmbt2 | 0,814949073 | 8,37E-06 | 0,000972438 |
| Irf7 | 0,873935422 | 5,24E-06 | 0,00076127 |
| Gm1840 | 1,051359828 | 6,80E-08 | 2,11E-05 |

**Table S1. DEGs in the Nac of 8 weeks-old males**
**Comparison (p<0.05 cut-off) : *R6/1*** $^{Tg/0}$ ***; Rarβ*** $^{+/-}$ ***vs. R6/1*** $^{0/0}$ ***; Rarβ*** $^{+/+}$

**Down regulated**

| Gene Name | Log2 Fold Change | p-value | adjusted p-value |
|---|---|---|---|
| Scn4b | -2,028579708 | 3,94E-46 | 6,73E-42 |
| Cd4 | -2,016640385 | 1,81E-32 | 9,74E-29 |
| Ptprv | -1,930709257 | 4,59E-24 | 1,03E-20 |
| Cartpt | -1,436053762 | 3,43E-20 | 6,86E-17 |
| Lrrc10b | -1,358124794 | 2,85E-18 | 3,12E-15 |
| Ier5l | -1,321761837 | 6,07E-20 | 1,10E-16 |
| Arc | -1,297893439 | 1,89E-13 | 1,88E-10 |
| Pde10a | -1,281490864 | 4,10E-24 | 1,03E-20 |
| Penk | -1,263385278 | 5,35E-27 | 3,04E-23 |
| Nr4a1 | -1,260108928 | 3,48E-16 | 3,08E-13 |
| Upb1 | -1,179562611 | 1,01E-09 | 3,40E-07 |
| Tnfrsf25 | -1,176395392 | 1,59E-10 | 6,40E-08 |
| Rasgrp2 | -1,164375799 | 2,50E-17 | 2,89E-14 |
| Npas4 | -1,163395288 | 2,61E-13 | 1,71E-10 |
| Gpr6 | -1,161966935 | 1,19E-16 | 1,43E-13 |
| Kcnh4 | -1,151054505 | 1,13E-13 | 7,51E-11 |
| Ptpn7 | -1,127174116 | 7,63E-10 | 2,95E-07 |
| Vip | -1,125504603 | 1,48E-11 | 5,50E-09 |
| Rgs9 | -1,118426852 | 1,26E-18 | 1,66E-15 |
| Abhd11os | -1,114489545 | 8,15E-09 | 2,44E-06 |
| Plk5 | -1,111886971 | 9,25E-12 | 5,50E-09 |
| Adora2a | -1,110368246 | 1,31E-20 | 2,74E-17 |
| Lamb3 | -1,087943553 | 1,86E-09 | 6,78E-07 |
| Obscn | -1,080847877 | 1,08E-08 | 2,16E-06 |
| Gabrd | -1,072397363 | 1,02E-16 | 1,52E-13 |
| Jsrp1 | -1,07078551 | 5,48E-09 | 1,58E-06 |
| Cpne9 | -1,05611718 | 1,47E-08 | 3,72E-06 |
| Plp1 | -1,046097357 | 3,67E-14 | 3,13E-11 |
| Oprk1 | -1,039081493 | 1,26E-08 | 3,30E-06 |
| Fos | -1,034420646 | 8,89E-08 | 1,68E-05 |
| Arpp19 | -1,022418856 | 3,43E-13 | 1,88E-10 |
| Inf2 | -1,007488348 | 1,03E-16 | 1,25E-13 |
| Trib3 | -0,987667188 | 2,50E-07 | 4,37E-05 |
| Rspo1 | -0,96302483 | 6,55E-07 | 0,000107004 |
| Dok3 | -0,955930204 | 5,01E-08 | 1,03E-05 |
| Ugt8a | -0,932993538 | 4,67E-11 | 2,15E-08 |
| Gpr83 | -0,918827257 | 6,35E-11 | 2,45E-08 |
| Fam212b | -0,916656105 | 2,46E-10 | 8,50E-08 |
| Gm36737 | -0,915259256 | 1,62E-06 | 0,000263703 |
| Ppp1r1b | -0,902439499 | 6,57E-19 | 1,25E-15 |
| Tesc | -0,898622918 | 1,31E-12 | 1,24E-09 |
| Syndig1l | -0,892599907 | 2,50E-15 | 1,94E-12 |
| Hrh3 | -0,891335716 | 1,89E-12 | 1,24E-09 |
| Cldn11 | -0,886143447 | 8,11E-11 | 3,24E-08 |
| Tle2 | -0,881429759 | 3,31E-08 | 7,39E-06 |
| 1700016P03Rik | -0,880438226 | 2,72E-06 | 0,000324723 |
| Rasd2 | -0,877156021 | 3,16E-11 | 1,68E-08 |
| Itpka | -0,875720293 | 1,23E-11 | 5,50E-09 |
| 4930452B06Rik | -0,875583404 | 7,91E-09 | 2,42E-06 |
| Doc2g | -0,873871923 | 3,43E-06 | 0,000385351 |
| Ryr1 | -0,864078278 | 8,02E-07 | 0,000113601 |
| Ppp1r1a | -0,863864565 | 1,93E-11 | 1,12E-08 |
| Car12 | -0,863526143 | 1,39E-08 | 3,54E-06 |
| Car7 | -0,8606282 | 1,80E-08 | 4,12E-06 |

| | | | |
|---|---|---|---|
| Fhl2 | -0,851945233 | 1,91E-09 | 7,16E-07 |
| Pde1b | -0,848074978 | 1,02E-14 | 9,19E-12 |
| Rasl10a | -0,846764989 | 2,00E-06 | 0,000278126 |
| Mobp | -0,835693247 | 1,02E-09 | 3,96E-07 |
| Opalin | -0,831866647 | 7,27E-08 | 1,49E-05 |
| Klf10 | -0,830767001 | 3,07E-08 | 7,39E-06 |
| Yjefn3 | -0,826836558 | 2,09E-07 | 2,65E-05 |
| Prss23 | -0,823147293 | 7,38E-07 | 0,000115856 |
| Cplx3 | -0,82234475 | 3,95E-07 | 5,95E-05 |
| Pdlim2 | -0,818363254 | 9,13E-06 | 0,000898479 |
| Tmem40 | -0,815882975 | 2,53E-05 | 0,001526425 |
| Slc4a11 | -0,815236602 | 1,74E-05 | 0,001567787 |
| Pgam2 | -0,814551757 | 2,39E-08 | 5,57E-06 |
| Serpina9 | -0,814142804 | 7,25E-10 | 2,48E-07 |
| D7Ertd443e | -0,813962576 | 9,26E-06 | 0,000905745 |
| Krt9 | -0,813604292 | 1,27E-06 | 0,000188081 |
| Crhbp | -0,811854282 | 9,22E-08 | 1,68E-05 |
| Acvr1c | -0,81077275 | 7,01E-06 | 0,000798865 |
| Them6 | -0,802136564 | 4,89E-10 | 2,20E-07 |
| Gm20379 | -0,79561832 | 6,31E-06 | 0,000664235 |
| Ptch2 | -0,794596445 | 1,42E-05 | 0,001219198 |
| Ccdc42 | -0,793821188 | 3,44E-05 | 0,002596559 |
| Rgs11 | -0,792476035 | 3,17E-07 | 4,72E-05 |
| Mal | -0,787429265 | 2,75E-08 | 6,38E-06 |
| Junb | -0,784067294 | 3,07E-08 | 6,43E-06 |
| Pcp4 | -0,783462934 | 1,61E-09 | 5,98E-07 |
| Traip | -0,77721187 | 6,92E-07 | 9,50E-05 |
| Ccdc187 | -0,77689459 | 6,98E-07 | 0,000102005 |
| Tnfaip6 | -0,775825287 | 2,83E-05 | 0,001659258 |
| Trank1 | -0,773756272 | 7,38E-09 | 2,29E-06 |
| Mbp | -0,772758774 | 5,07E-08 | 8,71E-06 |
| Ntng2 | -0,772699186 | 5,92E-07 | 0,000123833 |
| Wnk4 | -0,77268744 | 6,46E-05 | 0,00455661 |
| Col6a1 | -0,771897964 | 2,53E-06 | 0,000332045 |
| Fam78a | -0,768622997 | 6,72E-05 | 0,004489686 |
| Necab3 | -0,7677553 | 5,48E-09 | 1,50E-06 |
| Vipr1 | -0,767596462 | 9,74E-08 | 2,66E-05 |
| Gm5148 | -0,767550474 | 4,21E-05 | 0,003213397 |
| C030029H02Rik | -0,764350703 | 7,81E-05 | 0,005450006 |
| Tmod1 | -0,76226968 | 1,98E-08 | 6,91E-06 |
| Igfbp6 | -0,757537208 | 1,45E-05 | 0,000986503 |
| Gm15721 | -0,756973306 | 9,57E-05 | 0,006343097 |
| Itga9 | -0,753615889 | 1,17E-05 | 0,001021112 |
| Dnhd1 | -0,753424082 | 0,0001021 | 0,004678877 |
| Drd2 | -0,75245695 | 4,65E-09 | 1,58E-06 |
| 1810020O05Rik | -0,747884617 | 0,000114 | 0,007245152 |
| Gm12992 | -0,747253659 | 3,40E-05 | 0,002574089 |
| Kcnip2 | -0,746448016 | 1,64E-10 | 7,57E-08 |
| Egr2 | -0,745508285 | 8,03E-06 | 0,002315213 |
| Gp1bb | -0,741571414 | 8,61E-07 | 0,000113601 |
| Muc3a | -0,740114381 | 6,37E-05 | 0,004130703 |
| Prr18 | -0,736545574 | 2,55E-08 | 8,41E-06 |
| Stard9 | -0,736397847 | 9,48E-07 | 0,000127193 |
| Itga5 | -0,736313784 | 0,00011921 | 0,006625948 |
| Kcnh3 | -0,733208001 | 1,59E-07 | 2,75E-05 |
| Egr1 | -0,733035085 | 7,41E-07 | 0,000115856 |
| Gm37711 | -0,732244009 | 3,81E-05 | 0,00310087 |
| 1700001O22Rik | -0,731472162 | 0,000154264 | 0,00910148 |
| Slc5a7 | -0,731225167 | 4,36E-05 | 0,002951897 |

| | | | |
|---|---|---|---|
| Gm9754 | -0,73012528 | 4,77E-05 | 0,009434992 |
| Pvalb | -0,729860306 | 2,10E-05 | 0,001616336 |
| Tmem200b | -0,729094163 | 0,000124483 | 0,007738766 |
| Ighm | -0,728635939 | 2,91E-06 | 0,000372388 |
| Dhrs3 | -0,72671143 | 2,21E-06 | 0,000263703 |
| Scube3 | -0,723675777 | 1,91E-05 | 0,001744809 |
| 1110032F04Rik | -0,72353081 | 9,87E-06 | 0,000947345 |
| Drd1 | -0,722797649 | 3,64E-07 | 5,77E-05 |
| Sebox | -0,719377181 | 0,000196506 | 0,010098088 |
| Dpy19l1 | -0,718780174 | 4,83E-10 | 1,87E-07 |
| Fstl3 | -0,716774359 | 0,000204093 | 0,025569188 |
| Gprin3 | -0,715946265 | 1,27E-05 | 0,001097934 |
| Abi3bp | -0,714620178 | 2,90E-05 | 0,002123275 |
| Plekhg4 | -0,711572098 | 0,000237697 | 0,012568358 |
| Gm10282 | -0,709605299 | 0,000178447 | 0,008986641 |
| Neat1 | -0,708828715 | 1,25E-05 | 0,001349693 |
| BC049352 | -0,707525394 | 5,32E-05 | 0,00455661 |
| Epha10 | -0,706625767 | 1,52E-06 | 0,00018087 |
| Ctxn1 | -0,706574374 | 1,33E-09 | 5,03E-07 |
| Adamts16 | -0,705190668 | 0,000162388 | 0,009540169 |
| Krt12 | -0,704033477 | 9,87E-05 | 0,004566722 |
| Rasa4 | -0,702033792 | 2,71E-05 | 0,001950257 |
| Cyr61 | -0,700672029 | 0,000298189 | 0,012696963 |
| Josd2 | -0,696329219 | 4,26E-07 | 7,76E-05 |
| Podn | -0,691957457 | 0,000353247 | 0,016273498 |
| Aspa | -0,688283053 | 3,35E-05 | 0,002797162 |
| Gpd1 | -0,6839757 | 3,67E-06 | 0,000476199 |
| Rin1 | -0,683059889 | 4,63E-09 | 1,40E-06 |
| Gm38413 | -0,681957145 | 0,000360613 | 0,012200125 |
| Syt2 | -0,680729386 | 3,21E-05 | 0,003523616 |
| Olfr1393 | -0,676694743 | 0,000422036 | 0,017624988 |
| Spata2l | -0,67560453 | 2,37E-06 | 0,000332045 |
| Gsg1l | -0,670175504 | 5,86E-06 | 0,000677525 |
| Lingo3 | -0,669742328 | 8,13E-08 | 1,57E-05 |
| Nnat | -0,66813269 | 7,10E-08 | 1,19E-05 |
| Ido1 | -0,667084627 | 0,000342362 | 0,015043644 |
| Pdyn | -0,66329884 | 2,22E-07 | 3,23E-05 |
| Nptx2 | -0,663092784 | 8,50E-06 | 0,000878431 |
| Gadd45b | -0,661823543 | 5,90E-05 | 0,003774685 |
| Pradc1 | -0,660721527 | 4,43E-05 | 0,002984719 |
| Npas2 | -0,660609005 | 4,84E-07 | 6,91E-05 |
| Cabp1 | -0,657773303 | 8,71E-08 | 1,44E-05 |
| Ctgf | -0,657607524 | 5,34E-05 | 0,004301643 |
| Arpp21 | -0,657105682 | 2,33E-07 | 3,36E-05 |
| Gt(ROSA)26Sor | -0,657046556 | 0,000544103 | 0,016486336 |
| Homer1 | -0,656861436 | 4,40E-06 | 0,000532193 |
| Gm33651 | -0,653476498 | 0,000634267 | 0,023741238 |
| Akap13 | -0,646930518 | 2,01E-05 | 0,001500799 |
| Dnase1l2 | -0,644788809 | 0,000548896 | 0,016486336 |
| Epop | -0,642505924 | 5,93E-05 | 0,005564297 |
| Ly6g6e | -0,642187942 | 0,0009155 | 0,024216547 |
| Tpm2 | -0,638379959 | 0,000238284 | 0,011090009 |
| Cnr1 | -0,637995635 | 6,10E-05 | 0,004550132 |
| Aldh1a1 | -0,637948444 | 2,05E-05 | 0,001616336 |
| Htr4 | -0,637750896 | 0,00058 | 0,021049635 |
| Ddn | -0,634527174 | 1,33E-05 | 0,001349693 |
| Pls1 | -0,632994934 | 0,000149487 | 0,008935774 |
| Pmepa1 | -0,632418557 | 3,50E-06 | 0,000461314 |
| Plxnd1 | -0,631919844 | 3,82E-05 | 0,004079224 |

| | | | |
|---|---|---|---|
| Fa2h | -0,627463227 | 0,000130692 | 0,007554263 |
| Robo3 | -0,627163252 | 0,001119701 | 0,035216109 |
| Car2 | -0,62698653 | 5,16E-07 | 7,76E-05 |
| Tbc1d8 | -0,626213114 | 9,79E-07 | 0,000116673 |
| Col27a1 | -0,625025713 | 0,000289892 | 0,010463545 |
| Camk2n1 | -0,622381636 | 1,54E-05 | 0,001500511 |
| Hpca | -0,622080914 | 1,31E-07 | 2,65E-05 |
| Sst | -0,620450051 | 1,42E-05 | 0,001413493 |
| Plekhh1 | -0,618375147 | 0,000100165 | 0,005590453 |
| Gucy2g | -0,616556531 | 0,001070539 | 0,027121075 |
| Ppp1r14a | -0,616304169 | 0,000996762 | 0,032356451 |
| Mag | -0,616178424 | 0,000205638 | 0,009332634 |
| Wnt5b | -0,615128124 | 0,001500876 | 0,046524131 |
| Tac1 | -0,614983691 | 1,95E-05 | 0,001760435 |
| Cryab | -0,614825966 | 1,99E-05 | 0,001547169 |
| Gm34583 | -0,613649127 | 0,000693803 | 0,027634227 |
| Gjc3 | -0,613480258 | 1,09E-05 | 0,001472796 |
| Pdzd7 | -0,612239814 | 0,000587011 | 0,021194288 |
| Lmo2 | -0,611881214 | 2,49E-05 | 0,00193659 |
| Rreb1 | -0,610842967 | 0,000469392 | 0,018799936 |
| Ldlr | -0,61059191 | 7,97E-05 | 0,007027741 |
| Gucy1a3 | -0,610135744 | 5,61E-05 | 0,003285817 |
| Rnf112 | -0,609753678 | 2,09E-08 | 5,65E-06 |
| Myh7b | -0,608654105 | 0,000805113 | 0,027929413 |
| Col19a1 | -0,608232424 | 0,001686521 | 0,046524131 |
| Rxrg | -0,60812134 | 5,80E-07 | 8,65E-05 |
| Slc35d3 | -0,607387976 | 0,000122687 | 0,007810422 |
| Btg2 | -0,606568923 | 0,000638958 | 0,022661017 |
| Adra2b | -0,604756529 | 0,001786552 | 0,038716871 |
| Snhg6 | -0,604132594 | 0,000725565 | 0,028284447 |
| Epor | -0,601524644 | 0,00067492 | 0,027121075 |
| Npy5r | -0,599850718 | 0,001189446 | 0,040160724 |
| Nr4a2 | -0,598939989 | 0,00105873 | 0,034678153 |
| 2900079G21Rik | -0,598858916 | 0,000629113 | 0,022846338 |
| Per2 | -0,595603694 | 4,81E-06 | 0,000765554 |
| Dio2 | -0,595401962 | 1,34E-05 | 0,001403384 |
| Adamts3 | -0,592432579 | 0,000213774 | 0,011751398 |
| Usp28 | -0,591512287 | 9,94E-05 | 0,006789509 |
| Strip2 | -0,59104402 | 0,000163437 | 0,007766772 |
| Bcas1 | -0,590763326 | 7,01E-07 | 0,000113601 |
| Chrd | -0,589034641 | 0,000107119 | 0,007055133 |
| Zcwpw1 | -0,588339782 | 0,000503539 | 0,018799936 |
| Epha8 | -0,586765908 | 5,81E-05 | 0,004493725 |
| Mas1 | -0,585941138 | 0,001315893 | 0,04351001 |
| Camk4 | -0,585106668 | 1,81E-05 | 0,001349693 |
| Acy1 | -0,584456324 | 0,000413836 | 0,017330285 |
| Actn2 | -0,584451641 | 9,34E-06 | 0,000810869 |
| Ankrd35 | -0,584046938 | 0,000462829 | 0,018760095 |
| Lpar1 | -0,583334256 | 0,000119818 | 0,007554263 |
| Chrdl1 | -0,58242031 | 0,001118875 | 0,033973911 |
| Id2 | -0,580991376 | 4,79E-06 | 0,000515513 |
| Tspan2 | -0,579419415 | 9,05E-07 | 0,000122802 |
| Ttc9b | -0,578977172 | 0,000144159 | 0,007055133 |
| Dusp14 | -0,577436095 | 1,70E-05 | 0,001472796 |
| Chml | -0,574797689 | 0,001156724 | 0,036081126 |
| Ermn | -0,57456392 | 7,59E-05 | 0,00552384 |
| Idi1 | -0,573329046 | 7,95E-05 | 0,005669782 |
| Adcy5 | -0,572889334 | 2,25E-07 | 4,18E-05 |
| Col11a2 | -0,569448055 | 0,001074552 | 0,034007784 |

| | | | |
|---|---|---|---|
| Hapln2 | -0,567283806 | 0,001220764 | 0,036230441 |
| C78859 | -0,566133527 | 0,001066095 | 0,03790265 |
| Pitpnm2 | -0,564411815 | 9,30E-07 | 0,000139261 |
| Cck | -0,561660837 | 0,000481554 | 0,021194288 |
| Cttnbp2 | -0,560553304 | 2,53E-05 | 0,001760435 |
| Dkkl1 | -0,560480205 | 0,001197146 | 0,036961721 |
| Per3 | -0,560458589 | 0,000100043 | 0,008465081 |
| Stard10 | -0,560441779 | 3,74E-06 | 0,00038572 |
| Il1rap | -0,558734367 | 0,000216901 | 0,010230795 |
| Ldhd | -0,557297566 | 0,000688708 | 0,0195085 |
| Ivns1abp | -0,556740859 | 0,000426935 | 0,015312889 |
| Car11 | -0,555172846 | 1,51E-07 | 2,92E-05 |
| Rtn4r | -0,551999721 | 0,000530335 | 0,023380048 |
| Pik3c2b | -0,551301561 | 0,000331997 | 0,014659295 |
| Rgs14 | -0,549953323 | 0,000158935 | 0,007635841 |
| Litaf | -0,549385769 | 0,000627122 | 0,022831382 |
| Stard4 | -0,548924755 | 0,000266118 | 0,011910619 |
| Cecr6 | -0,548504675 | 0,000195183 | 0,009997739 |
| Npy | -0,545497985 | 3,26E-05 | 0,003551679 |
| Gria3 | -0,543968374 | 0,000373026 | 0,013868612 |
| Mog | -0,542018619 | 0,001156159 | 0,035973315 |
| Rhpn1 | -0,541959996 | 0,001449614 | 0,046847964 |
| Qdpr | -0,54024945 | 5,87E-06 | 0,000677525 |
| R3hdm1 | -0,539683202 | 6,26E-05 | 0,004566722 |
| Ndst4 | -0,538802748 | 0,001126884 | 0,033424056 |
| Baiap2 | -0,538016281 | 2,22E-06 | 0,000305355 |
| Ryr3 | -0,537074809 | 0,000635169 | 0,026062942 |
| Hebp1 | -0,536880559 | 0,001234638 | 0,037561454 |
| Grasp | -0,535867141 | 1,14E-05 | 0,001518429 |
| Malat1 | -0,535560047 | 0,000293861 | 0,014659295 |
| Nrn1 | -0,534291865 | 2,54E-05 | 0,001852177 |
| Ankrd33b | -0,534079902 | 5,09E-05 | 0,003147889 |
| Egr3 | -0,533184393 | 0,000106652 | 0,006249469 |
| Fam193b | -0,53026726 | 7,75E-05 | 0,004316793 |
| Nr4a3 | -0,528954658 | 0,000827723 | 0,032128139 |
| Kctd6 | -0,525995206 | 5,84E-05 | 0,00455661 |
| Msmo1 | -0,525739977 | 6,27E-05 | 0,005812353 |
| Ccnyl1 | -0,524741832 | 0,000794293 | 0,026430347 |
| Sema4a | -0,52346528 | 0,000224572 | 0,011198409 |
| Fam117a | -0,522910528 | 0,001463938 | 0,047132686 |
| Hrk | -0,52068883 | 0,0003561 | 0,014659295 |
| Pllp | -0,519613164 | 0,00063067 | 0,026910112 |
| Sox2ot | -0,518469958 | 0,000234282 | 0,012661373 |
| Fbxo2 | -0,518428606 | 0,000263342 | 0,011824141 |
| Pex5l | -0,517458961 | 0,000587281 | 0,01943805 |
| Syt12 | -0,517449787 | 8,72E-05 | 0,005382905 |
| Fam98c | -0,515713879 | 5,00E-05 | 0,004982478 |
| Nin | -0,514701141 | 0,000126786 | 0,010165278 |
| Slc25a37 | -0,514678814 | 0,001132051 | 0,040160724 |
| Nhp2 | -0,512474317 | 0,000176756 | 0,010634218 |
| Mapk11 | -0,511462942 | 0,000252913 | 0,013626377 |
| Tspoap1 | -0,508891647 | 0,000148544 | 0,00893146 |
| Gm43597 | -0,508556302 | 0,000738958 | 0,027144889 |
| Ap1s1 | -0,507647323 | 3,42E-06 | 0,000332045 |
| Tmem44 | -0,507186747 | 1,02E-05 | 0,000878431 |
| Flnb | -0,507161212 | 0,000268962 | 0,011943137 |
| A230056P14Rik | -0,506270912 | 0,000439021 | 0,035864318 |
| Rab40b | -0,50538686 | 0,000224715 | 0,015312889 |
| Lin7b | -0,50535584 | 0,00037522 | 0,022658622 |

| | | | |
|---|---|---|---|
| Mchr1 | -0,500011444 | 0,000115099 | 0,006625948 |
| Akt2 | -0,498280604 | 0,000149491 | 0,008935774 |
| Sntb2 | -0,497675581 | 0,001573666 | 0,049728944 |
| Kcnab3 | -0,497640843 | 0,00068505 | 0,023741238 |
| Strn | -0,49644299 | 0,000962277 | 0,031366512 |
| C1qtnf4 | -0,494926264 | 4,98E-05 | 0,002984866 |
| B3galt2 | -0,49248371 | 0,000199285 | 0,011640449 |
| Insig1 | -0,492103049 | 0,00022939 | 0,012661373 |
| Arl4d | -0,492027282 | 0,000821884 | 0,030813397 |
| Nefm | -0,491603704 | 0,0003475 | 0,016486336 |
| Slc24a4 | -0,489032535 | 0,001859191 | 0,048773847 |
| Rem2 | -0,488431813 | 0,000167666 | 0,012285197 |
| Ptprd | -0,488102517 | 0,001182595 | 0,036419619 |
| Igfbp4 | -0,487905238 | 0,000296942 | 0,015312889 |
| Chn1 | -0,487689949 | 7,70E-05 | 0,005382905 |
| Crocc | -0,48639952 | 0,000159025 | 0,011943137 |
| 1700037H04Rik | -0,485998463 | 7,69E-05 | 0,005382905 |
| H2afj | -0,485129251 | 0,001484471 | 0,049728944 |
| Clcn2 | -0,483456066 | 4,20E-05 | 0,002711412 |
| Pcp4l1 | -0,482376004 | 3,26E-05 | 0,002723749 |
| Scube1 | -0,48180238 | 0,000522909 | 0,023536163 |
| Ccdc88c | -0,481329972 | 0,000555417 | 0,019323124 |
| Ier5 | -0,478777228 | 0,00036806 | 0,022282956 |
| Sez6 | -0,475100465 | 0,000105366 | 0,006789509 |
| Tgfa | -0,474733743 | 0,000294803 | 0,015312889 |
| Fam131a | -0,473689798 | 0,000630059 | 0,025924945 |
| Kcnj4 | -0,473103067 | 5,79E-05 | 0,004375071 |
| Ssh3 | -0,472076414 | 0,000655842 | 0,024216547 |
| Lrrk2 | -0,471898553 | 0,000801648 | 0,026611865 |
| Ptpn5 | -0,470108198 | 1,89E-05 | 0,001736212 |
| Matk | -0,470072067 | 2,10E-05 | 0,001853944 |
| Abcc8 | -0,468839096 | 0,000862408 | 0,028306734 |
| Dlk2 | -0,468314375 | 0,000347885 | 0,014659295 |
| Kalrn | -0,467331993 | 0,000646066 | 0,026276365 |
| Aifm3 | -0,466421713 | 0,000136152 | 0,006789509 |
| Myo9b | -0,464095187 | 0,00038449 | 0,015345528 |
| Gm996 | -0,463033835 | 1,42E-05 | 0,001097934 |
| Arhgap33 | -0,461853473 | 1,41E-05 | 0,001413493 |
| Ccm2 | -0,461406369 | 0,000291271 | 0,012051399 |
| Scn1b | -0,460908405 | 8,45E-06 | 0,000922715 |
| Stac2 | -0,460839788 | 0,000128784 | 0,006521718 |
| Dgkz | -0,460442212 | 0,000451242 | 0,016061635 |
| Hras | -0,459915913 | 3,90E-05 | 0,003147889 |
| Kcnt1 | -0,458632079 | 0,000657165 | 0,022044349 |
| Rab26 | -0,458373708 | 0,001090271 | 0,039305292 |
| Rprml | -0,457452865 | 0,000860419 | 0,027780469 |
| Rasl11b | -0,456250513 | 0,000307477 | 0,012948152 |
| Fabp5 | -0,45349857 | 0,00059285 | 0,02265983 |
| Ccdc85b | -0,452645227 | 0,000189852 | 0,008465081 |
| Phactr1 | -0,452451265 | 5,05E-05 | 0,003898809 |
| Ckb | -0,45208256 | 0,000348189 | 0,016486336 |
| Nktr | -0,451692864 | 0,000846342 | 0,031279484 |
| Ptms | -0,446502952 | 4,88E-05 | 0,002951897 |
| Rbck1 | -0,444827086 | 0,000295809 | 0,012200125 |
| Jcad | -0,441767916 | 0,000923 | 0,027817822 |
| Kcnip3 | -0,440956308 | 0,000845345 | 0,026398815 |
| Samd1 | -0,437748067 | 0,0006932 | 0,028306734 |
| Crip2 | -0,437631543 | 0,000570677 | 0,022036046 |
| Fbxo32 | -0,437175053 | 0,001765277 | 0,047869249 |

| | | | |
|---|---|---|---|
| Psd | -0,435482277 | 0,000168341 | 0,009807318 |
| Vamp1 | -0,434787967 | 0,000201838 | 0,00893146 |
| Fam57a | -0,434294092 | 0,000684626 | 0,023741238 |
| Sept4 | -0,433814332 | 0,001348756 | 0,036250425 |
| Ssbp4 | -0,432244317 | 0,001397226 | 0,045690241 |
| Necab2 | -0,4307017 | 0,000154866 | 0,007523001 |
| Dhcr7 | -0,429982525 | 0,000901165 | 0,034007784 |
| Chst1 | -0,42805537 | 0,000894628 | 0,032500055 |
| Cacng4 | -0,4277625 | 0,000310188 | 0,012568358 |
| Grin2c | -0,425462863 | 0,001639853 | 0,046771264 |
| Hmgcs1 | -0,424018564 | 0,000643179 | 0,026276365 |
| Sh2d5 | -0,421754303 | 0,000272123 | 0,011252402 |
| Kcnn2 | -0,421057763 | 0,001775876 | 0,046939766 |
| Prkcb | -0,420877082 | 0,000685518 | 0,022705097 |
| Smarcd3 | -0,420684698 | 8,04E-05 | 0,00552384 |
| Inafm1 | -0,419721752 | 0,001307617 | 0,039305292 |
| Arhgef2 | -0,418561213 | 0,000282 | 0,011471543 |
| Snap25 | -0,417377604 | 0,001026674 | 0,02897164 |
| Lzts3 | -0,414974893 | 6,57E-05 | 0,004726554 |
| Phldb1 | -0,413676717 | 0,000991035 | 0,028284447 |
| Brk1 | -0,412028523 | 0,000469012 | 0,020755703 |
| Camkk2 | -0,411256842 | 0,000672919 | 0,0270403 |
| 5330417C22Rik | -0,410264304 | 0,000579673 | 0,019903023 |
| Sh3gl3 | -0,409537489 | 0,000908147 | 0,030190208 |
| Mpc1 | -0,406688273 | 0,00035172 | 0,013729585 |
| Fbxw7 | -0,405458267 | 0,00154331 | 0,049166946 |
| Focad | -0,404600021 | 0,001088768 | 0,031119914 |
| Tyro3 | -0,40299126 | 0,001160417 | 0,031401932 |
| Kazn | -0,402328155 | 0,000390737 | 0,018131008 |
| Dnajb5 | -0,40208098 | 0,000276939 | 0,012696963 |
| Gatm | -0,399014992 | 0,001383217 | 0,036961721 |
| Camkv | -0,398681934 | 0,001075657 | 0,037229331 |
| Dclk1 | -0,394706376 | 0,000801445 | 0,025569188 |
| Ngef | -0,393401611 | 0,000711861 | 0,027817822 |
| Klf16 | -0,392652664 | 0,002039994 | 0,049728944 |
| Cyp46a1 | -0,392418462 | 0,000238473 | 0,012568358 |
| Rnd2 | -0,383782312 | 0,000638022 | 0,023741238 |
| Dbp | -0,383564219 | 0,000700899 | 0,027769179 |
| Rogdi | -0,382769271 | 0,000505876 | 0,018079146 |
| Col4a2 | -0,380841436 | 0,001392044 | 0,039080186 |
| Fbxl16 | -0,378714263 | 0,000618642 | 0,025569188 |
| Ankrd13b | -0,376631768 | 0,000487224 | 0,021379914 |
| Zfyve28 | -0,375950956 | 0,001644283 | 0,044297307 |
| Gnb5 | -0,372252936 | 0,000519087 | 0,017818471 |
| Plk2 | -0,370050953 | 0,001166375 | 0,03254917 |
| Asphd2 | -0,365360046 | 0,001001906 | 0,029332462 |
| Carhsp1 | -0,359331515 | 0,001678552 | 0,047515058 |
| Fam160b2 | -0,35412083 | 0,001557433 | 0,040160724 |
| Sh3glb2 | -0,349171684 | 0,001598573 | 0,039765619 |
| Rap1gap | -0,344395581 | 0,001197111 | 0,040160724 |
| Znrf1 | -0,342487216 | 0,00162192 | 0,046524131 |
| Speg | -0,341970084 | 0,001591806 | 0,03972919 |
| Pcsk2 | -0,338376296 | 0,001718381 | 0,04351001 |

**Table S1. DEGs in the Nac of 8 weeks-old males**

**Comparison (p<0.05 cut-off) : *R6/1* $^{Tg/0}$ *; Rarβ* $^{+/-}$ *vs. R6/1* $^{0/0}$ *; Rarβ* $^{+/+}$**

**Up regulated**

| Gene Name | Log2 Fold Change | p-value | adjusted p-value |
|---|---|---|---|
| Usp22 | 0,330368542 | 0,001335938 | 0,036081126 |
| Sod2 | 0,348326741 | 0,001329894 | 0,043739822 |
| Gdi1 | 0,372186484 | 0,001393446 | 0,045670654 |
| Acp2 | 0,379903612 | 0,001710604 | 0,041697477 |
| Copb2 | 0,396326741 | 0,000617571 | 0,020202781 |
| Ckap5 | 0,396715483 | 0,000840894 | 0,031145685 |
| Dcaf5 | 0,400583227 | 0,001467874 | 0,037229331 |
| Brd3 | 0,401083742 | 0,000806562 | 0,038514235 |
| Pam | 0,401738118 | 0,001261819 | 0,033424056 |
| Txndc5 | 0,403164255 | 0,000538108 | 0,02302098 |
| Prune2 | 0,404488275 | 0,000361699 | 0,013568262 |
| Tspyl4 | 0,410508331 | 0,000136752 | 0,008366751 |
| Gnl3l | 0,411172392 | 0,000101891 | 0,006625948 |
| Stim1 | 0,412694516 | 0,000940722 | 0,033814066 |
| Hmg20a | 0,415160857 | 0,00096286 | 0,028507495 |
| Copa | 0,41883099 | 0,000794927 | 0,025429806 |
| Gprasp1 | 0,419195059 | 0,000200466 | 0,01125628 |
| Suv39h1 | 0,421899629 | 0,000926625 | 0,029087631 |
| Prkacb | 0,422129093 | 8,50E-05 | 0,005677413 |
| Scaf11 | 0,42232591 | 0,000363232 | 0,014053493 |
| Hsd17b11 | 0,422440665 | 0,000699195 | 0,024125512 |
| Snx4 | 0,426251559 | 0,000376829 | 0,018332621 |
| Gng4 | 0,426770369 | 0,000489197 | 0,01943805 |
| Snx12 | 0,427029429 | 0,000853475 | 0,040108739 |
| Rbl2 | 0,428234174 | 0,000820176 | 0,025924945 |
| Hap1 | 0,435673902 | 0,000390437 | 0,014886471 |
| Crebl2 | 0,436590028 | 0,000588794 | 0,030710835 |
| Mb21d2 | 0,444628633 | 0,000273202 | 0,012659268 |
| Epb41l4b | 0,447992048 | 0,000774822 | 0,037229331 |
| Lrrc41 | 0,448110236 | 0,000923064 | 0,04253675 |
| Armcx4 | 0,448645776 | 0,001054865 | 0,029370247 |
| Polr1a | 0,455403946 | 0,000290282 | 0,012568358 |
| Dpysl3 | 0,455860302 | 0,000471485 | 0,016486336 |
| Sufu | 0,457585307 | 0,001362499 | 0,041697477 |
| Prepl | 0,461699381 | 0,000250652 | 0,012926205 |
| Zfp146 | 0,461889129 | 0,001257129 | 0,042962097 |
| Fbxo10 | 0,464128165 | 0,000207966 | 0,011943137 |
| Chd3 | 0,467431681 | 0,001004439 | 0,029341138 |
| Ptgfrn | 0,47506495 | 0,000662364 | 0,032973347 |
| Rcan3 | 0,48132439 | 0,000599687 | 0,020482963 |
| Cachd1 | 0,485896509 | 0,000828702 | 0,027144889 |
| Pigv | 0,488612562 | 0,001969484 | 0,041595434 |
| Tcerg1 | 0,488615589 | 2,64E-05 | 0,00181838 |
| Bag2 | 0,489886126 | 0,002491646 | 0,049728944 |
| Sumf2 | 0,490810868 | 0,000995443 | 0,032356451 |
| Dis3 | 0,492601737 | 0,000794118 | 0,028306734 |
| Hrasls | 0,497562478 | 0,000288183 | 0,01826221 |
| Fkbp14 | 0,498295555 | 0,001528548 | 0,048844913 |
| Optn | 0,498588276 | 0,000705237 | 0,027780469 |
| Parva | 0,498725251 | 0,000312993 | 0,0195085 |
| Dnali1 | 0,508898519 | 0,001029455 | 0,046658714 |
| Gucy2f | 0,509181121 | 0,000748604 | 0,032018205 |
| Spint2 | 0,516006592 | 0,000392334 | 0,016486336 |
| Scaper | 0,52008755 | 0,000439051 | 0,025429806 |

| | | | |
|---|---|---|---|
| Stk3 | 0,520682472 | 0,000948641 | 0,030200681 |
| Brs3 | 0,523955526 | 0,00135917 | 0,049859764 |
| Cd99l2 | 0,525441448 | 4,51E-05 | 0,003548822 |
| Taok3 | 0,526159542 | 3,90E-05 | 0,002705021 |
| Arid5b | 0,528312104 | 0,000366629 | 0,018079146 |
| Iqca | 0,539077148 | 0,001034607 | 0,046771264 |
| Zfp110 | 0,542756744 | 0,000310202 | 0,013912786 |
| Zfp521 | 0,543580361 | 0,001245993 | 0,029816243 |
| Ppp1r26 | 0,549062208 | 0,000367504 | 0,017803962 |
| Trim66 | 0,550610378 | 8,66E-05 | 0,005077908 |
| Gprasp2 | 0,553370435 | 3,05E-06 | 0,000319554 |
| Glg1 | 0,556337452 | 0,000215196 | 0,009666106 |
| Ahi1 | 0,559877106 | 1,44E-05 | 0,001425028 |
| Lgals3bp | 0,561900769 | 0,001081419 | 0,032370704 |
| Gm1976 | 0,566092852 | 0,000173416 | 0,008614758 |
| Irgm1 | 0,568084578 | 0,001204764 | 0,029087631 |
| Kif26b | 0,568757387 | 0,000274246 | 0,012948152 |
| Tspan11 | 0,57204677 | 0,000840673 | 0,039765619 |
| Rgs22 | 0,573515779 | 0,000176031 | 0,011426047 |
| Col8a2 | 0,576282858 | 0,000178677 | 0,012908217 |
| Gstcd | 0,583194762 | 0,000833843 | 0,027780469 |
| Wdr6 | 0,584266716 | 1,27E-05 | 0,001310354 |
| Rdh12 | 0,58443698 | 0,001769132 | 0,049728944 |
| Brip1os | 0,587292383 | 0,000460758 | 0,018726232 |
| Gfra1 | 0,588655341 | 6,63E-06 | 0,000677525 |
| Il6ra | 0,591736029 | 0,001117703 | 0,033973911 |
| Gm42031 | 0,59407551 | 0,000937994 | 0,031115041 |
| Lck | 0,595412175 | 0,001071628 | 0,034007784 |
| Ptpro | 0,599567884 | 1,91E-05 | 0,001510626 |
| Igsf3 | 0,600791627 | 0,000150691 | 0,007704456 |
| Ece2 | 0,604581718 | 6,35E-06 | 0,000586788 |
| Fut10 | 0,607551693 | 0,00155736 | 0,034942827 |
| Ano2 | 0,615161076 | 0,000203252 | 0,01139279 |
| Ak7 | 0,616116489 | 0,001263943 | 0,04030276 |
| Ebf1 | 0,616438278 | 0,00083552 | 0,031119914 |
| 2410004P03Rik | 0,619676901 | 0,001257585 | 0,041697477 |
| Pcdhga12 | 0,624096146 | 0,000899048 | 0,029087631 |
| Igsf1 | 0,628240073 | 0,000761743 | 0,026276365 |
| Pcdhga10 | 0,636483919 | 0,000965982 | 0,029938831 |
| Irf7 | 0,639930956 | 0,000822138 | 0,027634227 |
| Igsf10 | 0,642171561 | 0,000353415 | 0,015312889 |
| Ccdc87 | 0,649173706 | 0,00049523 | 0,018649543 |
| Dnah12 | 0,651204191 | 1,07E-05 | 0,002123275 |
| Hmgcs2 | 0,652083733 | 0,000242696 | 0,011426047 |
| Capsl | 0,655089847 | 0,000400505 | 0,016818478 |
| Sfmbt2 | 0,658131703 | 0,000336865 | 0,011727582 |
| Cfap126 | 0,658685107 | 5,96E-05 | 0,005635671 |
| Polr2a | 0,678861139 | 2,67E-06 | 0,0003448 |
| Pcdhga6 | 0,680298517 | 0,000441857 | 0,018152713 |
| Sncaip | 0,684588953 | 2,29E-05 | 0,00181838 |
| Pcdhga8 | 0,686392732 | 0,000347162 | 0,016486336 |
| C77080 | 0,690614778 | 2,59E-05 | 0,002002965 |
| Kctd12b | 0,69125125 | 9,01E-05 | 0,005382905 |
| Cyb5d1 | 0,696242273 | 2,96E-05 | 0,001724717 |
| Fam183b | 0,702578304 | 0,000179967 | 0,010324527 |
| Scml4 | 0,719332133 | 0,000147899 | 0,008065032 |
| Arhgap36 | 0,739356321 | 5,63E-05 | 0,003881726 |
| Gpr165 | 0,741264897 | 8,83E-05 | 0,00515414 |
| Faxc | 0,744760246 | 1,40E-05 | 0,001287748 |

| | | | |
|---|---|---|---|
| Pcdhgb2 | 0,745882489 | 9,00E-05 | 0,00552384 |
| Ifit1 | 0,748841522 | 8,06E-05 | 0,005534 |
| Pcdhgb7 | 0,750664726 | 0,000102315 | 0,004678877 |
| Pcdhgb5 | 0,752502887 | 9,54E-05 | 0,005677413 |
| Gm14094 | 0,804955735 | 1,49E-06 | 0,000597222 |
| CT571252.1 | 0,822883914 | 1,46E-05 | 0,003513929 |
| Wdr66 | 0,826546804 | 3,09E-06 | 0,000340628 |
| Foxj1 | 0,838057082 | 1,38E-05 | 0,000947345 |
| Kcnmb1 | 0,865404996 | 2,51E-06 | 0,000372388 |
| Pcdhga2 | 0,869989217 | 7,03E-06 | 0,000677525 |
| Ccdc180 | 0,874189921 | 2,58E-08 | 1,81E-05 |
| Pcdhgb1 | 0,883698369 | 4,58E-06 | 0,000476199 |
| Fam196b | 0,905024825 | 4,42E-08 | 6,91E-06 |
| Ankfn1 | 0,90703772 | 7,79E-07 | 0,000319554 |
| Prkg1 | 0,925804886 | 1,54E-06 | 0,000194789 |
| Pcdhga11 | 0,929421574 | 1,27E-06 | 0,000165157 |
| Enkur | 0,942210706 | 1,11E-06 | 0,000155386 |
| Pcdhga9 | 0,944030269 | 1,02E-06 | 0,000151364 |
| Gm8420 | 0,958831318 | 2,16E-08 | 5,74E-06 |
| Gm5620 | 0,979323579 | 4,03E-07 | 6,30E-05 |
| Pcdhgb6 | 1,014673571 | 1,63E-07 | 3,14E-05 |
| Gm43305 | 1,01805116 | 1,42E-07 | 2,58E-05 |
| Slitrk6 | 1,045001988 | 3,95E-08 | 6,43E-06 |
| Gm10704 | 1,062292023 | 3,51E-08 | 2,21E-05 |
| Rsph4a | 1,094090703 | 3,78E-09 | 1,11E-06 |
| AW551984 | 1,239996513 | 9,43E-12 | 7,37E-09 |
| Gm6682 | 1,668972472 | 1,77E-18 | 2,74E-15 |

**Table S1. DEGs in the Nac of 8 weeks-old males**
**Comparison (p<0.05 cut-off) : *R6/1 $^{Tg/0}$ ; Rarβ $^{+/-}$ vs. R6/1 $^{Tg/0}$ ; Rarβ $^{+/+}$***

**Down regulated**

| Gene Name | Log2 Fold Change | p-value | adjusted p-value |
|---|---|---|---|
| Gm1840 | -1,223826435 | 3,69E-10 | 3,96E-06 |
| Gm5148 | -1,021025591 | 1,27E-07 | 0,000193921 |
| Rreb1 | -0,901130856 | 4,71E-08 | 6,25E-05 |
| Cd209c | -0,790855326 | 4,83E-05 | 0,010216348 |
| C1ql3 | -0,780354562 | 1,40E-06 | 0,000351097 |
| Gm23935 | -0,775964848 | 1,36E-10 | 3,15E-07 |
| Flywch2 | -0,763957663 | 1,01E-07 | 9,87E-05 |
| Gm26917 | -0,761986647 | 3,56E-06 | 0,000984653 |
| Col11a1 | -0,758329673 | 1,78E-05 | 0,003399563 |
| Stard9 | -0,747999591 | 6,48E-08 | 0,000162625 |
| Cd300lg | -0,728750216 | 0,000169349 | 0,044421442 |
| 4930426D05Rik | -0,72634494 | 0,000110931 | 0,035827903 |
| Gm3510 | -0,715052127 | 0,00020779 | 0,025208055 |
| Crybg2 | -0,713552191 | 0,00016761 | 0,022657897 |
| Pvalb | -0,707138599 | 1,21E-05 | 0,002193561 |
| Abi3bp | -0,692302038 | 1,67E-05 | 0,001942352 |
| 4930452B06Rik | -0,687652194 | 9,63E-07 | 0,000402512 |
| Aldh1a1 | -0,68554836 | 6,68E-07 | 0,000291092 |
| Igfbp6 | -0,670281136 | 5,05E-05 | 0,012811244 |
| Npas4 | -0,654036021 | 1,05E-05 | 0,001971097 |
| Igfbp2 | -0,651672722 | 2,23E-05 | 0,003325498 |
| Car7 | -0,64548402 | 5,11E-06 | 0,001531168 |
| Gm42418 | -0,64510931 | 2,32E-06 | 0,000351097 |
| Nrn1 | -0,639932599 | 2,52E-08 | 3,13E-05 |
| Nenf | -0,625996386 | 9,92E-06 | 0,001293435 |
| H2afj | -0,623600664 | 9,53E-06 | 0,001943008 |
| Cck | -0,622214457 | 3,19E-05 | 0,002467896 |
| Adcyap1 | -0,621545351 | 4,44E-05 | 0,012084987 |
| Pdzd7 | -0,606844833 | 0,000340185 | 0,042651954 |
| Nr4a1 | -0,590070586 | 3,75E-05 | 0,003757189 |
| Mir6240 | -0,589737323 | 7,17E-05 | 0,01566174 |
| Nktr | -0,589543831 | 1,70E-06 | 0,000299043 |
| Gng13 | -0,589353748 | 0,000149715 | 0,011602402 |
| Coro6 | -0,587509376 | 0,000346252 | 0,025448881 |
| Trf | -0,576049777 | 0,000243476 | 0,015773777 |
| Lars2 | -0,571709621 | 1,02E-05 | 0,001215699 |
| Polr2h | -0,567565182 | 5,54E-05 | 0,007558962 |
| Cryab | -0,566391853 | 1,82E-05 | 0,002321177 |
| C1qtnf4 | -0,565481188 | 2,85E-07 | 0,000110631 |
| Ttc9b | -0,562377893 | 6,27E-05 | 0,004306448 |
| Ethe1 | -0,559930062 | 0,000356021 | 0,044838632 |
| Uqcr10 | -0,557569166 | 1,53E-05 | 0,001781881 |
| Flnb | -0,557105489 | 1,17E-05 | 0,001836533 |
| Slc30a3 | -0,554040751 | 0,000372853 | 0,01929227 |
| Cox8a | -0,553493288 | 0,000145969 | 0,00765017 |
| Sec11c | -0,551531443 | 3,24E-07 | 0,000162625 |
| Tmem160 | -0,545333638 | 1,77E-05 | 0,003399563 |
| Chuk | -0,543775272 | 0,000603642 | 0,021620875 |
| Cabp1 | -0,540377793 | 1,16E-06 | 0,000299043 |
| Zfp950 | -0,537385055 | 0,001021871 | 0,024969045 |
| Necab3 | -0,533977487 | 8,36E-06 | 0,001435019 |
| Ppfibp1 | -0,53198153 | 0,000123269 | 0,013691628 |
| Gm5436 | -0,528772982 | 0,000124051 | 0,04094583 |
| Ndufa1 | -0,525857331 | 2,21E-06 | 0,00038527 |

| | | | |
|---|---|---|---|
| Ndufa13 | -0,522593271 | 5,64E-05 | 0,003586594 |
| Eml6 | -0,521975023 | 3,98E-05 | 0,00511662 |
| Atp5j2 | -0,521825556 | 0,000435189 | 0,014886974 |
| Abca7 | -0,521515577 | 0,000172066 | 0,012579742 |
| Mgst3 | -0,521488571 | 1,30E-05 | 0,001888324 |
| Uqcr11 | -0,520137652 | 1,43E-05 | 0,001531168 |
| Ndufa6 | -0,519329432 | 1,37E-06 | 0,00028814 |
| Fabp3 | -0,517161371 | 3,20E-05 | 0,003150312 |
| Akap13 | -0,516547169 | 0,000224228 | 0,012085485 |
| Atp5k | -0,515744661 | 0,000495448 | 0,018538257 |
| Ndufa3 | -0,515710911 | 0,000305956 | 0,012084987 |
| Polr2f | -0,515710613 | 0,000321053 | 0,020908861 |
| Ndufb7 | -0,51221314 | 0,000313488 | 0,011436928 |
| Chchd10 | -0,511702874 | 6,78E-05 | 0,00441316 |
| Uqcrq | -0,510502346 | 8,89E-05 | 0,00687265 |
| Cox6a1 | -0,509109242 | 0,00015357 | 0,00742772 |
| Cox6c | -0,508242699 | 5,41E-05 | 0,003533712 |
| Lsm7 | -0,508223627 | 0,000249735 | 0,01566174 |
| Mecr | -0,506441589 | 0,000153937 | 0,013766469 |
| D930016D06Ril | -0,506212126 | 0,001139226 | 0,048007361 |
| AC121965.1 | -0,503713004 | 0,000331169 | 0,014796494 |
| Adamts10 | -0,503477363 | 7,48E-05 | 0,005108536 |
| Tmem256 | -0,499594395 | 0,000357064 | 0,015740311 |
| Ndufa11 | -0,49742956 | 0,000176607 | 0,009678465 |
| Ndufs6 | -0,496683809 | 0,000110732 | 0,00788448 |
| Fhl2 | -0,495706073 | 0,000139915 | 0,011294757 |
| Ndst4 | -0,494154809 | 0,001392024 | 0,037623476 |
| Tsr3 | -0,493358141 | 0,000124198 | 0,010184581 |
| Atp5e | -0,49303512 | 0,000409513 | 0,016682025 |
| Ndufb9 | -0,492596157 | 2,84E-05 | 0,002321177 |
| Myo9b | -0,491444618 | 3,45E-05 | 0,003325498 |
| Atp11b | -0,489684245 | 0,002141996 | 0,04896184 |
| Fkbp1b | -0,487668745 | 0,000198414 | 0,015740311 |
| Cuta | -0,487433767 | 0,000186645 | 0,01314944 |
| Hbb-bs | -0,487073792 | 0,00048498 | 0,023128816 |
| Cisd3 | -0,485583357 | 0,000561407 | 0,017621249 |
| Rnaseh2c | -0,485248542 | 0,00125215 | 0,043389486 |
| Atf4 | -0,482598272 | 3,47E-06 | 0,000485062 |
| 2010107E04Rik | -0,482440853 | 0,000272503 | 0,010184581 |
| Fam193b | -0,482237183 | 7,83E-05 | 0,004670791 |
| Cox7b | -0,479006399 | 5,92E-05 | 0,003728188 |
| Cttnbp2 | -0,478592893 | 7,65E-05 | 0,006044535 |
| Mir6236 | -0,477149834 | 0,000399643 | 0,013034936 |
| Srsf11 | -0,475121212 | 0,001852153 | 0,037484267 |
| Mapk11 | -0,473800492 | 0,000209759 | 0,013994165 |
| Enho | -0,47179349 | 0,000221943 | 0,012066438 |
| Rprml | -0,468473024 | 0,00018201 | 0,009918862 |
| Scand1 | -0,465059803 | 0,000499621 | 0,019247979 |
| Ccdc85b | -0,463993432 | 2,27E-05 | 0,001971097 |
| Smg1 | -0,462954263 | 0,001543232 | 0,030049251 |
| 1110065P20Rik | -0,456655387 | 0,0005885 | 0,0212825 |
| Ndufb2 | -0,454972202 | 0,000103929 | 0,007558962 |
| Fmc1 | -0,454147372 | 0,002477974 | 0,046179182 |
| Cnr1 | -0,453961986 | 0,00212134 | 0,037029526 |
| Fahd2a | -0,453632805 | 0,000324613 | 0,018742842 |
| Arhgdig | -0,452970836 | 0,000258654 | 0,013034936 |
| Cox7a2 | -0,452816237 | 0,000155042 | 0,00788448 |
| Dgcr6 | -0,451943519 | 0,000557231 | 0,02022997 |
| Selenom | -0,451361233 | 0,000481874 | 0,01822353 |

| | | | |
|---|---|---|---|
| Dynll1 | -0,451261651 | 0,000287736 | 0,011436928 |
| Hddc3 | -0,449514775 | 0,000965055 | 0,043598725 |
| Ndufb10 | -0,449177717 | 0,000287832 | 0,011676595 |
| 2010111I01Rik | -0,446834789 | 0,000390963 | 0,019891037 |
| Mrpl28 | -0,443254052 | 0,000173667 | 0,010356123 |
| Yeats2 | -0,443244812 | 0,000626331 | 0,027976154 |
| Acyp2 | -0,443160926 | 0,000221229 | 0,016872946 |
| Nme1 | -0,442917122 | 3,95E-05 | 0,002914426 |
| Edf1 | -0,44283797 | 0,000900023 | 0,022851974 |
| Lamtor4 | -0,440914483 | 0,00052859 | 0,016570401 |
| Mpc1 | -0,440224024 | 1,69E-05 | 0,00215233 |
| Lrrc45 | -0,439926996 | 0,000326549 | 0,015016547 |
| Rsrp1 | -0,43876052 | 0,002792558 | 0,043886569 |
| Ndufa2 | -0,438733497 | 0,000424153 | 0,01475641 |
| Mif | -0,437394778 | 0,00070322 | 0,019478344 |
| Mrps7 | -0,436793071 | 0,00012653 | 0,007206722 |
| Fras1 | -0,436657947 | 0,000580631 | 0,02022997 |
| Nhp2 | -0,436120767 | 0,000467578 | 0,018299455 |
| Mrpl41 | -0,43603148 | 0,000488362 | 0,019010211 |
| Ndufb11 | -0,434787727 | 0,000467583 | 0,014131934 |
| Mrpl20 | -0,433479009 | 0,000171642 | 0,010184581 |
| Lamtor2 | -0,433252126 | 0,001311761 | 0,037029526 |
| Pcp4 | -0,430827268 | 0,000257433 | 0,010078663 |
| Kctd6 | -0,430781431 | 0,000294818 | 0,017178525 |
| Exosc9 | -0,426841647 | 0,000268825 | 0,016872946 |
| Jtb | -0,425680789 | 0,000575952 | 0,027631378 |
| Atp5d | -0,425579703 | 7,61E-05 | 0,004799894 |
| Rplp2 | -0,425425268 | 0,001780301 | 0,036658238 |
| Timm13 | -0,425104507 | 0,000190464 | 0,010614907 |
| Vwa5b2 | -0,425101566 | 0,001192482 | 0,034543034 |
| Smdt1 | -0,42468476 | 0,001917186 | 0,035827903 |
| Plp1 | -0,423322927 | 0,000781699 | 0,019247979 |
| Ndufs7 | -0,422857209 | 0,002163486 | 0,037483394 |
| Psmb5 | -0,422676066 | 0,000667917 | 0,021085462 |
| Selenow | -0,420745049 | 0,000300864 | 0,010971054 |
| Uqcrb | -0,420630106 | 0,000439072 | 0,017621249 |
| Cox5a | -0,420572836 | 6,92E-05 | 0,004119416 |
| Mpdz | -0,420528055 | 0,000432261 | 0,014796494 |
| Ndufb8 | -0,419989017 | 0,000182727 | 0,0080887 |
| Pmvk | -0,419493598 | 0,000545866 | 0,02022997 |
| Ddit3 | -0,418687713 | 0,000988929 | 0,044350192 |
| Cox5b | -0,417870015 | 0,000390951 | 0,013994165 |
| Kcnab3 | -0,417640342 | 0,001923511 | 0,045673148 |
| Atg16l2 | -0,416800115 | 0,001204886 | 0,042375378 |
| Ptprd | -0,416702356 | 0,00264313 | 0,046672778 |
| Cox4i1 | -0,414314804 | 0,000574788 | 0,016998206 |
| Fkbp2 | -0,412880131 | 0,000902395 | 0,027697142 |
| Trank1 | -0,411548471 | 0,00072394 | 0,018072117 |
| Pop5 | -0,409823 | 0,000537507 | 0,019891037 |
| Mrpl46 | -0,407495482 | 0,000359024 | 0,022657897 |
| Commd4 | -0,407354018 | 0,000249245 | 0,012811244 |
| Kcnt1 | -0,406550503 | 0,000904973 | 0,02068253 |
| Smpd4 | -0,406026027 | 0,000337412 | 0,01424844 |
| Cox6b1 | -0,405987673 | 0,001481605 | 0,029352643 |
| Mrpl36 | -0,405693208 | 0,001153096 | 0,04096405 |
| Mdn1 | -0,405524454 | 0,00158549 | 0,041508952 |
| Matk | -0,404256181 | 4,60E-05 | 0,003533712 |
| Pkd1 | -0,401740628 | 0,000415927 | 0,013349771 |
| Mrpl13 | -0,398932773 | 0,000663257 | 0,018756419 |

| | | | |
|---|---|---|---|
| Ndufs5 | -0,398549056 | 0,002264012 | 0,047466673 |
| Ndufa8 | -0,395573116 | 0,000703787 | 0,023038801 |
| Ndufa4 | -0,395548291 | 0,000227412 | 0,009259447 |
| Hint1 | -0,39378022 | 0,000341968 | 0,012084987 |
| Unc13b | -0,393350329 | 0,00174975 | 0,04340421 |
| Ap2s1 | -0,393018418 | 0,000941108 | 0,023395863 |
| Pcsk1n | -0,392244103 | 0,000595767 | 0,017254465 |
| Zfp428 | -0,390328928 | 0,001083161 | 0,032468292 |
| Atp5o | -0,389941405 | 0,000531923 | 0,016682025 |
| Atpif1 | -0,388660576 | 0,00184326 | 0,033922868 |
| Dmxl2 | -0,387111126 | 0,002330951 | 0,039500464 |
| Atp5g3 | -0,386893005 | 0,000186273 | 0,0080887 |
| Cpne7 | -0,38613134 | 0,003474661 | 0,04940765 |
| Ppp1r1a | -0,384987625 | 0,000998558 | 0,030528087 |
| Higd2a | -0,384022048 | 0,000970007 | 0,030049251 |
| Mrps34 | -0,383413861 | 0,001570287 | 0,040970578 |
| Tpgs1 | -0,380059935 | 0,002129446 | 0,04896184 |
| Uqcrfs1 | -0,379888118 | 9,00E-05 | 0,00511662 |
| Rps6kb2 | -0,379543637 | 0,002207714 | 0,049776491 |
| Rnf112 | -0,377579471 | 0,000109865 | 0,005930544 |
| Tmsb4x | -0,374686496 | 0,000794802 | 0,01929227 |
| Uqcrh | -0,373139519 | 0,001699465 | 0,031923461 |
| Stmn3 | -0,373072138 | 0,00031105 | 0,011436928 |
| Pfdn5 | -0,372945415 | 0,002892417 | 0,04896184 |
| Fis1 | -0,371916526 | 0,000653815 | 0,017254465 |
| Tbcb | -0,369737029 | 0,002068029 | 0,037483394 |
| Fam173a | -0,369719166 | 0,00134248 | 0,037483394 |
| Szt2 | -0,368810857 | 0,001229747 | 0,032247202 |
| Prelid1 | -0,368495481 | 0,002064348 | 0,036491998 |
| Nrgn | -0,368409111 | 0,002573431 | 0,041877388 |
| Mrps26 | -0,367164701 | 0,001214128 | 0,031923461 |
| Ubc | -0,366789786 | 0,001756679 | 0,042651954 |
| Nkiras1 | -0,366368369 | 0,000385869 | 0,012969619 |
| Tspoap1 | -0,363985138 | 0,002859579 | 0,044357455 |
| Celsr3 | -0,363753992 | 0,001191292 | 0,02526324 |
| Per2 | -0,363401297 | 0,00215078 | 0,041026465 |
| Camk2n2 | -0,362830097 | 0,000324893 | 0,011436928 |
| Tmem234 | -0,362402964 | 0,000240807 | 0,012200548 |
| 2700060E02Rik | -0,361760623 | 0,002126818 | 0,048704578 |
| Coa3 | -0,36038406 | 0,001803617 | 0,04340421 |
| Psmb4 | -0,357866267 | 0,001509024 | 0,029689556 |
| Scn1b | -0,354694048 | 0,000128401 | 0,007110733 |
| Gapdh | -0,350592476 | 0,003488998 | 0,04968483 |
| Emc4 | -0,350543654 | 0,001652056 | 0,031206674 |
| Haghl | -0,34624049 | 0,002690015 | 0,042692465 |
| Qdpr | -0,342843946 | 0,001436113 | 0,030811145 |
| Ndufc1 | -0,342501232 | 0,001731805 | 0,042375378 |
| Abcc5 | -0,342477012 | 0,002043962 | 0,036372211 |
| Gls | -0,341955073 | 0,001896043 | 0,035091144 |
| Hagh | -0,3387 | 0,00215789 | 0,048007361 |
| Pebp1 | -0,337105307 | 0,000541524 | 0,016872946 |
| Pmm1 | -0,335864939 | 0,002104532 | 0,040955733 |
| Atp5j | -0,335403546 | 0,003007286 | 0,049739462 |
| Ice1 | -0,335072394 | 0,002781 | 0,048007361 |
| Rtca | -0,332194868 | 0,002139521 | 0,04896184 |
| Myl12b | -0,331980486 | 0,001375422 | 0,030811145 |
| Dot1l | -0,330665299 | 0,002286299 | 0,04940765 |
| Eif1 | -0,329545149 | 0,001775198 | 0,035815718 |
| Gpatch8 | -0,327761524 | 0,001403901 | 0,031206674 |

| | | | |
|---|---|---|---|
| Mpc2 | -0,3253973 | 0,002338557 | 0,048007361 |
| Cdk10 | -0,325143686 | 0,002078306 | 0,044838632 |
| Arhgef25 | -0,322494112 | 0,001435589 | 0,030811145 |
| Anxa5 | -0,320604477 | 0,0021918 | 0,04968483 |
| Ppia | -0,319327416 | 0,001902248 | 0,038017203 |
| Mycbp2 | -0,318531344 | 0,000777897 | 0,018877798 |
| Arf5 | -0,313793459 | 0,00250087 | 0,044838632 |
| Gpx4 | -0,313425365 | 0,001907576 | 0,0447881 |
| Tmem191c | -0,308593754 | 0,001931853 | 0,035039171 |
| Snrpn | -0,30486673 | 0,000941217 | 0,022851974 |
| Sncb | -0,302986116 | 0,0023581 | 0,04340421 |
| Slc25a4 | -0,300226935 | 0,000937046 | 0,021468942 |
| Cend1 | -0,292557821 | 0,001530744 | 0,033108117 |
| Rab3a | -0,287068509 | 0,002275747 | 0,041609751 |
| Bsg | -0,284493457 | 0,002929911 | 0,044838632 |
| Bex2 | -0,276444124 | 0,003038398 | 0,04896184 |

**Table S1. DEGs in the Nac of 8 weeks-old males**
**Comparison (p<0.05 cut-off) : *R6/1 $^{Tg/0}$ ; Rarβ $^{+/-}$  vs. R6/1 $^{Tg/0}$ ; Rarβ $^{+/+}$***

**Up regulated**

| Gene Name | Log2 Fold Change | p-value | adjusted p-value |
|---|---|---|---|
| Slc1a2 | 0,272504471 | 0,002968038 | 0,045320379 |
| Gng7 | 0,276230416 | 0,002543569 | 0,044350192 |
| Scd2 | 0,285044953 | 0,003105025 | 0,047466673 |
| Efnb3 | 0,295461452 | 0,002476528 | 0,040970578 |
| Ski | 0,299599302 | 0,003242189 | 0,047943339 |
| Rbfox2 | 0,303444595 | 0,002331411 | 0,042330113 |
| Slc1a3 | 0,303701609 | 0,000604776 | 0,017802784 |
| Gatad2b | 0,304840404 | 0,002126478 | 0,037932154 |
| Ddah1 | 0,315636584 | 0,003013723 | 0,049776491 |
| Slc38a1 | 0,318821187 | 0,003138912 | 0,047309751 |
| Cmtm4 | 0,321637194 | 0,003013449 | 0,045626468 |
| Scg3 | 0,323612169 | 0,002970804 | 0,045276066 |
| Copb2 | 0,324748872 | 0,001813404 | 0,04340421 |
| Acsl3 | 0,327251683 | 0,000669685 | 0,017254465 |
| D430019H16Ril | 0,346471905 | 0,002230167 | 0,042330113 |
| Ckap5 | 0,347534391 | 0,001168451 | 0,024660813 |
| Tex2 | 0,348139626 | 0,001678423 | 0,0324946 |
| Itsn1 | 0,357190763 | 0,000727526 | 0,018598225 |
| Caln1 | 0,361783468 | 0,002317342 | 0,039980684 |
| Rgs8 | 0,361871557 | 0,000708339 | 0,018299455 |
| Slc32a1 | 0,362283138 | 0,001097549 | 0,023770494 |
| Nr2c2 | 0,363135672 | 0,001159973 | 0,02521404 |
| Prkacb | 0,363183023 | 0,000160902 | 0,0080887 |
| Sec14l1 | 0,363341481 | 0,000846947 | 0,019891037 |
| Sox8 | 0,366019823 | 0,001554507 | 0,040955733 |
| Stox2 | 0,36710755 | 0,00057789 | 0,02022997 |
| Wdr6 | 0,370086268 | 0,002350188 | 0,042375378 |
| Timp3 | 0,371206053 | 0,000380544 | 0,013766469 |
| Lsamp | 0,372342926 | 0,000201964 | 0,009356945 |
| Socs7 | 0,373861699 | 0,003220796 | 0,047644298 |
| Zmiz1 | 0,373946104 | 0,000330136 | 0,011865414 |
| Prepl | 0,379883664 | 0,000869553 | 0,020314522 |
| Bcl9 | 0,383184709 | 0,000245357 | 0,012313266 |
| Tmem164 | 0,383407272 | 0,000887009 | 0,022657897 |
| Fbxo10 | 0,383744921 | 0,000692041 | 0,019247979 |
| Adcy5 | 0,38515638 | 0,000107241 | 0,006429745 |
| Mon1b | 0,386240945 | 0,002456629 | 0,044350192 |
| Foxk1 | 0,390884318 | 0,002446923 | 0,049012932 |
| Dlg2 | 0,391269459 | 0,000220284 | 0,009008995 |
| Rcan3 | 0,39590806 | 0,001989111 | 0,047466673 |
| Rragd | 0,396612379 | 0,000493703 | 0,015828923 |
| Msi2 | 0,397758442 | 0,002738941 | 0,047644298 |
| Rgs20 | 0,400508127 | 0,001103585 | 0,040550679 |
| Bhlhe41 | 0,403056064 | 0,001398228 | 0,038127878 |
| Lima1 | 0,403253522 | 0,001903553 | 0,039779495 |
| Sin3a | 0,403989468 | 0,000356415 | 0,01566174 |
| Ntrk2 | 0,404033326 | 2,72E-05 | 0,002415667 |
| Lrrc8d | 0,404817066 | 0,001037864 | 0,031206674 |
| Doc2b | 0,407188537 | 0,002910502 | 0,04896184 |
| Rrm1 | 0,414166048 | 0,002800165 | 0,049403218 |
| Gad2 | 0,414367046 | 0,000163195 | 0,00765017 |
| Tnfrsf19 | 0,414906915 | 0,000106235 | 0,007650515 |
| Midn | 0,417081508 | 0,002070073 | 0,048401098 |
| Syndig1l | 0,418595706 | 4,47E-05 | 0,003855695 |

| | | | |
|---|---|---|---|
| Tshz1 | 0,421094632 | 0,000665026 | 0,022851974 |
| Dclk3 | 0,42138389 | 0,000359175 | 0,012084987 |
| Hap1 | 0,424405576 | 0,000130855 | 0,00742772 |
| Tcf20 | 0,424914297 | 0,001990095 | 0,035815718 |
| Gm5607 | 0,426543715 | 0,001175242 | 0,027631378 |
| Erf | 0,427110579 | 0,000922909 | 0,028781863 |
| Amotl1 | 0,427967111 | 0,000374087 | 0,015995577 |
| Rims3 | 0,42826141 | 0,000140358 | 0,00765017 |
| Atp2b4 | 0,430184013 | 0,002142454 | 0,04110512 |
| Hrasls | 0,434446942 | 0,00050991 | 0,025208055 |
| Clic4 | 0,435270312 | 0,00035477 | 0,01566174 |
| Arl4c | 0,436255352 | 0,000119791 | 0,0080887 |
| 2310022B05Rik | 0,43782764 | 0,000212098 | 0,027631378 |
| Plpp3 | 0,442091311 | 0,000158814 | 0,007558962 |
| Dpysl3 | 0,442553416 | 0,000182718 | 0,0080887 |
| Bahcc1 | 0,442828953 | 0,000808469 | 0,032709062 |
| Wfs1 | 0,444203945 | 0,000453166 | 0,015269157 |
| Sp1 | 0,444853711 | 0,000318842 | 0,017488691 |
| Ccdc153 | 0,447495443 | 0,00028046 | 0,031040988 |
| Ldb2 | 0,450609584 | 0,000310901 | 0,01466359 |
| Ptch1 | 0,451207883 | 1,53E-05 | 0,002063053 |
| Senp5 | 0,451477613 | 0,001339551 | 0,044838632 |
| Dock10 | 0,453975903 | 0,00035443 | 0,01566174 |
| Ccdc85c | 0,454150076 | 5,64E-05 | 0,003586594 |
| Nr1d1 | 0,454614394 | 0,000167878 | 0,00788448 |
| Ptgfrn | 0,454843072 | 0,000354372 | 0,022565933 |
| Meis2 | 0,458419961 | 3,08E-05 | 0,002424017 |
| 1700024G13Rik | 0,458996642 | 9,77E-05 | 0,034839902 |
| Baiap3 | 0,459149634 | 0,00025733 | 0,009957001 |
| Chst11 | 0,461224881 | 3,48E-05 | 0,003325498 |
| Erich3 | 0,461499665 | 2,75E-05 | 0,003093506 |
| Kcnj10 | 0,467008911 | 0,00020618 | 0,008744734 |
| Sparc | 0,467499839 | 1,08E-05 | 0,001369852 |
| Sox9 | 0,467991677 | 0,000322891 | 0,0177967 |
| Zfp609 | 0,469138267 | 7,32E-05 | 0,006044535 |
| Myo10 | 0,470384482 | 3,57E-05 | 0,003586594 |
| 1700001C02Rik | 0,473549929 | 0,00012625 | 0,04096405 |
| Akap5 | 0,476767615 | 0,000166507 | 0,008250417 |
| Grid2 | 0,47690647 | 0,000203124 | 0,014407484 |
| Sox2 | 0,480519607 | 0,001183728 | 0,034739718 |
| Gfra1 | 0,482335106 | 4,69E-05 | 0,003977882 |
| Pou3f3 | 0,482586383 | 5,73E-06 | 0,00104553 |
| Crkl | 0,485417142 | 0,000841978 | 0,045435423 |
| Grk5 | 0,486377008 | 0,000629361 | 0,027561865 |
| Reln | 0,487321209 | 3,87E-05 | 0,004799894 |
| Pard3 | 0,488687816 | 0,000575601 | 0,036369913 |
| Lrrc36 | 0,491467378 | 0,000167157 | 0,033818137 |
| Lrrc71 | 0,492864886 | 5,80E-05 | 0,017488691 |
| Pcdh9 | 0,492983924 | 0,000347563 | 0,01566174 |
| Tacr1 | 0,496591592 | 0,000104644 | 0,00914864 |
| Ankrd52 | 0,496653944 | 0,000305166 | 0,01445485 |
| Papss2 | 0,49863059 | 0,000228988 | 0,010431373 |
| Nos1 | 0,506217399 | 0,000383193 | 0,040955733 |
| 2010001K21Rik | 0,506638358 | 5,61E-05 | 0,025995514 |
| Ston2 | 0,507559626 | 0,000435654 | 0,025854096 |
| Hmgn2 | 0,508398911 | 0,000139594 | 0,008746569 |
| Notch1 | 0,508424959 | 7,05E-05 | 0,007650515 |
| Glg1 | 0,509644852 | 0,000230988 | 0,00996347 |
| Pbx3 | 0,511506829 | 1,12E-05 | 0,001781881 |

| | | | |
|---|---|---|---|
| Ankrd63 | 0,514882402 | 8,02E-07 | 0,000274066 |
| Arx | 0,516399601 | 0,000411518 | 0,028956963 |
| Cachd1 | 0,523806635 | 8,54E-05 | 0,0080887 |
| Dpy19l3 | 0,529381235 | 2,18E-06 | 0,000416417 |
| Ptprt | 0,53196785 | 9,70E-06 | 0,001193705 |
| Sox11 | 0,532991611 | 0,001097913 | 0,030049251 |
| Dusp18 | 0,538219634 | 0,000123244 | 0,010184581 |
| Camk1g | 0,539754246 | 0,000381982 | 0,023395863 |
| Ndst1 | 0,544193422 | 0,000247142 | 0,012805618 |
| Ednrb | 0,549413364 | 0,000334704 | 0,015269157 |
| Itih3 | 0,553598366 | 0,000206513 | 0,016118307 |
| Gja1 | 0,55453851 | 5,51E-08 | 4,30E-05 |
| Epb41l5 | 0,554637994 | 0,000387649 | 0,047466673 |
| Ano2 | 0,554692853 | 0,00033676 | 0,021620875 |
| Id4 | 0,556484349 | 1,22E-05 | 0,001369852 |
| Nr2e1 | 0,55666315 | 8,17E-05 | 0,017178525 |
| Stom | 0,558301235 | 0,000180448 | 0,028222732 |
| Nme9 | 0,564840061 | 6,71E-05 | 0,027090499 |
| Nedd9 | 0,565284233 | 0,000265523 | 0,036658238 |
| Sema3f | 0,568020991 | 5,12E-05 | 0,005834051 |
| Map3k13 | 0,58436363 | 1,55E-05 | 0,002588586 |
| Rnd3 | 0,58571188 | 0,000926955 | 0,042651954 |
| Sox4 | 0,599155558 | 6,10E-05 | 0,005301459 |
| Fads2 | 0,608096749 | 0,000121621 | 0,0080887 |
| Fndc9 | 0,609675135 | 8,24E-05 | 0,017488691 |
| Gpx3 | 0,614515968 | 0,000311148 | 0,040925266 |
| Igfbp5 | 0,614852169 | 1,83E-05 | 0,001946821 |
| Fgd5 | 0,615707899 | 0,000192889 | 0,03005987 |
| Cfap52 | 0,622752041 | 9,77E-05 | 0,035843158 |
| Ccnd2 | 0,632918031 | 1,94E-05 | 0,001971097 |
| Gli3 | 0,63812216 | 0,000112551 | 0,020754548 |
| Pou3f4 | 0,649763552 | 3,36E-05 | 0,004306448 |
| Gm6123 | 0,650169848 | 4,30E-05 | 0,014811807 |
| Hdc | 0,651373194 | 0,000165735 | 0,04896184 |
| Spata13 | 0,654340899 | 3,50E-07 | 0,000170698 |
| Tk1 | 0,655211053 | 0,000156362 | 0,048007361 |
| Cfap126 | 0,658009269 | 0,000162732 | 0,04896184 |
| Mex3a | 0,659839561 | 0,000194551 | 0,030234392 |
| Gm45396 | 0,661631211 | 0,000242478 | 0,028148257 |
| Tead1 | 0,668232133 | 5,95E-06 | 0,00297221 |
| Zic1 | 0,668804681 | 3,90E-06 | 0,00104553 |
| Tmem212 | 0,669856302 | 3,40E-06 | 0,004020489 |
| Fam84b | 0,672672382 | 0,000126439 | 0,022851974 |
| Dlx2 | 0,676144973 | 9,39E-05 | 0,006044535 |
| Pcdhb6 | 0,686271025 | 0,00023754 | 0,035843158 |
| Dnah12 | 0,688825579 | 1,45E-05 | 0,011436928 |
| Pcdhgb4 | 0,691100103 | 0,000369953 | 0,03878901 |
| Vat1l | 0,69122465 | 5,08E-09 | 7,38E-06 |
| Emp2 | 0,692587215 | 1,39E-06 | 0,000416417 |
| Mis18bp1 | 0,696028356 | 0,000169105 | 0,037837471 |
| Ckap2 | 0,697263574 | 0,000231448 | 0,037029526 |
| Arhgap36 | 0,69966356 | 7,15E-05 | 0,01566174 |
| Dlec1 | 0,70154372 | 0,000275938 | 0,030327748 |
| Gpr165 | 0,70456885 | 0,000128295 | 0,037484267 |
| Kif20a | 0,707959246 | 0,000188658 | 0,023038801 |
| Dmkn | 0,710021051 | 0,000244929 | 0,027976154 |
| Cd24a | 0,712754366 | 2,23E-05 | 0,003360368 |
| Pou3f1 | 0,7132691 | 4,87E-07 | 0,000299043 |
| Dlk1 | 0,714889425 | 0,000245372 | 0,009655238 |

| | | | |
|---|---|---|---|
| Pde7b | 0,718633287 | 1,92E-08 | 2,55E-05 |
| Iqca | 0,719599003 | 3,69E-05 | 0,013021182 |
| Ecel1 | 0,720944472 | 1,56E-07 | 0,000127305 |
| Gm7224 | 0,721213745 | 0,000198083 | 0,033442298 |
| Dgkk | 0,722455313 | 0,000157906 | 0,042651954 |
| Rorc | 0,723093684 | 0,000213101 | 0,032342927 |
| Rtl1 | 0,724301208 | 0,000163622 | 0,022400487 |
| Rsph1 | 0,729265349 | 0,000130491 | 0,023197185 |
| Tsku | 0,730677432 | 0,000184787 | 0,031923461 |
| Pcdhgc4 | 0,735164936 | 8,37E-05 | 0,028224765 |
| Nuf2 | 0,741307374 | 0,000115413 | 0,017846643 |
| Fzd5 | 0,745987599 | 2,24E-05 | 0,0080887 |
| Dnali1 | 0,746424159 | 6,60E-06 | 0,004799894 |
| Rgs22 | 0,747277372 | 4,88E-06 | 0,003888469 |
| Hydin | 0,747362074 | 0,000132453 | 0,025232373 |
| Mcm3 | 0,752253819 | 4,58E-05 | 0,012313266 |
| Myb | 0,75311497 | 9,55E-06 | 0,0080887 |
| Shisa2 | 0,75477051 | 4,94E-05 | 0,013171042 |
| Chodl | 0,758635323 | 8,82E-05 | 0,019286387 |
| Faxc | 0,758941583 | 2,57E-06 | 0,000738532 |
| Ckap2l | 0,759782369 | 9,60E-05 | 0,02022997 |
| Gm973 | 0,763114378 | 3,89E-05 | 0,0171994 |
| Bub1b | 0,763637372 | 7,03E-05 | 0,015740311 |
| Soga1 | 0,764049129 | 1,80E-05 | 0,002914426 |
| Slc18a2 | 0,772877688 | 3,97E-05 | 0,019478344 |
| Prdm16 | 0,774274485 | 2,18E-07 | 0,000299043 |
| Klf5 | 0,779142928 | 2,36E-06 | 0,001888324 |
| Cenpf | 0,78646821 | 4,00E-05 | 0,00914568 |
| Celsr1 | 0,790112549 | 1,79E-05 | 0,011436928 |
| Gm43305 | 0,791060628 | 3,71E-05 | 0,002799897 |
| Pcdhga2 | 0,800143671 | 3,95E-05 | 0,011436928 |
| Sstr5 | 0,804507221 | 2,63E-05 | 0,015498355 |
| Sncaip | 0,805122597 | 9,23E-08 | 0,000195804 |
| Hmgb2 | 0,806324206 | 2,71E-05 | 0,013994165 |
| Aspm | 0,807231622 | 2,33E-05 | 0,007110733 |
| 2410004P03Rik | 0,80997466 | 3,39E-05 | 0,008717991 |
| Dlx1 | 0,816143449 | 1,44E-07 | 9,87E-05 |
| Ttc21a | 0,817730069 | 2,17E-05 | 0,013766469 |
| Pcdhgb7 | 0,820337087 | 2,64E-05 | 0,009051115 |
| Pcdhgb1 | 0,82206536 | 2,57E-05 | 0,008717991 |
| Pcdhga9 | 0,822870812 | 1,68E-05 | 0,007110733 |
| Rsph4a | 0,827476923 | 3,62E-06 | 0,002358129 |
| Lmnb1 | 0,831715072 | 1,57E-07 | 0,00025156 |
| Gabre | 0,84460479 | 1,12E-05 | 0,009008995 |
| Cdhr4 | 0,845482302 | 1,72E-07 | 0,000351097 |
| Col8a2 | 0,845798509 | 2,64E-07 | 0,000721455 |
| Uhrf1 | 0,84791268 | 1,41E-05 | 0,004670791 |
| Fam167a | 0,851129644 | 1,14E-05 | 0,005617132 |
| Sfrp1 | 0,854802405 | 2,92E-08 | 9,87E-05 |
| Ascl1 | 0,85758199 | 7,88E-06 | 0,007110733 |
| Zfp521 | 0,876725341 | 3,13E-08 | 9,87E-05 |
| Drc7 | 0,881412241 | 6,44E-06 | 0,002914426 |
| Vwa5b1 | 0,882866823 | 4,03E-06 | 0,001971097 |
| Prelp | 0,888561311 | 1,62E-07 | 0,00028814 |
| AW551984 | 0,891880214 | 2,90E-07 | 0,000176569 |
| Kcnmb1 | 0,907539924 | 2,05E-06 | 0,003033 |
| Ebf1 | 0,907712109 | 3,46E-07 | 0,000416417 |
| Prc1 | 0,912581108 | 2,60E-06 | 0,001943008 |
| Pcdhgb5 | 0,915868893 | 2,76E-06 | 0,001605804 |

| | | | |
|---|---|---|---|
| Gm14094 | 0,939791349 | 1,12E-07 | 0,000351097 |
| Ak7 | 0,941801792 | 1,38E-06 | 0,001280532 |
| Pcdhga6 | 0,946885464 | 9,10E-07 | 0,000936311 |
| St8sia2 | 0,950885159 | 1,25E-07 | 0,000243898 |
| Hmgcs2 | 0,959260288 | 1,40E-08 | 6,25E-05 |
| Foxm1 | 0,987072496 | 1,29E-07 | 0,00025156 |
| Lrrc23 | 0,994530496 | 2,68E-07 | 0,000299043 |
| Rrm2 | 1,002963135 | 2,40E-07 | 0,000323446 |
| Gm10704 | 1,01562347 | 2,01E-07 | 0,000299043 |
| Gm5620 | 1,022443669 | 1,60E-07 | 0,000291092 |
| CT571252.1 | 1,034905366 | 1,05E-07 | 0,00017369 |
| Ccdc180 | 1,042235239 | 4,74E-10 | 1,08E-05 |
| Pcdhga11 | 1,056235864 | 2,18E-08 | 9,70E-05 |
| Pcdhgb6 | 1,056682781 | 5,43E-08 | 0,000160049 |
| Gm8420 | 1,087275478 | 1,82E-09 | 1,12E-05 |
| Foxj1 | 1,447623178 | 3,41E-14 | 2,02E-09 |
| Gm6682 | 1,844211808 | 3,09E-21 | 1,97E-16 |

**Table S1. DEGs in the Nac of 8 weeks-old males**
**Likelihood Ratio Analyses for synergy between R6/1 and *Rarβ***

**Down regulated**

| Gene Name | Log2 Fold Change | p-value | adjusted p-value |
|---|---|---|---|
| Gm6969 | -16,62900151 | 0,000101601 | 0,017353994 |
| Gm5148 | -3,48779589 | 2,63E-06 | 0,00145893 |
| Gm15920 | -2,374550636 | 0,000294826 | 0,034093974 |
| C1ql3 | -1,85560514 | 1,98E-08 | 5,95E-05 |
| Col19a1 | -1,838063844 | 0,000422641 | 0,044757355 |
| 4930426D05Rik | -1,707124574 | 0,000260542 | 0,032846721 |
| Hba-a1 | -1,620855064 | 0,000200783 | 0,026228856 |
| Hbb-bs | -1,439857138 | 3,92E-09 | 2,23E-05 |
| Gm26917 | -1,386633341 | 2,76E-05 | 0,014065223 |
| Adamts3 | -1,126167841 | 4,49E-05 | 0,011666088 |
| Gm23935 | -0,998720122 | 1,67E-06 | 0,001586252 |
| Gm42418 | -0,944061077 | 0,000101979 | 0,037259006 |
| Aldh1a1 | -0,912554686 | 0,000201205 | 0,026851195 |
| C1qtnf4 | -0,736523276 | 5,65E-05 | 0,022206472 |
| Sphkap | -0,647113152 | 0,000133835 | 0,044757355 |

**Table S1. DEGs in the Nac of 8 weeks-old males**
**Likelihood Ratio Analyses for synergy between R6/1 and *Rarβ***

**Up regulated**

| Gene Name | Log2 Fold Change | p-value | adjusted p-value |
|---|---|---|---|
| Ndnf | 0,809498377 | 5,91E-05 | 0,022873384 |
| Gm13111 | 0,968022398 | 0,000427063 | 0,044777892 |
| Igfbp5 | 1,022452993 | 8,28E-05 | 0,026851195 |
| Zic1 | 1,022750974 | 0,000142106 | 0,047197702 |
| Arx | 1,072504626 | 6,35E-05 | 0,026228856 |
| Prdm16 | 1,092355243 | 6,67E-05 | 0,014065223 |
| Epb41l5 | 1,141653007 | 0,000174155 | 0,025500948 |
| Hmgcs2 | 1,293639092 | 0,000235876 | 0,033696427 |
| Insm1 | 1,303711462 | 0,00015572 | 0,025902468 |
| Nrep | 1,307948645 | 4,57E-06 | 0,004748225 |
| Btg2 | 1,326937995 | 0,000113269 | 0,018658772 |
| Tead1 | 1,375326517 | 6,70E-07 | 0,001069265 |
| Prelp | 1,391092803 | 8,10E-05 | 0,015568163 |
| Sox4 | 1,394460472 | 8,78E-07 | 0,001207107 |
| Fzd5 | 1,407503938 | 0,000223517 | 0,028737588 |
| Zmynd10 | 1,42357367 | 9,30E-05 | 0,015754933 |
| Sox11 | 1,439713145 | 1,40E-05 | 0,011666088 |
| Nt5dc2 | 1,497887598 | 3,46E-05 | 0,009398067 |
| Rnd3 | 1,510454026 | 0,000141457 | 0,038018908 |
| Ccnd2 | 1,519843636 | 5,66E-08 | 0,000171125 |
| Tpx2 | 1,526575029 | 0,000507303 | 0,047855348 |
| St8sia2 | 1,582095757 | 8,32E-05 | 0,015568163 |
| Cd24a | 1,74534894 | 8,91E-07 | 0,001460814 |
| Fezf2 | 1,761311406 | 5,49E-05 | 0,014065223 |
| Dlx2 | 1,768076781 | 3,11E-06 | 0,0014788 |
| Shisa2 | 1,803777967 | 6,13E-05 | 0,014065223 |
| Pipox | 1,811757245 | 0,000348734 | 0,037495627 |
| Lmnb1 | 1,823472221 | 5,09E-09 | 1,34E-05 |
| Mcm3 | 1,8862095 | 1,57E-05 | 0,004982264 |
| Mex3a | 1,930508017 | 1,15E-06 | 0,00080728 |
| Ascl1 | 1,936753971 | 0,00015426 | 0,023569867 |
| Sfrp1 | 1,981276127 | 4,08E-11 | 2,15E-07 |
| Ccna2 | 2,065815185 | 0,000201977 | 0,026228856 |
| Knstrn | 2,152267946 | 0,000191051 | 0,026228856 |
| Kif11 | 2,173290281 | 0,000159233 | 0,023558833 |
| Tcte1 | 2,289614258 | 0,000327621 | 0,037495627 |
| Dlx1 | 2,292402793 | 6,84E-14 | 2,27E-09 |
| Mapk15 | 2,31199383 | 0,00015039 | 0,023558833 |
| Stbd1 | 2,363720423 | 0,000230333 | 0,029043981 |
| Nuf2 | 2,38865631 | 3,90E-06 | 0,001586252 |
| Espl1 | 2,480439679 | 0,000474761 | 0,04732201 |
| Prr11 | 2,562473855 | 0,000158726 | 0,023674596 |
| Ak7 | 2,581610968 | 8,45E-05 | 0,015568163 |
| Dmkn | 2,628936636 | 0,000308065 | 0,035004931 |
| CT571252.1 | 2,651109515 | 0,000166899 | 0,023674596 |
| Kif15 | 2,676466019 | 0,00028909 | 0,033819329 |
| Foxj1 | 2,690844699 | 7,96E-08 | 8,34E-05 |
| Igfbpl1 | 2,731320446 | 0,000472371 | 0,04732201 |
| Hmgb2 | 2,754915348 | 2,69E-07 | 0,000218145 |
| Vwa5b1 | 2,764944994 | 0,000316616 | 0,03557224 |
| Foxm1 | 2,794538105 | 1,31E-09 | 4,40E-06 |
| Drc7 | 2,81092341 | 1,46E-05 | 0,004699321 |
| Uhrf1 | 2,872601721 | 3,07E-06 | 0,001460814 |
| Ttc21a | 2,88880034 | 0,000101348 | 0,017353994 |

| | | | |
|---|---|---|---|
| Kif20a | 2,909708584 | 0,00043626 | 0,044513117 |
| Bub1b | 2,919755323 | 0,000220717 | 0,028518657 |
| Melk | 2,920645562 | 0,000514722 | 0,049879979 |
| Lrrc23 | 3,010287255 | 3,64E-05 | 0,009623962 |
| Cdca2 | 3,081830006 | 2,44E-05 | 0,006773217 |
| Rrm2 | 3,131467482 | 5,12E-08 | 5,95E-05 |
| Ect2 | 3,153207973 | 8,36E-05 | 0,015568163 |
| AC159264.1 | 3,189797938 | 0,000385884 | 0,041635387 |
| Cdk1 | 3,237152211 | 8,98E-05 | 0,015568163 |
| Prc1 | 3,359816238 | 3,89E-06 | 0,001586252 |
| Ckap2l | 3,497889223 | 6,27E-07 | 0,00044816 |
| Cenpf | 3,531939485 | 2,23E-05 | 0,00659999 |
| Ppp1r36 | 3,604032865 | 0,000447925 | 0,046297534 |
| Ttk | 3,643391388 | 0,000521234 | 0,049009376 |
| Ccnb1 | 3,698523617 | 3,81E-05 | 0,009757962 |
| Pbk | 3,70050448 | 5,90E-05 | 0,014065223 |
| Dlgap5 | 3,809210938 | 0,000335474 | 0,037794101 |
| Gtse1 | 3,931680466 | 1,88E-06 | 0,00117279 |
| Ube2c | 3,999795631 | 3,33E-06 | 0,001524796 |
| Ndc80 | 4,017037361 | 0,000211042 | 0,027811984 |
| 3110053B16Rik | 4,022119397 | 7,67E-05 | 0,014833972 |
| Gm6682 | 4,138130651 | 1,73E-08 | 2,62E-05 |
| Mis18bp1 | 4,226471595 | 1,20E-05 | 0,004195129 |
| Pimreg | 4,255509256 | 5,20E-05 | 0,012219774 |
| Mki67 | 4,29683741 | 0,000372745 | 0,040934907 |
| Kif2c | 4,352264486 | 7,52E-05 | 0,014747285 |
| Lrrc74b | 4,394571711 | 8,88E-06 | 0,003192143 |
| Spc25 | 4,537979518 | 1,15E-05 | 0,003979346 |
| Esco2 | 4,682888747 | 5,25E-05 | 0,012219774 |
| Kif18b | 4,691977946 | 0,000110185 | 0,018521438 |
| Tk1 | 4,87005843 | 7,79E-06 | 0,003041496 |
| Lbp | 4,965223075 | 0,000307212 | 0,036148392 |
| Col8a2 | 5,091298901 | 6,96E-05 | 0,014065223 |
| Cfap52 | 5,183564209 | 8,27E-05 | 0,015568163 |
| Ankle1 | 5,328226882 | 1,04E-05 | 0,017353994 |
| Rec8 | 5,804774305 | 2,28E-05 | 0,006773217 |
| Cdhr4 | 8,167389657 | 8,68E-09 | 1,84E-05 |
| 2010001K21Rik | 10,26147403 | 6,42E-06 | 0,014065223 |
| Tmem212 | 10,74281485 | 2,76E-08 | 0,000216524 |
